# Mapping the spatial architecture of glioblastoma from core to edge delineates niche-specific tumor cell states and intercellular interactions

**DOI:** 10.1101/2025.04.04.647096

**Authors:** Saad M. Khan, Anthony Z. Wang, Rupen R. Desai, Gyeong Dae Kim, Colin R. McCornack, Rui Sun, Sonika M. Dahiya, Jennifer A. Foltz, Ngima D. Sherpa, Katherine Xu, Harrison Olszewski, Markus Anzaldua-Campos, Yuhui Song, Elizabeth Seider, Natan Pirete Gadens, Jose Maldonado, Lydia Leavitt, Timothy West, Alexander F. Wang, Bryan D. Choi, Eric C. Leuthardt, Bhuvic Patel, Kathleen Pfaff, Linda Nieman, Aleksandar Krbanjevic, Al Charest, Albert H. Kim, Gavin P. Dunn, Allegra A. Petti

## Abstract

Treatment resistance in glioblastoma (GBM) is largely driven by the extensive multi-level heterogeneity that typifies this disease. Despite significant progress toward elucidating GBM’s genomic and transcriptional heterogeneity, a critical knowledge gap remains in defining this heterogeneity at the spatial level. To address this, we employed spatial transcriptomics to map the architecture of the GBM ecosystem. This revealed tumor cell states that are jointly defined by gene expression and spatial localization, and multicellular niches whose composition varies along the tumor core-edge axis. Ligand-receptor interaction analysis uncovered a complex network of intercellular communication, including niche- and region-specific interactions. Finally, we found that *CD8 GZMK* T cells colocalize with *LYVE1 CD163* myeloid cells in vascular regions, suggesting a potential mechanism for immune evasion. These findings provide novel insights into the GBM tumor microenvironment, highlighting previously unrecognized patterns of spatial organization and intercellular interactions, and novel therapeutic avenues to disrupt tumor-promoting interactions and overcome immune resistance.

## INTRODUCTION

Glioblastoma (GBM), the most common primary malignant brain tumor in adults, continues to be a devastating disease, with a median survival of 15-20 months^1–3^. One of many factors impeding the development of new therapies is the multi-level heterogeneity of the GBM ecosystem, which has been extensively studied from the perspective of genomics^4–6^, transcriptomics^7,8^, and single-cell RNA-sequencing (scRNA-seq)^9–16^. In particular, exome and genome sequencing studies have catalogued extensive intratumoral genetic heterogeneity in GBM. More recently, scRNA-seq studies have established that GBMs contain cells in distinct transcriptionally-defined states, and have revealed an immune microenvironment characterized by sparse, dysfunctional cytotoxic T cells^17–21^ and enrichment for immunosuppressive myeloid cells^22^ that drive T cell exhaustion^23^.

Spatial heterogeneity in GBM is particularly multifaceted, ranging from microscopic histopathological hallmarks (e.g. regions of microvascular proliferation and necrotic foci surrounded by palisading cells) to larger spatial domains visible using MRI (necrotic core, viable tumor core, infiltrative margin). Recently, spatial transcriptomic (ST) studies have established several principles regarding GBM tissue organization. Hypoxia has emerged as a key spatial organizer of the tumor core, driving genomic instability, macrophage polarization, and a reactive-mesenchymal transcriptional program^24–27^. The immune landscape varies systematically with tumor cell phenotype, with mesenchymal tumor states consistently co-localizing with macrophages, and microglia predominating in glial-enriched regions^25,28,29^. And the tumor core and periphery are enriched for different scRNA-seq-derived tumor cell states^25,28,30–33^. Despite this progress, the multicellular niches that constitute the GBM microenvironment, the intercellular interactions that sustain them, and how both vary systematically along the core-to-edge axis have remained incompletely defined. Such variation may have profound clinical implications, given that most recurrences arise from the tumor margin^34^.

Here, we leveraged spatial transcriptomics, scRNA-seq, and multi-region tissue sampling to comprehensively define the spatial architecture of GBM along the core-edge axis (Fig. 1, Fig. S1). We first used the Visium platform to analyze 22 samples from nine IDH-wildtype GBM patients. By integrating these data with scRNA-seq from 11 GBM patients and a single-cell nonmalignant brain atlas,^35^ we derived a novel set of tumor cell states defined jointly by transcriptional profile and spatial localization, and identified spatially segregated multicellular niches that vary systematically from core to edge. Ligand-receptor analysis revealed a complex, spatially organized network of intercellular communication within and between these niches. We also identified colocalization of *GZMK+* T-cells and *LYVE1+CD163+* macrophages near the vasculature. We validated our findings using the Xenium platform (15 samples, 8 patients), published scRNA-seq data from 240 patients^36,37^, and for selected interactions, multiplexed immunofluorescence (mIF) or RNAscope. Together, these analyses provide a spatially resolved portrait of the GBM microenvironment across multiple resolutions and identify potentially targetable intercellular interactions underlying its organization.

**Figure 1:**
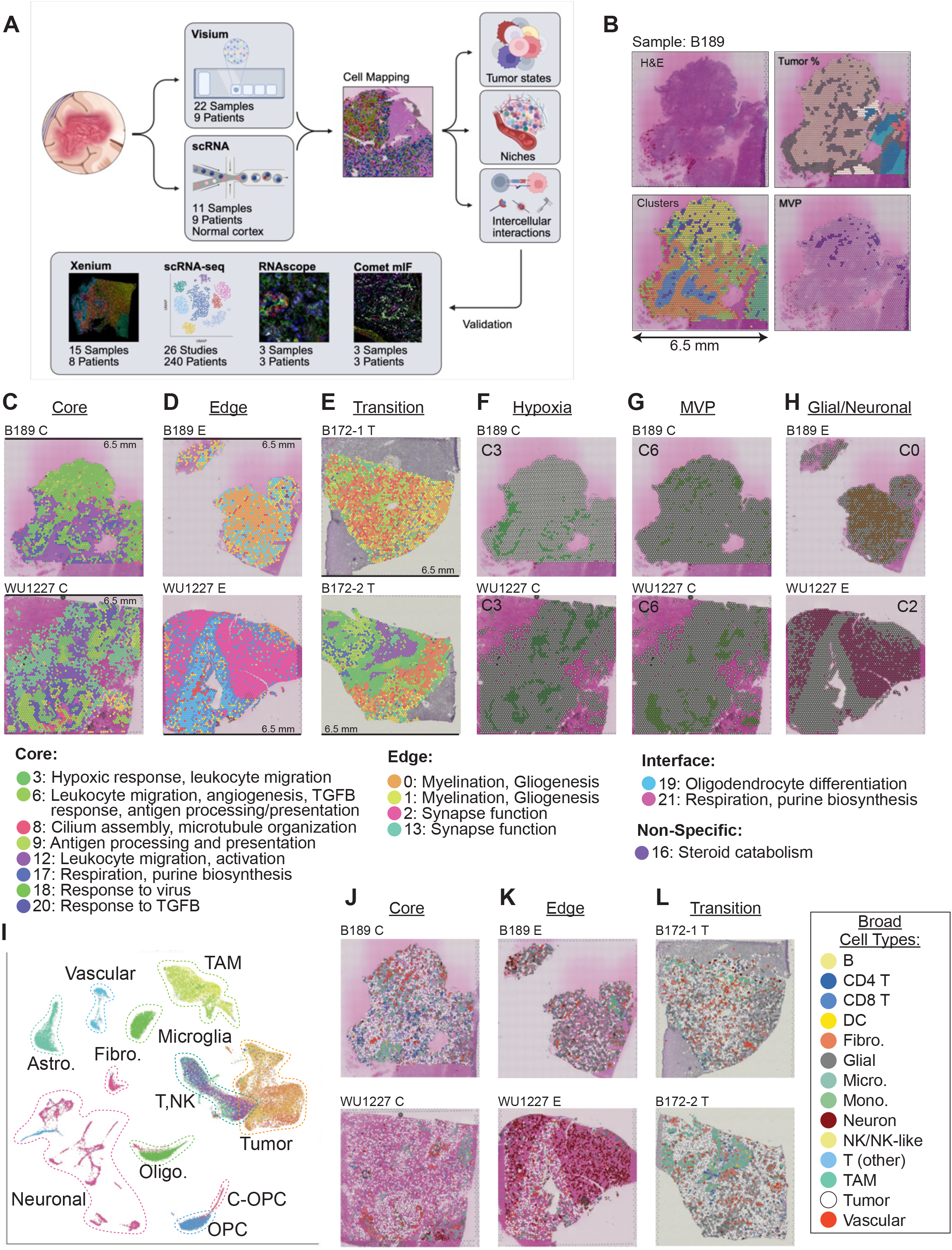
Overview of data, clustering, and cell mapping. **(A)** Summary of samples, patients, data types, analyses; further details in Table S1. **(B)** Clockwise from top left: H&E image for core tumor sample from B189, neuropathologist’s tumor content estimates, neuropathologist’s identification of microvascular proliferation (MVP), and unsupervised graph-based clusters depicting MVP and additional variation. **(C)** Louvain gene expression clusters in representative core samples, with cluster IDs and Gene Ontology enrichments as indicated below. **(D)** Clusters in representative edge samples. **(E)** Clusters in representative transitional 5ALA+ samples. **(F)** Hypoxia-response cluster, C3, in core samples. **(G)** MVP cluster, C6, in core samples and transitional 5ALA+ samples. **(H)** Glial (C0, C1) and neuronal (C2) clusters in edge samples. **(I)** UMAP projection of scRNA-seq data used as a reference for cell mapping. **(J)** Cell mapping results for representative core tumor sections showing dense tumor cellularity, vascularization, and pockets of immune infiltration, with cell type legend at right. **(K)** Cell mapping results for representative edge sections showing minimal tumor content and abundant neurons or glia. **(L)** Cell mapping results for representative transitional 5-ALA+ sections showing dense tumor cellularity, vascularization, and pockets of immune infiltration.

## RESULTS

### Identification of tumor core, edge, and core-to-edge transition zones

We obtained 22 samples along the core-to-edge axis from nine patients undergoing standard-of-care surgery for primary GBM. We employed pre-operative MRI and/or intraoperative 5-aminolevulinic acid (5-ALA) fluorescence to assess the relative position of each sample along the core-to-edge axis^38^ (Fig. S2, Table S1, Methods). Using this information, samples were designated as originating from the tumor *core* (*n*=7), tumor *edge* (*n*=6), or core-to-edge *transition* zone (*n*=9). Each sample was preserved in OCT or FFPE, stained with H&E, and analyzed using the Visium spatial transcriptomic platform. Six of these samples were also analyzed using Xenium *in situ* transcriptomics (Fig. 1A, Fig. S3-S4, Table S1-S2).

We then used two independent approaches to complement the core and edge designations established intraoperatively. First, we incorporated histopathologic tumor core and edge designations made by a board-certified neuropathologist (S.D.). These were consistent with designations based on 5-ALA fluorescence, MRI, and surgical evaluation, but indicated that most samples - even those from the tumor edge - contained some regions of high tumor cell density. Within each sample, manual histologic annotation indicated regions of high tumor cell density (consistent with tumor “core”), low tumor cell density (infiltrative tumor “edge”). When possible, annotation was also provided for microvascular proliferation (MVP), palisading necrosis, predominantly non-malignant white matter, and predominantly non-malignant gray matter (Fig. 1B, Fig. S5). Second, InferCNV confirmed that low-malignancy spots predominantly localized to edge-rich regions, and high-malignancy spots to core-rich regions (Fig. S6, Methods).

### Spot-level analysis of Visium data indicates that the tumor edge and core are transcriptionally distinct

Graph-based clustering of the spot-level data (Methods) revealed features that were apparent in the H&E images, such as microvascular proliferation, and captured extensive transcriptional heterogeneity that was not otherwise evident in the images. Clustering was performed at multiple resolutions, capturing low-resolution features shared across samples (resolution = 0.1: Fig. S7 A-C, Table S4), and high-resolution variation, including features enriched in the tumor core or edge (resolution = 0.9: Fig. 1C,D,F-H, Fig. S8, Table S4). Tissue slices resected from the transition zone contained a mixture of clusters but typically resembled the core more closely than the edge at a transcriptional level (Fig. 1E). Each cluster was classified as edge-associated, core-associated, interface-associated (between core and edge), or spatially non-specific (“N.S.”) based on its predominant histologic location as defined by pathology. Core-associated clusters were enriched for transcripts associated with hypoxia, microvascular proliferation, inflammation, myeloid cell activation, and related processes (Fig. 1F-G), while edge-associated clusters were enriched for myelination and synaptic signaling genes (Fig. 1H, Fig. S8C, Table S5), reflecting large-scale spatial differences in cellular composition between the tumor core and edge.

### Single-cell mapping reveals distinct cell type composition at the tumor core and edge

To analyze Visium data at single-cell resolution, the cellular composition of each spot was inferred using a random forest-based algorithm called CellTrek,^39^ which maps cells from a single-cell reference to x and y coordinates within the Visium data and enables downstream analyses to be performed at the level of discrete mapped cells. We created a single-cell reference by generating scRNA-seq data from 11 GBM samples, including two previously published samples^40^, and combining it with published data^35^ from non-malignant brain cell types that are often under-represented in scRNA-seq data, including neurons, glia, endothelial cells, and immune cells (Methods, Fig. 1I, Fig. S1A). The resulting cell mappings revealed distinct regions enriched for tumor, immune, endothelial, neuronal, and glial cells. Core samples were rich with tumor cells and infiltrating immune cells, particularly macrophages, which often resided in distinct islands (Fig. 1J). In contrast, edge samples contained abundant neurons and glia, as expected (Fig. 1K), and transitional samples displayed a mixture of cell types but were generally more core-like (Fig. 1L). Thus, the cell mapping was consistent with both histopathological annotations and functional enrichment of spot clusters. Spot composition was also inferred using Cell2Location^41^, which yielded comparable predictions (but lacked single-cell resolution).

### GBM tumor cell states are defined by both gene expression and spatial localization

Several studies have employed elegant computational methods to classify GBM cells into distinct transcriptional cell states based on scRNA-seq data^9,12–14,42^. Each study has yielded a different taxonomy of cell states, which are regularly updated to reflect data from new samples and technologies^9,10,16^. ST offers potential benefits over scRNA-seq for cell state inference because it does not introduce dissociation-associated artifacts, captures a more representative set of cells, and is more likely to reflect the true state of cells that are programmed by their spatial context^9^. We therefore asked whether the spatial organization of the mapped tumor cells could inform our understanding of tumor cell states. To this end, we analyzed the Visium data at single-cell resolution by clustering the mapped tumor cells in each sample using a spatially-weighted consensus clustering approach (Methods, Fig. S1A). We found that similar spatially-segregated tumor cell clusters recurred across samples with only subtle differences. Therefore, we used consensus clustering and hierarchical clustering to derive five “consensus” clusters, or “spatial tumor cell states,” shared across samples (Fig. 2A, Methods).

**Figure 2:**
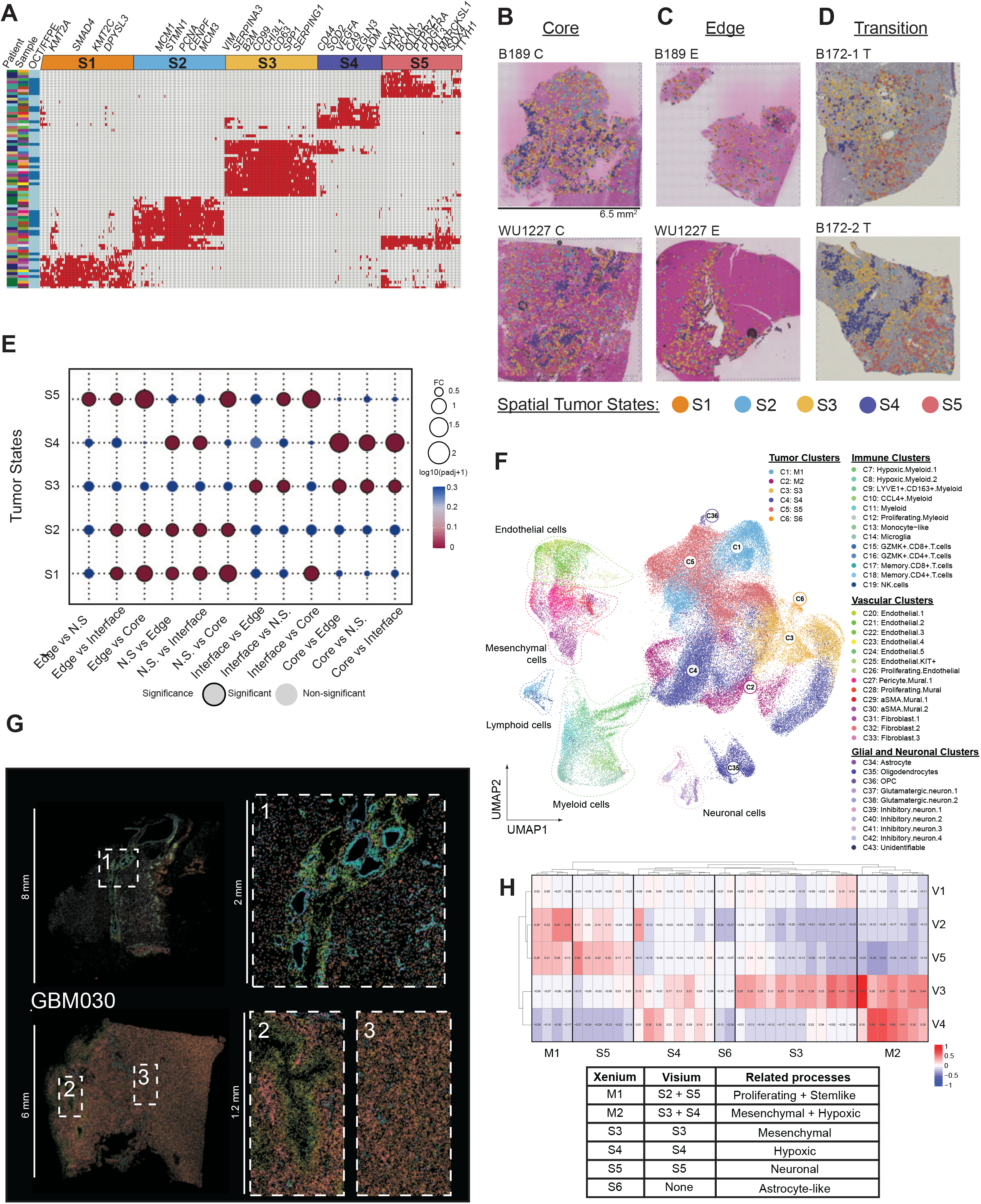
Derivation of spatial tumor cell states. **(A)** Consensus clustering of marker genes for spatial tumor cell states derived from mapped tumor cells. **(B)** Spatial tumor cell states in core tumor slices showing abundant Hypoxic (S4) and Mesenchymal (S3) tumor cells. **(C)** Spatial tumor states in edge samples, showing enrichment for Proliferating (S2) and Stem-like (S5) states in regions of high neuronal/glial content. **(D)** Spatial tumor cell states in transitional 5-ALA+ tumor slices showing a gradient from core Mesenchymal/Hypoxic to edge (Proliferative/Stem-like) states. **(E)** Fisher exact test for enrichment of Tumor states between domains. Dot color represents the Bonferroni-corrected p-value of enrichments and dot-size is the ratio of tumor state proportion (relative to total tumor cells) between domains. **(F)** UMAP visualization of all cells from primary Xenium cohort samples. **(G)** Xenium data from GBM026I and GBM030 with cell types colored as in panel F. **(H)** Hierarchical clustering of tumor clusters from primary and secondary Xenium cohorts based upon expression of Visium-based tumor signatures.

Each tumor state was enriched for specific biological processes: (1) Chromatin modification, especially via histone methylation (e.g. *KMT2A*, *KMT2C*, *NR2F1*, *SPAG9*); (2) Cell cycle and proliferation (e.g. *STMN1*, *CDKN2C*, *TUBA1B*); (3) Apoptosis, wound-healing, antigen presentation, and mesenchymal cell state (e.g. *VIM*, *CD99*, *CHI3L1*, *CD63*, *SPP1*, *ANXA2*, *HLA\**); (4) Response to hypoxia (e.g. *CD44*, *SOD2*, *VEGFA*, *MIF, ADM, CA9, EGLN3*, *PGK1*); and (5) Regulation of gliogenesis and neuronal stem cell differentiation (e.g. *SOX4/6/8/11*, *PTPRZ1*, *NRXN1*, *VCAN*, *BCAN*, *PDGFRA*, *OLIG2*, *DLL3*) (Table S6). We refer to these states as Histone Methylation (S1), Proliferating (S2), Mesenchymal (S3), Hypoxic (S4), and Stem-like (S5). Next, each tumor cell mapped by CellTrek was assigned to one of the five states S1-S5 based on marker gene expression, revealing spatially segregated mesenchymal regions, hypoxic regions, and mixed regions containing stem-like and proliferating tumor cells (Fig. 2B-D, Methods). This revealed that the tumor core, edge, and transition zone each display a characteristic, unique tumor cell composition (Fig. 2E), which we quantified further below. These states were also robustly recovered in the complete scRNA-seq dataset, where they exhibit high state-specificity, suggesting that the inclusion of spatial information helped clarify cell state definitions compared to single-cell data alone (Fig. S9B).

Although derived using independent, spatially-informed approaches, this cell state classification combines elements of previously published GBM cell state taxonomies, particularly the Neftel states^9,12,13,16,25^, including mesenchymal-like, stem-like, neuronal, proliferating, and hypoxic states (Fig. S9, Fig. S10). (The exception is our S1/Histone methylation state, which was the rarest in our data. This state most closely resembles the Greenwald^25^ “Chromatin” state, but does not significantly overlap any published cell state.) Critically, the Neftel GBM cell states are highly plastic *ex vivo*^9^. The fact that similar states can be derived based largely on spatial localization suggests that the microenvironment — through intercellular interactions — contributes to the specification of tumor cell states in conjunction with cell-intrinsic programming. *Ex vivo* plasticity may thus emerge partly from the loss of these contextual signals. This reasoning led us to systematically characterize intercellular interactions (below).

### Validation of spatial tumor cell states at single-cell resolution

To validate and extend these results at single-cell resolution, we employed Xenium *in situ* transcriptomics, a panel-based technology that quantifies transcript abundance at high resolution (<30 nm XY precision and <100 nm Z localization precision). We extended the pre-designed Brain/GBM Xenium panel (10X Genomics) with custom oligonucleotide probes, ultimately targeting 362 cell markers (Table S7). We used this platform to generate high-quality data for 15 primary GBM tumor sections (Methods, Tables S1,S8). Samples spanned three cohorts (of five, four, and six samples), with the third cohort comprising sections from matched Visium blocks.

Cell typing was performed on the primary cohort, and cell types were superimposed on the other samples using transfer learning (Methods). Our custom probe panel enabled us to annotate numerous key cell populations (Fig. 2F-G, Fig. S11, Table S9-S10). We identified several myeloid populations that were consistent with macrophages, but due to the limited number of probes, we conservatively referred to these as “myeloid” cells rather than “macrophages.” Two of these myeloid populations (C7, C13) expressed low levels of MHC class II and were consistent with E-MDSC and M-MDSCs, respectively^43^. Consistent with our previously published data, we identified *CD8^+^* and *CD4^+^ GZMK*^+^ T cells expressing low levels of *IL7R* and *GNLY*^21^. Xenium and Visium/CellTrek cell type assignments were highly concordant across the six matched tissue blocks (Pearson correlation = 0.76-0.99).

Next, we verified the presence of the Visium spatial tumor cell states in the Xenium data. Briefly, tumor cells were scored for the Visium-based cell state signatures, then hierarchically clustered (Fig. 2H). All Visium-derived tumor cell states were recapitulated in the Xenium data except for the Methylation state (S1), which was the rarest cell state in both the Visium data and scRNA-seq data. We also detected a Xenium-specific “Astrocytic” tumor cell state (S6) marked by high *RYR3* expression, and two “meta states,” M1 and M2, which were defined by combinations of Visium-based signatures: M1 represented a combination of the Proliferative (S2) and Stem-like (S5) tumor cell states, and M2 a combination of the Mesenchymal (S3) and Hypoxic (S4) tumor cell states, which we interpreted as a “Hypoxic Mesenchymal” cell state and has been previously reported^9^. Thus, tumor cell states derived from deconvolved Visium data were largely recapitulated by the single-cell resolution Xenium data.

### The tumor core and edge are composed of distinct multicellular niches with characteristic tumor cell content

We next sought to understand how tumor cell states co-localize with other cell types by defining cellular "niches," or spatially coherent multicellular communities. In the Visium data, we defined 23 niches by inferring the single-cell composition of each spot cluster via CellTrek (Methods, Fig. 3A). Classified according to the most-enriched cell type, these included tumor, immune, vascular, glial, and neuronal niches. Tumor-enriched niches were further differentiated according to the tumor cell states (and in some cases immune cells) for which they were enriched, with Mesenchymal and Hypoxic tumor cells often co-enriched for immune cells, particularly TAMs. These niches are highly correlated with – and add resolution to – the ten domains and four Cellular Communities recently identified in Lin et al^29^ (Fig. S13B). Quantitative comparison of our niches with the five Greenwald^25^ layers also indicated strong concordance (Fig. S13C). For example, the Greenwald Hypoxic layer correlated with our core-enriched niches, including the Hypoxic, Mes/TAM, Immune, T-cell, and Proliferative Tumor niches. In contrast, the Greenwald “Infiltrated Brain” layer primarily correlated with our Edge-associated neuronal and glial niches. The Greenwald Angiogenesis-Immune Hub correlated with our core-enriched microvascular proliferation niche.

**Figure 3:**
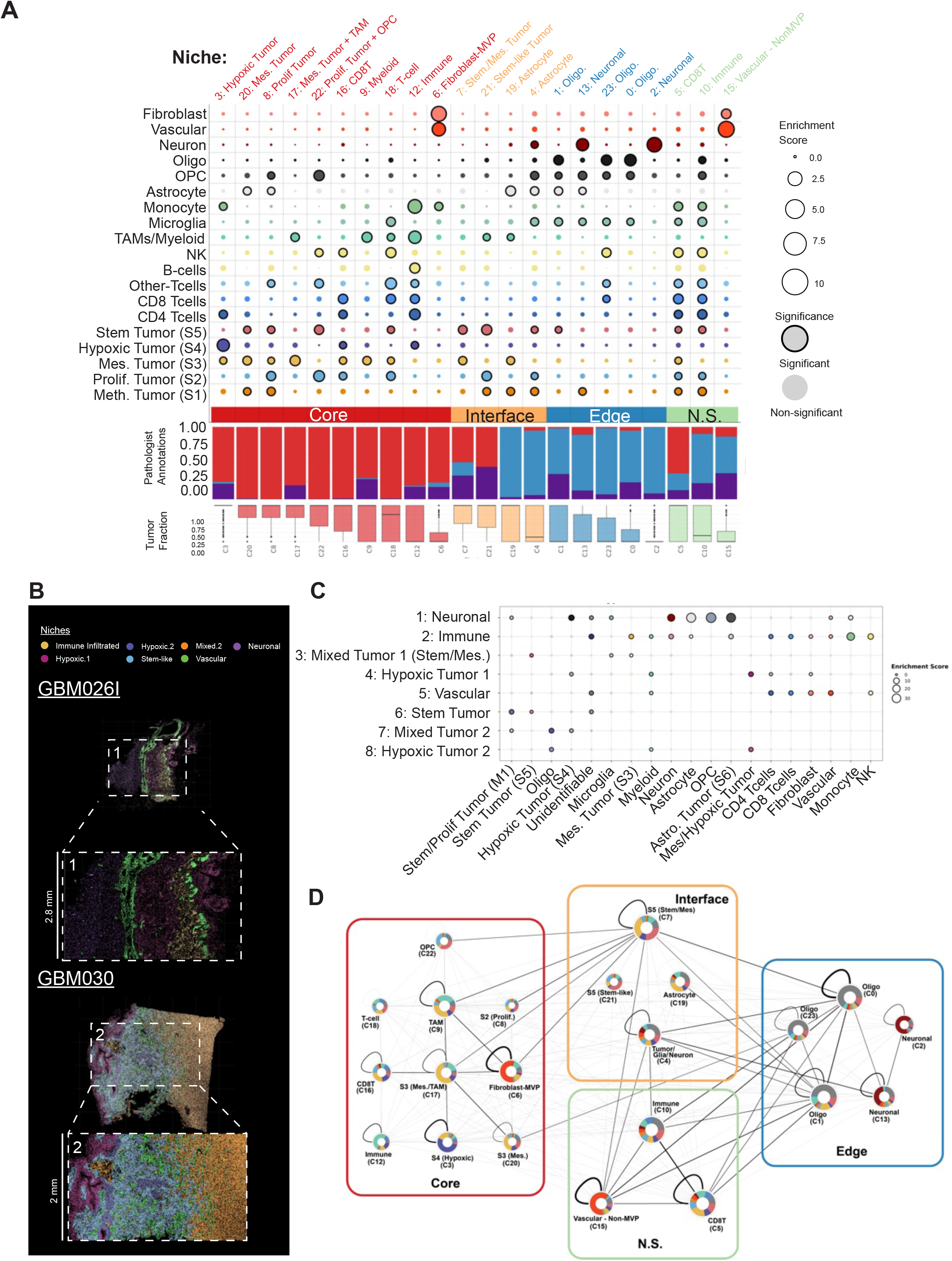
Tumor niches defined using Visium and Xenium. **(A)** Composition of each Visium cluster (columns, ordered by tumor location and tumor cell percentage) with respect to mapped cell types (rows). Edge/Core/Transition classifications were made by the surgeons and refined by the pathologist based on tumor cell density. Edge/Core/Interface/N.S. classifications were based on cluster distribution within the tissue. Circle size represents cell-type enrichment, and black borders represent statistically significant enrichment. Enrichment was calculated with respect to all cell types. The niche designation is indicated above each column based on the most-enriched cell type. **(B)** Xenium data from GBM026I and GBM030 with cells colored according to Banksy-defined niches. **(C)** Dot plot of enrichment scores for each cell type per respective Xenium niche. The enrichment score was calculated by dividing the cell type frequency per niche by the cell type frequency within the entire data set and signified by the size of each circle. Significance was calculated using the hypergeometric test and denoted by a black border around each respective circle. **(D)** Spatial proximity map of niches, where edge width is proportional to Weighted Mean Adjacency and edge color is proportional to the fraction of samples in which each niche pair is adjacent.

Niches were then independently defined in Xenium (Cohorts 1 and 2) using BANKSY^44^, using both the original Xenium cell types (Fig. S13D) *and* cell types harmonized with Visium (Fig. 3B-C, Table S11, Methods). This revealed eight distinct niches that corresponded to Visium niches, including Tumor, Immune, Vascular, and Neuronal/Glial niches. As with Visium, tumor-rich Xenium niches were enriched for distinct tumor states, including Stem-like/Proliferative tumor, Hypoxic tumor, Mesenchymal Tumor, and mixed Stem and Mesenchymal tumor. To compare Visium and Xenium niches quantitatively, we calculated pairwise correlation coefficients of cell type enrichment (Table S12, Fig. S13E, Methods). This indicated that the Xenium niches – despite the different platform and distinct patient cohort — were highly correlated with the Visium niches. However, the lower plexity of Xenium (and fewer samples) likely reduced our ability to resolve niches with the same granularity as Visium, resulting in fewer niches overall.

By ordering niches along the core-edge axis using histopathologic annotations and predicted tumor content, we found that each niche occupied a distinct position along the axis. In the tumor core, Visium revealed niches dominated by Hypoxic (S4) and Mesenchymal (S3) tumor cells, frequently co-enriched for TAMs, monocytes, T cells, and NK cells (Fig. 3A, Fig. S14A). The most immune-rich core niche (12) was co-enriched for Hypoxic tumor cells. Xenium independently confirmed this pattern (Fig. 3C). Although GBM is typically considered "immune-cold"^45^, these data suggest that T cells are present but sequestered within specific niches that may limit their function. In contrast to the tumor core, the tumor edge was composed of oligodendrocyte and neuronal niches enriched for Stem-like (S5) and Proliferating (S2) tumor cells (Fig. 3A). This was confirmed by our third Xenium cohort as well as matched core-edge scRNA-seq (Fig. S14C,D). Xenium further localized the astrocytic (S6) and Proliferating/Stem-like (M1) tumor cell states to the Glial/Neuronal niche.

Whereas Visium captured a single hypoxic tumor niche (3) in the core, Xenium resolved two distinct niches enriched for Hypoxic (S4) and/or Mesenchymal/Hypoxic (M2) tumor cells, but with differing immune and vascular compositions. Using the Xenium-specific cell annotations (Fig. S13B), both hypoxic niches were enriched for Hypoxic.Myeloid.1 cells consistent with E-MDSCs^43^, but differed in composition: Hypoxic 1 was additionally enriched for T cell populations and *KIT*⁺ endothelial cells, suggesting co-localization of T cells with immunosuppressive myeloid cells and angiogenic endothelial progenitors. Monocyte-like cells (M-MDSCs)^43^ were instead enriched in the Immune Infiltrated niche, highlighting distinct, niche-specific immunosuppressive programs.

Visium identified two distinct vascular niches with markedly different spatial profiles: one (6) was core-associated, co-localized with histologically identifiable regions of microvascular proliferation (MVP), and enriched for Mesenchymal (S3) tumor cells; the other (15) corresponded to dispersed spots distributed throughout the tissue and was not enriched for any particular tumor cell state (Fig. 3A). Xenium confirmed a discrete vascular niche populated by diverse endothelial and mural cell types, alongside *LYVE1*⁺*CD163*⁺ myeloid cells and *GZMK*⁺*CD8*⁺ T cells (Fig. S13B), explored further below.

Finally, we developed the Cluster Adjacency metric (Methods) to perform spatial proximity analysis of Visium niches. This indicated that Hypoxic, Mesenchymal, TAM, and MVP niches were mutual neighbors in the tumor core, while neuronal and glial niches clustered at the edge (Fig. 3E, Fig. S15). Analogous Xenium analysis confirmed co-localization of Tumor, Immune, and Vascular niches, as well as their spatial separation from Neuronal/Glial niches (Fig. S14B, Table S13).

### Ligand-receptor (LR) interactions mediate dense, redundant, bidirectional intercellular communication in the GBM microenvironment

This spatial organization — with highly ordered hypoxic regions in the core^24,25^ associated with mesenchymal and hypoxic tumor states, and stem-like and proneural states at the edge^25,28,30–33^ — recapitulates and extends prior observations, and implies that the microenvironment actively shapes tumor cell identity. We therefore systematically characterized ligand-receptor (LR) interactions, which mediate cell-cell communication, drive spatial patterning, and are of significant therapeutic interest.

To comprehensively characterize the LR interactions mediating intercellular communication in GBM, we used computational tools to predict LR interactions in four complementary datasets: our Visium spatial transcriptomics data, Xenium single-cell spatial transcriptomics data, in-house scRNA-seq reference dataset, and the GBMap scRNA-seq reference^36^ (Methods, Tables S14, S15). We integrated the LR predictions across these data sets to identify high-confidence interactions. LR interactions were included if detected in ≥2 datasets, with at least one spatial (Visium or Xenium) and one single-cell-resolution dataset (Xenium, scRNA-seq reference, or GBMap), and present in ≥3 samples/patients per technology. Cross-dataset integration used the harmonized cell typing scheme.

This yielded 438 unique cross-validated LR pairs mediating a total of 3,577 unique sender→receiver cell type interactions across 150 sender→receiver combinations among the cell types examined. Closer inspection of interactions involving tumor cells revealed a strikingly asymmetric communication landscape (Fig. 4A, Fig. S16A). Vascular, glial, and neuronal cells were the densest non-tumor senders (34–54 LR pairs per tumor state), while Lymphoid cells were the sparsest (1–10 LR pairs), consistent with GBM’s immunologically cold nature. Communication was predominantly bidirectional, with lymphocytes as notable outliers — most signaling was directed toward lymphocytes, consistent with tumor-driven immunosuppression. Communication between many cell-type pairs was highly redundant, driven largely by ECM biology (43 collagen/laminin pairs engaging *CD44*, *DAG1*, *SDC4*, *ITGB1*, or *SV2A*) and the Neurexin–Neuroligin group. Mesenchymal and Hypoxic tumor states sent the most outward signals, particularly through Pleiotrophin/Midkine (*MDK*/*PTN*) and Pro-Angiogenic families. Interestingly, Methylation and Stem-like tumor, which were enriched at the tumor edge, stood out for their very high Neurexin–Neuroligin output as senders (19 LR pairs), far exceeding other tumor states.

**Figure 4:**
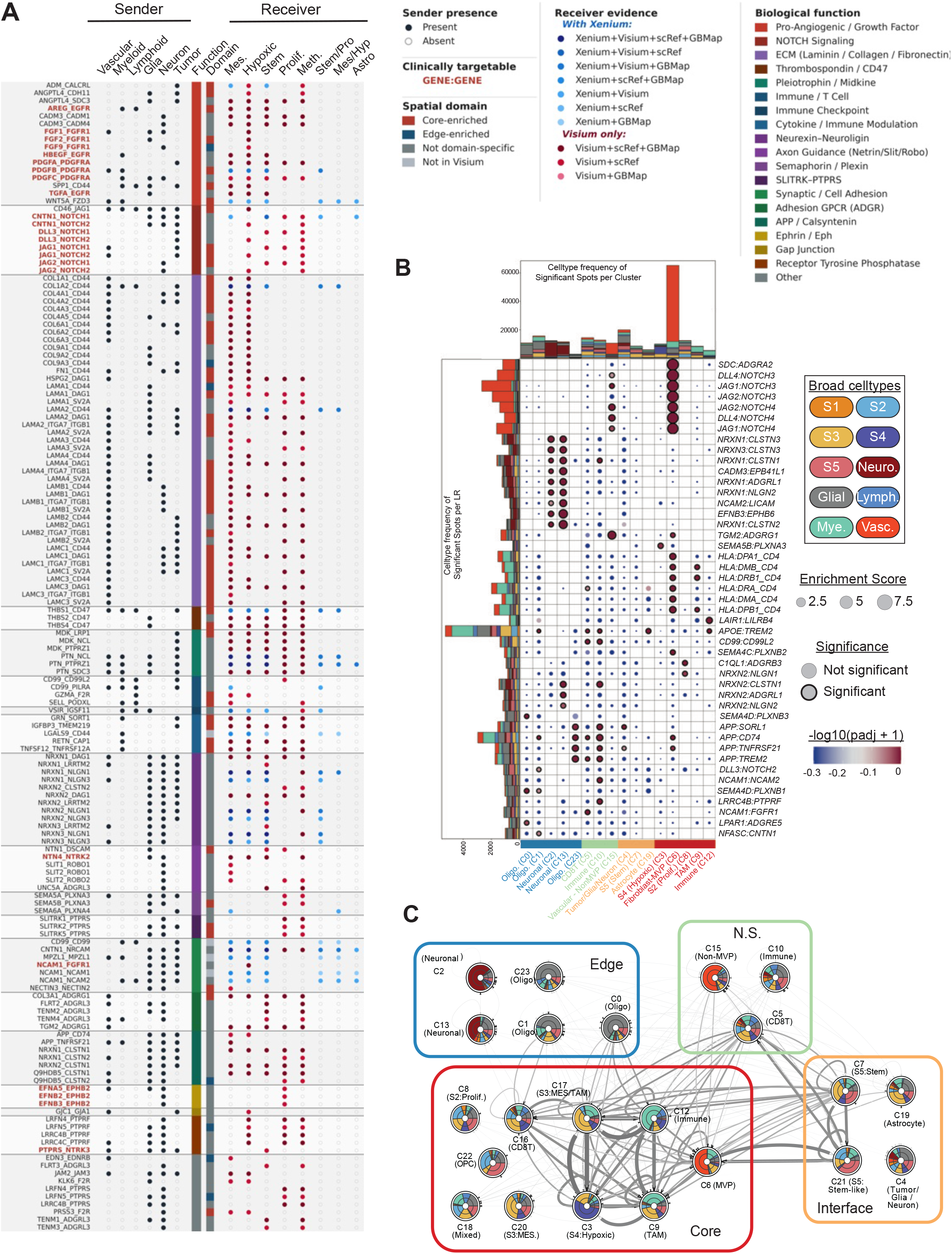
Ligand-Receptor (LR) analysis. (A) Dot plot showing high-confidence integrated ligand-receptor (LR) interactions from any cell type (sender, rows) to tumor cell states (receiver, columns). LR pairs are grouped by biological function (color bar, “Function”) and Visium domain enrichment (“Domain”). Each row represents a unique LR pair; bold red row labels indicate potentially targetable interactions (approved DGIdb^87^ antineoplastic agents targeting the receptor). Sender presence is indicated by filled (present) or open (absent) circles. Receiver dot color reflects the combination of platforms confirming the interaction: Interactions passing filter criteria: ≥2 independent evidence sources, ≥1 spatial platform (Visium or Xenium), ≥1 single-cell resolution platform (scRef, GBMap, or Xenium), with Xenium-only interactions excluded. **(B)** Contact-dependent LR interactions (rows) within each Visium niche (columns). Dot sizes indicate LR enrichment scores, and black dot borders indicate statistical significance (adjusted *p*-values). Row bar plots show cell type composition for the corresponding LR; column bar plots indicate cell type composition of the corresponding cluster. Clusters are ordered from edge-rich (left) to core-rich (right). **(C)** Inter-niche communication via diffusion-based LR’s predicted from Visium data. Edge width is proportional to the *LR_connectivity_*(*Cj, Cj*) score, indicating higher connectivity between core-rich clusters than edge-rich clusters. Circos plots on the network nodes indicate sender and receiver cell type frequencies.

### Intratumoral communication

The intratumoral LR landscape is also dominated by ECM signaling, driven largely by Mesenchymal and Hypoxic tumor states. Pleiotrophin (PTN) and Midkine (MDK), key growth factors in GBM, signal throughout the tumor, with virtually all states serving as both senders and receivers. Neurexin–Neuroligin signaling is abundant and asymmetric, with Stem-like and Methylation tumor states as the dominant senders, consistent with the enrichment of these states at the tumor edge. Consistent with recent work showing that intratumoral LR interactions contribute to the specification of tumor cell states^32^, we observed enrichment of *CD44* interactions in Mesenchymal/Hypoxic states and *NOTCH* interactions in Stem-like states (Fig. S16B-C), though most intratumoral LRs were shared across states.

### Mesenchymal tumor cells are a hub of intercellular communication

Mesenchymal (S3/M2) tumor cells consistently mediated the greatest number of unique cross-validated LR interactions with non-tumor cell types (Fig. 4A, Fig. S16B). Mesenchymal tumor received the most inbound LR pairs from every non-tumor sender (171 total incoming LR pairs summed across all non-tumor cell types) and sent the most outward (193 total outgoing LR pairs). These LRs were replete with ECM-receptor signaling and Pro-Angiogenic signaling. Within the tumor compartment, Mesenchymal tumor cells were again the most active senders and receivers across both platforms. Together, these results establish the Mesenchymal tumor cell state as the primary hub of intercellular signaling in GBM.

### Immunosuppressive signaling

Tumor states signaled to myeloid cells through 34 unique LR pairs and to lymphoid cells through 28 unique LR pairs, with Hypoxic tumor being the most active sender. Key interactions with established immunosuppressive functions included *CSF1*:*CSF1R*, an established driver of macrophage recruitment and polarization; *VSIR*:*IGSF11*, a targetable immune checkpoint in diffuse midline glioma^46^; *THBS1*:*CD47*, the well-characterized "don’t-eat-me" signal, present bidirectionally between multiple tumor states and both Myeloid and Lymphoid cells^47^; *SPP1*:*CD44*, secreted by Mesenchymal and Hypoxic tumor states to both Myeloid and Lymphoid cells, promoting immunosuppressive macrophage polarization and immune evasion^48^; *VEGFA*:*VEGFR1* from Hypoxic Tumor to Myeloid, Lymphoid, and Vascular cells, potentially suppressing the immune response and driving angiogenesis^49–51^; *MDK*:*LRP1*/*NCL* and *ANXA1*:*FPR*, likely driving immunosuppressive macrophage polarization^52–54^; *CD99*:*PILRA*, which inhibits myeloid cells^55^; and *APP*:*CD74*, potentially inhibiting phagocytosis of tumor cells^56^.

### Protumorigenic signaling

Signaling into the tumor compartment was again dominated by extracellular martrix (ECM) signaling, comprised of collagens and laminins signaling through integrin complexes (e.g. *ITGA7*-*ITGB1*), *DAG1*, *CD44*, and *SV2A*. Other notable pro-tumorigenic pathways included: Pro-Angiogenic and growth factor (e.g. *PDGF*, *FGF*, *VEGF*, *EGF* ligands) signaling to diverse tumor cells from vascular, glial, myeloid, and neuronal cells; Neurexin–Neuroligin signaling, which can promote GBM cell proliferation in response to neuronal activity^57,58^; NOTCH Signaling including *DLL3/JAG1/JAG2/CNTN1*:*NOTCH1/2*, which primarily impinged upon Hypoxic and Mesenchymal tumor states, consistent with the role of NOTCH in promoting stemlike behavior in GBM^59,60^; and signaling via the classical GBM mitogens Pleiotrophin (*PTN*) and Midkine (*MDK*)^61^.

### Vascular-tumor interactions are enriched for Mesenchymal tumor cells and reveal niche- and tumor cell state-specific signaling

Vascular cells were the most active non-tumor senders to tumor cells in the cross-validated analysis, with more unique LR pairs directed to Mesenchymal tumor cells than to any other tumor state. This selectivity is consistent with the spatial co-localization of Mesenchymal tumor cells with microvascular proliferation established in the Visium and Xenium niche analyses, and with the enrichment of pro-angiogenic LR interactions in the tumor core. The dominant pro-angiogenic signals in this data include *VEGFA:VEGFR1/VEGFR2*; *PDGFA/B/C/D:PDGFRA/B* (sent primarily from Mesenchymal tumor); and *ADM:CALCRL* (sent primarily by Hypoxic tumor). As detailed below, the tumor-vascular axis is notable as a context in which spatially restricted, tumor cell state-specific signaling can be identified; the two interactions described below illustrate this principle.

### LR interactions display spatial and niche specificity

To examine spatial variation in LR utilization, we first used the Visium data to assess contact-dependent LR interactions occurring *within* the same spatial niche. This revealed highly niche-specific communication patterns (Fig. 4B, Supplementary Data). For instance, the vascular niches (6 and 15) were strikingly enriched for the pro-angiogenic Notch signaling interactions, and each used distinct combinations of *DLL*, *JAG*, and *NOTCH* genes. Meanwhile, edge-associated neuron- and glia-rich niches (e.g. niches 2 and 13) were enriched for diverse *NRXN*-mediated interactions, which likely mediate pro-tumorigenic electrophysiological communication between neurons and GBM tumor cells^62^. The TAM-enriched core niche (9) was highly enriched for MHC Class II interactions, suggesting potential CD4 driven responses within the tumor core.

Next, to investigate LR interactions *between* different niches, we generated a high-level map of inter-niche communication involving secreted ligands. This analysis, which explicitly incorporated spatial information, reflected the spatial relationships among niches and again suggested that Mesenchymal (S3) tumor cells were a hub of intercellular communication (Fig. 4C, Methods, Supplementary Data). Core niches exhibited dense inter-communication, with extensive crosstalk within and between Mesenchymal tumor niches, TAM niches, Hypoxic tumor niches, and regions of MVP. Analogous analyses performed on Xenium data confirmed that Mesenchymal tumor is a signaling hub (Fig. S17, Fig. S18A-C). The majority of the LR interactions involved in this crosstalk included those known to drive angiogenesis in GBM, specifically *ANGPTL4-*, *SDC-*, *TNC-*, *THBS1-*, *ADM-*, *FGF-*, *VEGFA-*, and *VEGFC-*mediated interactions, *CXCL12*:*CXCL4*, and *WNT4*:*FZD*. Finally, we analyzed interactions within and among larger spatial domains (edge, core, and interface) which further supported the spatial specificity of intercellular communication (Fig. S19, Supplementary Data).

### *ADM*:*CALCRL* as a model for spatially and directionally specific vascular-tumor signaling

All of our LR analyses showed that Hypoxic (and Mesenchymal Hypoxic) tumor cells interact with Vascular cells via *ADM*:*CALCRL*. Although this interaction has been previously documented in GBM between hypoxic macrophages and vasculature^63^, it is not widely considered a canonical mediator of tumor:vascular signaling (cf. *VEGFA*:*VEGFR*, the target of the anti-angiogenic agent bevacizumab). We therefore chose to analyze *ADM*:*CALCRL* more closely using the Xenium data, which provided the requisite spatial and single-cell resolution. Xenium suggested that hypoxic myeloid cells *and* hypoxic tumor cells use *ADM:CALCRL* to communicate in a unidirectional manner with vascular cells, especially *KIT+* endothelial cells (Fig. 5A-B). This finding was consistent with the known role of hypoxia in inducing *ADM* expression via the transcription factor HIF1A. To understand how these interactions depended on the broader spatial context, we then investigated the *ADM:CALCRL* interaction at the *niche* level, and found that only one hypoxic niche (“Hypoxic 1”) interacted with the vascular niche via *ADM:CALCRL* across multiple samples (Fig. S18E). Importantly, niche-specific expression of *ADM* and *CALCRL* seemed to underlie the unidirectionality of this inter-niche interaction (Fig. 5C). For example, Hypoxic Mesenchymal (M2) tumor cells expressed *ADM* more highly when they were located in the Hypoxic 1 niche than in the vascular niche; similarly, endothelial.2 cells expressed *CALCRL* more highly when they were in the vascular niche than in the Hypoxic 1 niche. We observed similar region-specific expression of *ADM* and *CALCRL* in two orthogonal spatial data sets: our Visium data, and bulk RNA-sequencing data from the Ivy Glioblastoma Atlas Project (Ivy-GAP)^64^. In the Ivy-GAP data, *ADM* was higher in the perinecrotic zone and pseudopalisading cells, regions associated with hypoxia, while *CALCRL* was higher in hyperplastic blood vessels and microvascular proliferation (Fig. 5D). Finally, we used RNAscope *in situ* hybridization (ISH) to validate the existence and proximity of *ADM*+ tumor cells and *CALCRL*+ vasculature. Through this alternative modality, we found that across five samples approximately 20% of tumor cells (defined by *SOX2* expression) expressed *ADM* (Fig. 5E) while approximately 80% of vascular cells (defined by *PECAM1* expression) expressed *CALCRL* (Fig. 5F, Fig. S20). When calculating the median distance between *ADM+* tumor cells and vascular cells, we found *ADM+* tumor cells trended towards being closer to *CALCRL+* vascular cells compared to *CALCRL-* vascular cells (Fig. 5G). Regardless, on average, the average median distance across samples between *ADM+* tumor cells and *CALCRL+* vascular cells was less than 100 µm, well within the distance for sufficient paracrine signaling as previously demonstrated^65^. Together, the convergence of evidence across four datasets, spatial domain specificity, and ISH validation establishes *ADM:CALCRL* as a niche-specific, hypoxia-driven mediator of the tumor-vascular interface in GBM, and illustrates how spatially-restricted gene expression can give rise to directionally unique intercellular interactions.

**Figure 5:**
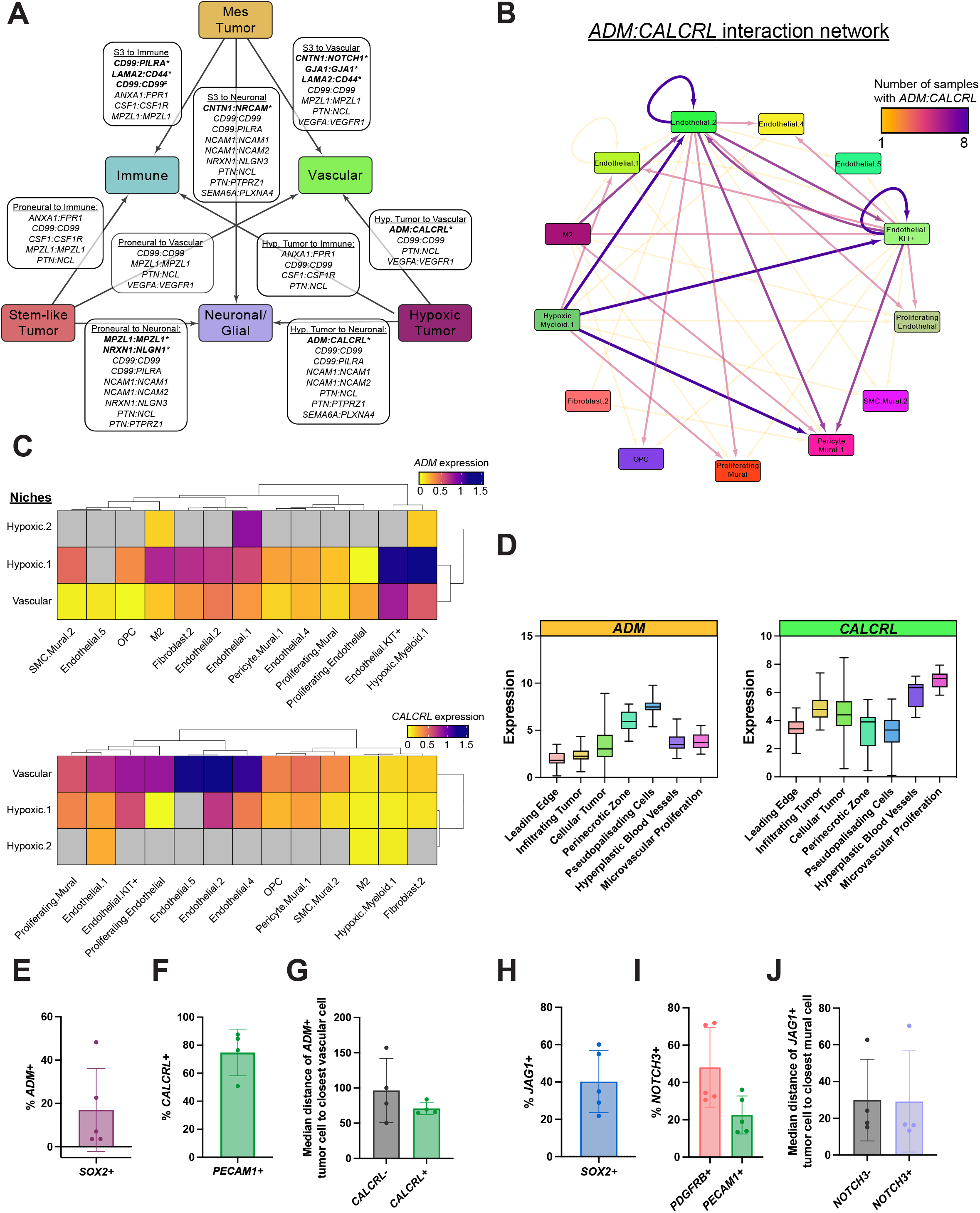
Tumor:Vascular interactions using Xenium and RNAscope. **(A)** Ligand-receptor interactions of tumor cells (Hypoxic: S2 or S4, Stem-like: S1 or S5) as senders and Myeloid, Neuronal/Glial, and Vascular cells as receivers. Ligand-receptor pairs unique to a given sender-receiver pair are italicized and labeled with an asterisk. **(B)** Interaction network of cell type sender-receiver pairs of the *ADM:CALCRL* ligand-receptor pair. The color and density of arrows were proportional to the number of samples the *ADM:CALCRL* interaction was detected as significant for a given sender-receiver pair. **(C)** Heatmap of *ADM* and *CALCRL* expression per cell type per niche. **(D)** Expression of *ADM* and *CALCRL* based upon localization in tumor region. Data derived from IVY-GAP. **(E)** Bar plots of % ADM^+^ of SOX2^+^ cells and **(F)** % CALCRL^+^ of PECAM1^+^ cells across all samples (n=5). **(G)** Bar plots showing median distance per sample between ADM^+^ SOX2^+^ tumor cells to closest CALCRL^-^ or CALCRL^+^ PECAM1^+^ cells. **(H)** Bar plots of % JAG1^+^ of SOX2^+^ cells and **(I)** % NOTCH3^+^ of PDGFRB^+^ or PECAM1^+^ cells across all samples. **(J)** Bar plots showing median distance per sample between JAG1^+^ SOX2^+^ tumor cells to closest NOTCH3^-^ or NOTCH3^+^ PDGFRB^+^ cells.

### Notch signaling at the hypoxic tumor-vascular interface

As noted above, redundant signaling between the vasculature and the tumor included additional non-canonical LRs. For instance, multiple Notch ligand-receptor pairs were consistently detected from Hypoxic tumor cells to vascular cells across three datasets (scRNA-seq, GBMap, and Visium) and exhibited Core-specificity: *JAG1:NOTCH1*, *JAG1:NOTCH2*, and *JAG1:NOTCH3*. In addition, *DLL3:NOTCH3* was detected from Stem-like and Proliferating tumor cells in two datasets. Because *JAG1*-mediated Notch activation at the tumor-vascular interface has been described in some cancer types^66^, we selected *JAG1:NOTCH3* for further analysis. The literature establishes *NOTCH3* as expressed predominantly by pericytes and vascular smooth muscle cells^67–69^, where it mediates mural cell differentiation and vessel stabilization. We again used RNAscope (with antibodies to JAG1 and NOTCH3) as a complementary method to support our Visium and Xenium findings. First, we observed that JAG1 was expressed by approximately 40% of *SOX2+* tumor cells across samples (Fig. 5H), and mural cells expressed higher levels (∼45%) of NOTCH3 than vascular cells (∼20%) (Fig. 5I). While no difference was found in median distance between JAG1*+ SOX2+* tumor cells and NOTCH3*-* or *NOTCH3+* mural cells, the average median centroid-to-centroid distances to both were within 20 um (i.e. within the range of direct cell contact) (Fig. 5J). These results show through an orthogonal method that JAG1 and NOTCH3 are robustly detected at the protein level, and distance measurements indicate co-localization at a distance compatible with ligand-receptor interaction.

### Heterotypic intercellular interactions shape the tumor transcriptome and tumor microenvironment (TME)

We next sought to systematically understand how signals from the TME integratively shape the tumor transcriptome. To this end, we applied NicheNet^70^ to predict which TME-expressed ligands regulate tumor gene expression, separately for each TME cell type, using the highest-confidence LR interactions from our integrated analysis (Methods, Fig. S21-S27, Table S16). Predictions were supported by ChIP-seq data identifying transcription factors linking LR interactions to downstream target genes. We integrated the results across all TME cell types to generate an overview of signaling from the TME to the tumor (Fig. 6), and from the tumor to the TME (Fig. S27).

**Figure 6:**
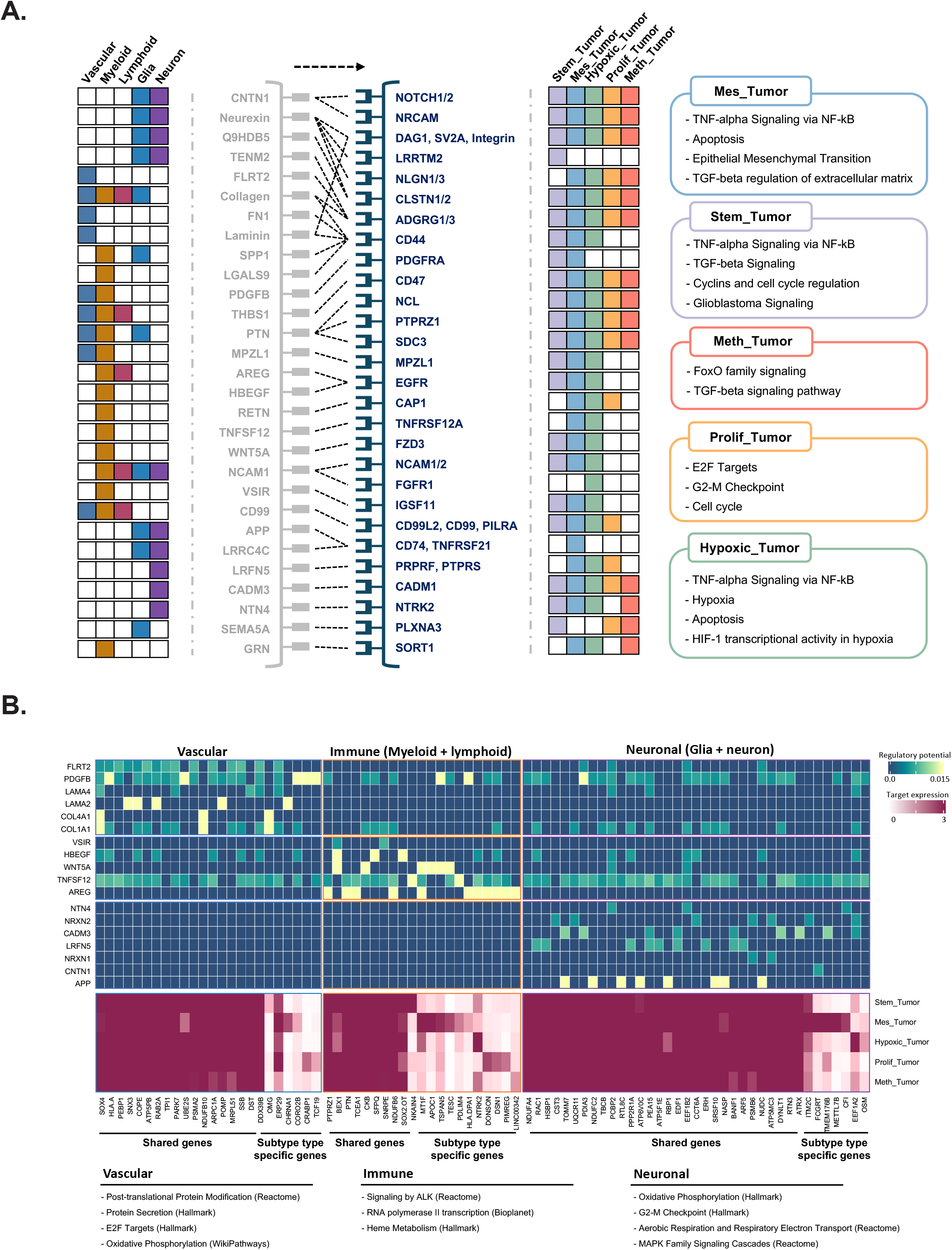
Interaction map from non-tumor cells to tumor cells. (A) Heatmaps illustrating interactions from non-tumor cells (including vascular, myeloid, lymphoid, and neuronal cells) to multiple tumor cell states, and biological pathways enriched among the predicted target genes associated with the corresponding interactions at each tumor cell state. Color indicates cell type. (B) Heatmaps illustrating the predicted regulatory potential of sender-specific (non-tumor cells) ligands and the expression of their predicted target genes in tumor cells. Predicted target genes are shown as tumor state-specific or shared across tumor states. Biological pathways enriched among ligand-regulated targets in each subtype are shown beneath.

The resulting TME→tumor gene regulation map highlights the LR interactions that are most likely to influence tumor cell biology (Fig. 6A). Interestingly, different TME senders converged on similar transcriptional programs in the tumor. These included pan-GBM programs canonically associated with tumorigenesis, as well as tumor cell-state-specific programs. For instance, multiple cell types induced Wnt-beta-catenin signaling and angiogenesis in Stem-like tumor cells, induced EMT and hypoxia-related signatures in Mesenchymal and Hypoxic tumor cells, and upregulated cell cycle-related genes in Proliferating tumor cells (Fig. S22-S26). Next, we sought to determine which target genes and pathways were regulated in a sender-specific manner (Fig. 6B, Methods). This revealed that vascular, immune, and neuronal senders regulate distinct target genes and pathway signatures within the tumor (e.g. *SOX4*, *PTN*/*PTPRZ1*, *ATRX* respectively (Fig. 6B).

The tumor→TME gene regulation map revealed distinct receiver-specific interaction patterns (Fig. S27). Tumor cells influenced myeloid cell gene expression via the immunosuppressive pairs *ANXA1*:*FPR1*, *SPP1*:*CD44*, *APP*:*CD74*, and *CSF1*:*CSF1R*; neuronal gene expression via Neurexins, *NCAM1*, *PTN*; vascular gene expression via *ADM*:*CALCRL, VEGFA/VEGFB:VEGFR1*, and Notch signaling. Sender (i.e. tumor state) specific interactions were also identified: *HLA-E* from Mesenchymal tumor regulated lymphoid gene expression, while *ADM*- and *ANGPTL4*-mediated interactions were characteristic of Hypoxic tumor. Notably, NicheNet predicted that *RAMP2* — required for ADM:CALCRL receptor complex formation — is co-regulated by *VEGFA* and *DLL3*, suggesting a coordinated multi-layered vascular circuit.

### Spatial sequestration of *GZMK*^+^ *CD8*^+^ T cells in vascular niches may contribute to the “immune-cold” phenotype

In recent work, we and others found that *GZMK*^+^ *CD8*^+^ T cells were the most abundant T cell population in several tumor types^21,71^, especially GBM, and were selectively clonally expanded in the tumor compared to matched peripheral blood. *GZMK^+^ CD8*^+^ T cells exhibited a highly non-uniform spatial distribution in our ST data, so we sought to further characterize the cellular environment surrounding *GZMK*^+^ *CD8*^+^ T cells, or *GZMK^+^ CD8^+^*T cell niches. To this end, we defined an enrichment score to identify cell types enriched within a given radius surrounding each *GZMK^+^ CD8*^+^ T cell (Fig. 7A, Methods). This showed that the most highly enriched cell types within a 0.5-20 µm radius of *GZMK^+^ CD8^+^* T cells included *CD8*^+^ T cells, CD4^+^ T cells, NK cells, and *LYVE1*^+^ *CD163*^+^ myeloid cells. The proximity of *LYVE1*^+^ *CD163*^+^ myeloid cells was striking, because these cells are thought to regulate migration of T cells into tumors, although the directionality of their influence is ambiguous^72–74^. In the homeostatic brain, *LYVE1^+^ CD163^+^* myeloid cells reside within the perivascular space^75–77^; given that T cells accumulate within the perivascular space in gliomas ^78,79^, we hypothesized that the colocalization of *LYVE1^+^ CD163^+^* myeloid cells and *GZMK^+^ CD8^+^* T cells may be specific to the vascular niche in GBM. To test this, we used several approaches to quantify the distance of each *GZMK*^+^ *CD8*^+^ cell to the nearest *LYVE1*^+^ *CD163*^+^ myeloid cell, and found that these cells were significantly closer to each other in the vascular niche compared to non-vascular niches (22.45 µm vs. 156.7 µm, p.value < 0.0001, Mann-Whitney U-test) (Fig. 7B-E, Methods). This relationship was not seen with *CD4*^+^ memory or *CD4*^+^ *GZMK*^+^ T cells across samples, and was not due to differences in the distribution of lymphoid cells and *LYVE1*^+^ *CD163*^+^ myeloid cells across the various niches (Fig. S26A-C). We then used an orthogonal data set to support the spatial relationship between *GZMK*^+^ *CD8*^+^ T cells and *LYVE1*^+^ *CD163*^+^ myeloid cells. In bulk RNA-seq data from IVY-GAP, *GZMK* and *LYVE1* expression were significantly correlated (R^2^=0.3418, p<0.001) in vascular regions only (Fig. 7F). Notably, *GZMB* was not correlated with *LYVE1* in either vascular or non-vascular regions (Fig. S26D).

**Figure 7:**
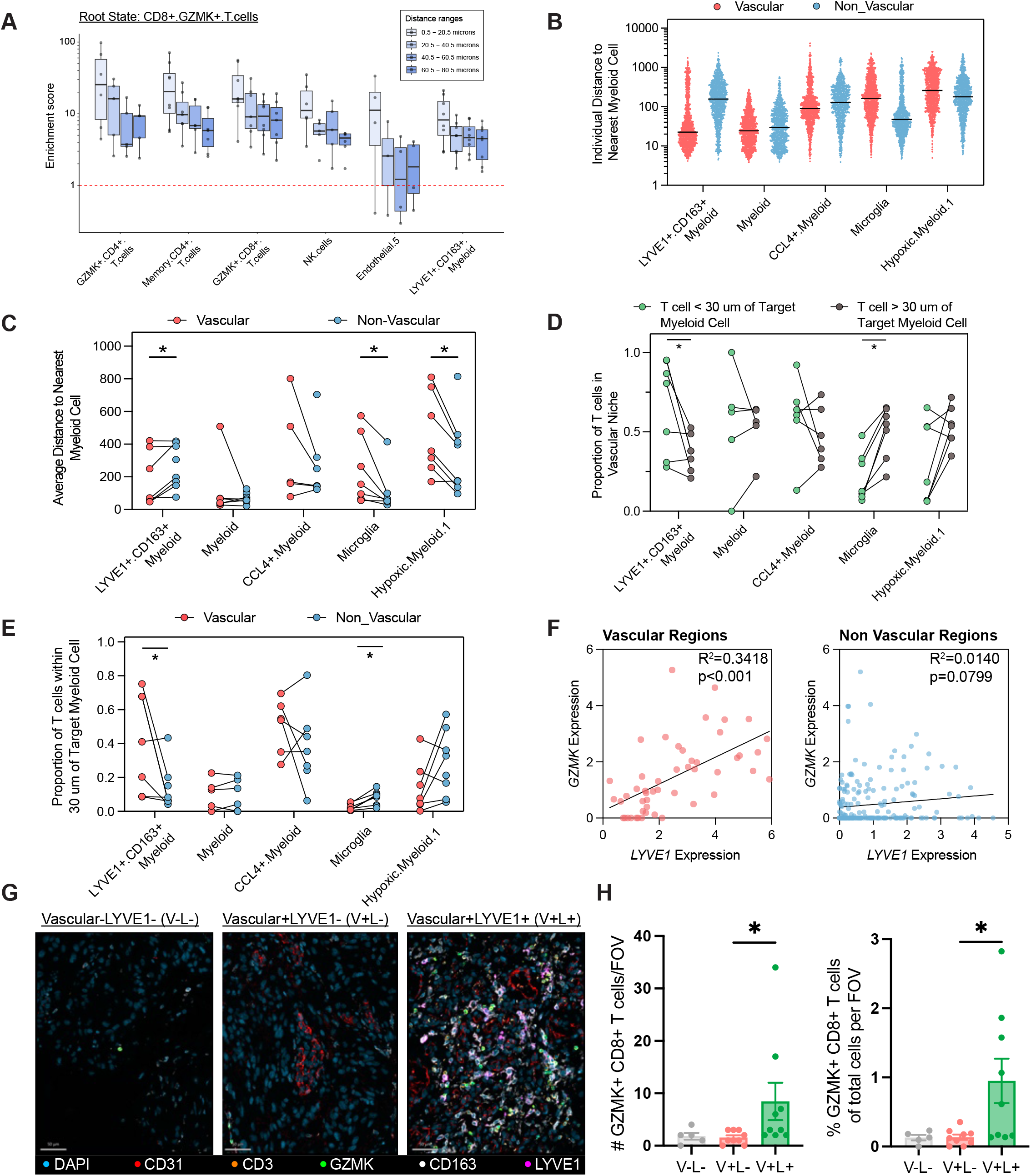
*CD8^+^GZMK^+^*T cells preferentially localize with *LYVE1^+^CD163^+^* myeloid cells in vascular niches. (A) Top 6 cell types enriched within 20 micron radius of *CD8^+^GZMK*^+^ T cells. Enrichment score was calculated by dividing the frequency of a target cell type within a given radius of a root state (*CD8^+^GZMK*^+^ T cells) by the frequency of the target cell type within the sample. Each dot represents a patient sample. (B) Distance to the nearest target myeloid cell of each *CD8^+^GZMK*^+^ T cell separated by localization within a vascular or non-vascular niche. Each dot represents a single *CD8^+^GZMK*^+^ T cell, with all *CD8^+^GZMK*^+^ T cells across all samples visualized. (C) Distances from Panel B averaged on a per sample basis. Each dot represents an individual sample. Significance was calculated using the Wilcoxon matched-pairs signed rank test. (D) Frequency of CD8^+^GZMK^+^ T cells in the vascular niche given proximity to the target myeloid cell (either within 30 μm or further than 30 μm). (E) Frequency of CD8^+^GZMK^+^ T cells within 30 μm of target myeloid cell given presence of T cell in vascular or non-vascular niche. (F) Representative images of V-L-, V+L-, and V+L+ regions captured via multi-plex immunofluorescence imaging with respective antibody and color pairs labeled. V-L-indicates non-vascular regions without LYVE1^+^CD163^+^ cells (# FOV = 5); V+L-indicates vascular regions without LYVE1^+^CD163^+^ cells (# FOV = 9); V+L+ indicates vascular regions with LYVE1^+^CD163^+^ cells (# FOV = 9). Total number of patient samples was 3. (G) Total count and relative frequency of CD8^+^GZMK^+^ T cells in respective regions. Statistical significance was calculated using Dunn’s multiple comparisons test. *indicates p <0.05 and ** indicates p-value < 0.01.

Surprisingly, *LYVE1*^+^ *CD163*^+^ myeloid cells were not detected in our scRNA-seq data set, nor in many other published scRNA data sets, although a recent publication highlighted *LYVE1* as a component of the “scavenger suppressive” profile in glioma myeloid cells^80^ (Fig. S26E). As a result, to validate the presence of these myeloid cells, we performed multiplexed immunofluorescence imaging on human GBM tumor sections and made several key observations. First, we found that LYVE1 and CD163 were present in human GBM tumors at the protein level (Fig. 7G) and co-expressed by the same cell (Fig. 7H). Furthermore, we categorized several FOVs according to the presence of (a) both LYVE1^+^ myeloid cells and vascular cells (V+L+), (b) only vascular cells (V+L-), or (c) absence of both (V-L-). Notably, we found that the number and overall frequency of CD8^+^GZMK^+^ T cells was significantly enriched only in V+L+ FOVs, supporting our hypothesis that CD8^+^GZMK^+^ T cells co-localize with LYVE1^+^CD163^+^ myeloid cells in specific vascular niches (Fig. 7H). Given the potentially immunosuppressive nature of *LYVE1*^+^ *CD163*^+^ myeloid cells in other tumor types, this interaction may present one possible mechanism by which T cells are sequestered within the perivascular space in the GBM TME.

## DISCUSSION

It is increasingly appreciated that solid tumors function as multicellular ecosystems in which the spatial organization and interactions among diverse malignant and non-malignant cells influence the development and behavior of the tumor, including its resistance to therapy. Here, using a combination of complementary spatial and single-cell transcriptomic technologies applied to multi-region tissue sampling along the tumor core-edge axis, we characterized the spatial organization of GBM and the ligand-receptor interactions that may govern intercellular communication across the tumor landscape. This revealed spatially specific tumor cell states, a core-to-edge gradient of multicellular niches, and spatially organized intercellular communication networks, with implications for the development of new therapies.

Notably, prior studies have identified spatial variation in tumor cell state, and are particularly unanimous on the organized clustering of hypoxic and mesenchymal cell states, the enrichment of neural-like tumor cells in regions of low tumor content, and the colocalization of mesenchymal tumor states with macrophages^24–26,28,29,32,81^. These concordant observations lend confidence to the broad spatial organization we describe, although precise niche definitions vary across studies. Critically, the intercellular communication networks sustaining niche organization have not been comprehensively characterized. Our work addresses this gap, providing a cross-platform, multi-resolution map of niche-specific ligand-receptor interactions and demonstrating that the core and edge are maintained by fundamentally different intercellular signaling programs.

Our first main finding is that GBM tumor cell states similar to published scRNA-seq-derived states can be derived *de novo* based on their spatial colocalization. Spatially-weighted consensus clustering of CellTrek-mapped tumor cells yielded five states — Histone Methylation (S1), Proliferating (S2), Mesenchymal (S3), Hypoxic (S4), and Stem-like (S5) — that are spatially coherent within samples and reproducible across patients. Xenium validation at single-cell resolution further resolved an Astrocytic tumor state (S6) and two composite tumor meta-states, M1 (Proliferative+Stem-like) and M2 (Mesenchymal+Hypoxic). Unlike states derived from scRNA-seq, their derivation through spatial colocalization offers new insights into tumor cell state specification. GBM cell states as measured by scRNA-seq are notoriously plastic, shifting dramatically upon tissue dissociation and culture *ex vivo*. That reproducible, spatially coherent states can be recovered using spatial approaches suggests that tumor cell state is partly specified by signals from the spatial microenvironment, rather than cell-intrinsic transcriptional programs alone, and that *ex vivo* plasticity arises in part because this spatial context has been eliminated. A recent study corroborated this perspective, demonstrating that intratumoral *CD44* and *NOTCH* signaling supports colocalization of specific tumor cell states^32^. This view motivated the subsequent questions we addressed: What is the multicellular context of each tumor cell state, and what molecular interactions contribute to niche specification?

Our second finding is that GBM is organized into spatially distinct multicellular niches whose composition varies systematically along the core-edge axis. Niche analysis using Visium and Xenium — two independent platforms applied to partially overlapping patient cohorts — yielded strikingly concordant results. Core niches were dominated by Mesenchymal (S3) and Hypoxic (S4) tumor cells, frequently co-enriched for TAMs, monocytes, T cells, DCs, and NK cells. Xenium further divided the hypoxic compartment into two distinct niches: Hypoxic Tumor 1, enriched for *KIT*⁺ endothelial cells (consistent with an angiogenic progenitor population) alongside multiple T cell populations and E-MDSC-like myeloid cells; and Hypoxic Tumor 2, with a distinct immune and vascular composition. This has important implications for immunotherapy: T cells within hypoxic niches are subject to compounding layers of immunosuppression — direct hypoxic dysfunction^82–84^, suppression by E-MDSCs^43^ and other myeloid cells^23,85,86^, and potential additional signals from the surrounding microenvironment. Immune checkpoint blockade alone may therefore be insufficient in these regions, and combination strategies targeting immunosuppressive myeloid cells or hypoxia itself may be required. In contrast, the tumor edge was dominated by oligodendrocyte and neuronal niches enriched for Stem-like (S5) and Proliferating (S2) tumor cells — states associated with invasiveness and tumor recurrence. The colocalization of these tumor cells with neurons and glia is consistent with prior evidence that tumor-neuron interactions drive tumorigenesis^58^, and suggests that therapeutic strategies at the tumor margin may need to target neurons and glia in addition to the tumor cells themselves.

Our third major finding is that intercellular communication in GBM is spatially organized, niche-specific, bidirectional, and highly redundant. Integrating LR interaction analyses across four complementary datasets — Visium ST, Xenium single-cell spatial, our in-house scRNA-seq reference, and the publicly available GBMap reference — we identified 438 cross-validated LR pairs mediating 3,577 unique sender→receiver interactions across 150 cell-type combinations. Several properties of this landscape stand out. First, interactions are strikingly asymmetric: Mesenchymal (S3/M2) tumor cells are the primary hub of intercellular communication, receiving the greatest number of inbound LR pairs from every non-tumor sender and deploying the most outbound interactions, which were enriched in ECM-receptor and pro-angiogenic pathways. Second, communication is largely bidirectional — with one notable exception: lymphoid cells receive far more tumor-directed signaling than they send in return, consistent with active, tumor-driven suppression of T cell function. Third, signaling is highly redundant: vascular cells alone utilize more than 50 unique cross-validated LR pairs directed toward Mesenchymal tumor cells. This redundancy has direct therapeutic implications — disrupting a single LR interaction is unlikely to meaningfully abrogate communication between a given cell-type pair, underscoring the need for combination approaches. According to the Drug/Gene Interaction Database^87^, many of the LR interactions we identified could potentially be targeted by at least one existing drug or compound.

The tumor-vascular axis is particularly illustrative of spatially specific LR signaling. *ADM:CALCRL* emerged as a model for niche-specific, hypoxia-driven vascular-tumor signaling: it was consistently detected across all four datasets, restricted to a single hypoxic niche (Hypoxic Tumor 1) in Xenium, confirmed by niche-specific expression of *ADM* and *CALCRL* in both Xenium and Visium, corroborated by region-specific expression in Ivy-GAP bulk RNA-seq, and validated by RNAscope ISH. NicheNet further predicted that *ADM* signaling drives *APOLD1* expression in vascular cells through the transcription factor *CREB1*, and that *RAMP2* (required for *ADM*:*CALCRL* receptor complex formation) is simultaneously regulated by *VEGFA* and *DLL3* — suggesting a coordinated, multi-layered vascular regulatory network. In parallel, *JAG1:NOTCH3* signaling from Hypoxic tumor cells to endothelial and mural cells was cross-validated across three datasets and confirmed by multiomic RNAscope experiments, establishing Notch signaling as an additional non-canonical mediator of the hypoxic tumor-vascular interface. Taken together, these findings offer a potential explanation for why bevacizumab, which targets only VEGF-A-mediated interactions, shows minimal efficacy in primary^88^ and recurrent^89^ GBM: the tumor-vascular interface is sustained by dozens of redundant, spatially organized LR interactions, of which VEGFA is only one.

Our fourth major finding concerns the spatial sequestration of *GZMK*⁺*CD8*⁺ T cells within the vascular niche and their proximity to *LYVE1*⁺*CD163*⁺ myeloid cells. *GZMK*⁺*CD8*⁺ T cells are the most abundant T cell population in GBM and are selectively clonally expanded within the tumor relative to matched peripheral blood. Despite their prevalence, they exhibit a highly non-uniform spatial distribution, and our data demonstrate that they are preferentially concentrated within the vascular niche, specifically near *LYVE1*⁺*CD163*⁺ myeloid cells. This spatial relationship was confirmed orthogonally in Ivy-GAP bulk RNA-seq (significant correlation of *GZMK* and *LYVE1* expression in vascular regions only, but not *GZMB*) and by multiplex immunofluorescence imaging, which showed enrichment of CD8^+^GZMK^+^ T cells only within vascular regions that contained LYVE1^+^CD163^+^ myeloid cells in additional GBM tissue. In the homeostatic brain, LYVE1⁺CD163⁺ myeloid cells reside within the perivascular space and regulate T cell trafficking^72,74^; their enrichment in the GBM vascular niche, near GZMK⁺CD8⁺ T cells, raises the possibility that they actively sequester T cells within the perivascular compartment and impede their infiltration into the tumor parenchyma. This would represent a spatially specific mechanism contributing to GBM’s "immune-cold" phenotype, and suggests that targeting LYVE1⁺CD163⁺ myeloid cells or the interactions that retain T cells in the perivascular space could improve the efficacy of immunotherapy. Studies in breast cancer have shown that ablation of LYVE1+ macrophages leads to significant infiltration of CD8^+^ T cells into the tumor core^72,74^.

Taken together, our data indicate that the tumor core and tumor edge constitute distinct ecosystems — each with characteristic tumor cell states, niche compositions, and intercellular communication networks — that are likely to respond differently to therapy. The tumor core is dominated by Hypoxic and Mesenchymal tumor states embedded in an immunosuppressive, angiogenic microenvironment, while the tumor edge harbors Stem-like and Proliferating tumor cells interacting with neurons and glia through synaptic adhesion pathways including Neurexin– Neuroligin signaling. These observations are consistent with two non-exclusive models of tumor development: Stem-like/Proneural tumor cells may be inherently more invasive and migrate to the infiltrating front, or tumor cells may be transcriptionally reprogrammed by their local microenvironment — receiving neuronal and glial signals at the edge, and immune, vascular, and hypoxic signals at the core. Our data on spatial variation in LR interactions and their transcriptional consequences lend support to the latter model. More speculatively, Stem-like/Proneural tumor cells may represent the founding tumor cell state from which Mesenchymal and Hypoxic states emerge as the niche matures — driven by increasing cell density, vascularization, and immune recruitment.

Most GBM recurrences arise from the infiltrative margin, yet most therapies are designed to target the contrast-enhancing tumor core. Our findings suggest that effective treatment will require strategies tailored to the distinct biology of each spatial compartment, including approaches capable of reaching and disrupting the interactions sustaining tumor cells at the edge. A key limitation of our study is that the LR interactions reported here were inferred computationally using databases of known interactions, and extensive experimental work will be required to validate and functionally characterize the most promising therapeutic candidates. Nevertheless, our integrated, multi-platform dataset provides a spatially comprehensive portrait of the GBM tumor microenvironment, and serves as a foundation for the development of spatially informed GBM therapeutics.

## METHODS

### Patient recruitment and sample collection

Adult patients undergoing neurosurgical intervention were screened for the following criteria: (1) age > 18 years and (2) likely presence of primary glioblastoma with clinical indications for surgical resection. Final diagnosis of primary IDH^WT^ GBM was later confirmed by histopathological analysis and molecular testing. Informed consent from patients was obtained prior to surgery following the IRB Protocol #201409046 (Washington University in St. Louis) or the Dana Farber/Harvard Cancer Center IRB Protocol #10-417 and Massachusetts General Hospital Secondary IRB Protocol #2022P001982. Patient characteristics were summarized in Table S1.

Patients underwent surgical management according to institutional standard of care, which includes preoperative administration of aminolevulinic acid (5-ALA, Gleolan). Initial samples were obtained using 5-ALA fluorescence, as evaluated by the surgeon, to define the tumor core. Subsequent regions were preoperatively selected based on MRI characteristics and sampled using intraoperative neuronavigation (Medtronic Stealth S8). For these MRI-guided samples, tumor core was defined with contrast-enhancement and tumor edge was defined as the peritumoral, non-enhancing, T2/FLAIR hyperintense tissue (representative image in Figure S2). Of note, samples were obtained immediately upon dural opening prior to tumor resection to limit brain shift and thereby maximize navigational accuracy. Resected samples were placed in normal saline and maintained on ice until further processing. All procedures and experiments were performed in accordance with the Helsinki Declaration.

### Visium data generation and processing

Using image-guided surgical biopsies and 5-aminolevulinic acid (5-ALA) fluorescence, we collected 22 samples from tumor regions ranging from the core to the margin in nine patients with IDH^WT^ GBM undergoing surgical resection of radiographic, primary high-grade glioma. Each section (i.e. sample) was preserved, stained with hematoxylin and eosin (H&E), imaged, and processed to generate a spatial transcriptomic library. The first 13 samples were fresh-frozen and OCT-embedded, and cDNA libraries were constructed according to the Visium Spatial 3’ Gene Expression User Guide Rev. D. The next nine samples were preserved in FFPE, and cDNA libraries were generated according to the Visium Spatial Gene Expression for FFPE User Guide Rev. E. All libraries were sequenced on an Illumina Novaseq S2 flow cell to a target depth of 50,000 reads/spot and yielded high quality transcriptome data (median values: 2300 spots under tissue, 55917 mean reads under tissue spot, 2275 panel genes per spot, 4544 median panel UMIs/spot, and 22508 (OCT) or 16334 (FFPE) total genes detected) (Table S2). Demultiplexing, image and transcript alignment, and transcript quantification was performed using the Spaceranger pipeline (10x Genomics, v1.2, GRCh38 reference). For comparability across OCT-preserved and FFPE-preserved samples, all samples were analyzed using the genes in the Visium FFPE gene panel. Spots were filtered to exclude those with fewer than 500 unique transcripts, and initial downstream analysis— including normalization for per-spot sequencing depth using the SCTransform algorithm, principal component analysis, UMAP layout, graph-based clustering, differential gene expression analysis, and spatially variable gene identification — was performed using the R package Seurat (V4)^90^. Previously defined gene expression modules associated with tumor and immune cell states were incorporated into this analysis using the AddModuleScore Seurat function^9,91^. Spatially variable biological processes and cell types were also inferred using clusterProfileR^92,93^.

### Annotation and analysis of histological regions

H&E images of all tissue sections were manually annotated with guidance from a board-certified neuropathologist (S.D., A.K.). In FFPE-preserved samples, regions of microvascular proliferation, necrosis, palisading necrosis, hemorrhage, and normal brain parenchyma were identified (e.g., Fig. 1, Fig S5). In OCT-preserved samples, H&E quality permitted only the identification of gross features such as regions of high and low tumor content. Tumor cell percentages were visually estimated, with extensive variation noted within and between sections. Importantly, Visium data does not have single-cell resolution; by overlaying the Visium oligonucleotide “spots” on the H&E images, we estimated that each spot contained 5-8 cells.

### Single-cell RNA-sequencing (scRNA-seq) data generation and analysis

We used the 10x Genomics 3’ v2 and 5’ v2 Chromium Gene Expression Solution to generate high-quality scRNA-seq data for 11 GBM samples (two already published^40^) from nine patients, four of whom also contributed Visium data. Because the size of tumor samples was often limiting, we could not obtain scRNA-seq samples from every patient analyzed using Visium. Sequencing reads were demultiplexed, aligned, and counted using the Cellranger 5.0 pipeline^94^. Downstream processing steps, including filtering, doublet removal, normalization, scaling, identification of variable genes, dimensionality reduction, clustering, and cell typing were performed using a WDL pipeline we developed for tailored analysis of GBM scRNA-seq data (https://dockstore.org/organizations/PettilabMGH). Briefly, doublets were identified using multiple tools, including “doubletCells”^95^,“cxds”, “bcds”, “hybrid”^96^, “scDblFinder”^97^, and “DoubletFinder”^98^. Cells predicted to be doublets by a majority of these were removed from the data. Next, cells were retained for further analysis if they met the following filtering criteria: mitochondrial gene percentage < 5%, nCount (UMIs/cell) < 93^rd^ percentile, and nFeature (unique genes/cell) > 700. Data was then log-normalized, scaled, and corrected for cell cycle gene expression using functions from Seurat V4^99^. The optimal number of Principal components (PCs) for graph-based clustering^90^ and UMAP layout was determined based on the Jackstraw test and statistical significance after Bonferoni correction. Cell types were then determined using a combination of reference-based (SingleR)^100^ and marker-based (scSorter)^101^ tools. To this end, cells were first partitioned into *CD45*+ (immune) and *CD45*-(non-immune) clusters based on *PTPRC* expression. Immune cells were further classified by training these tools on the single-cell Brain immune atlas^102^ and the immune component of the STAB^103^ atlas (adult samples only). Non-immune cells were further classified by training these tools on a combination of published^9^ GBM data and the non-immune component of STAB^103^. Putative tumor cells were further classified based on the cell states in Neftel et al^9^ using a marker-based prediction method called scSorter^101^. The malignant status of putative tumor cells was further confirmed using copy number variant (CNV) predictions obtained using CONICSmat^104^. Cells with ambiguous or noisy predictions (e.g. tumor cells without CNVs and nonmalignant cells with CNVs) were excluded from the data set.

### Generation of a brain/GBM single-cell reference for Visium data deconvolution

We combined this data with normal brain cells from the cortical dissections reported in a state-of-the-art whole-brain cell atlas^35^ and T cells from our recent single-cell study of GBM T cells^21^. The entire combined data set was normalized and scaled. Because downstream analysis steps are computationally intensive, we downsized this data by randomly sampling from within each data set. The resulting single-cell reference contained 40,000 cells and 48 cell types.

### Integration of spatial transcriptomic and scRNA-seq data

There are two classes of tools for inferring Visium spot composition, “deconvolution” tools, which estimate cell type proportions for each spot, and “cell mapping tools,” which estimate tissue coordinates for individual cells based on a reference single cell dataset. Here we used CellTrek^39^, which uses a random-forest based approach to map single cells to their most likely coordinates in Visium data. Cell mappings were consistent with marker gene expression, graph-based clustering, H&E annotations by pathologist, and other tools (such as Cell2Location^41^, data not shown). In addition, they were robust to the composition of the single-cell reference, and accurately identified tumor cells in GBM mouse models in which the location of tumor cells is unambiguous (data not shown). CellTrek allowed us to interpolate between spots to predict the cellular composition across the entire tissue slice, and and was used in two modes, “with interpolation” (for *de novo* cell state discovery) and “without interpolation” (to define multicellular niches).

### Superimposing Visium spots on mapped cells to define multicellular niches

CellTrek source code was modified to obtain single-cell mappings for each spot (without interpolation). We limited the maximum number of spots to which one cell can be charted to 10 as well as the maximum number of cells that can be mapped to a spot were limited to 8. This enabled us to interpret spot clusters as multicellular niches. Cell-to-spot mappings were then used to calculate niche composition and cell type enrichment.

### Derivation of consensus spatial tumor cell states

Spatially variable tumor cell clusters were identified using the ‘Scoexp’ module in CellTrek, which clusters mapped cells using spatially weighted gene co-expression analysis. This yielded a set of tumor cell clusters that were *approximately* conserved across patients and samples. To derive a set of consensus clusters across patients and samples, the DEGs for each tumor cluster in each sample were combined in a binary matrix of genes across samples. This matrix was clustered using a consensus clustering algorithm called cola^105^ (version 2.7.1). The ATC method was used for consensus partitioning and spherical k-means was used for sub-group classification. This approach yielded 5 major groups of DEGs that were further used to define the consensus spatial tumor cell states. To further standardize the number of genes in resulting tumor cell states, genes not present in at least 3 patients overall and at least 1 patient in both techniques (OCT & FFPE) were removed from the final set. The resulting gene set was ranked using cumulative mutual information, and the top 100 genes for each program were used to further classify tumor cells in the single cell reference dataset. Mapped cells were assigned to the state for which they scored most highly using AddModuleScore.

### Ligand receptor interaction analysis in scRNA-seq data

CellChat^106^ was used to define ligand receptor interactions to better understand how tumor cells (and cell states) communicate. CellChat defines the directionality of the interaction by calculating a communication probability based on a mass-distance algorithm. We identified interactions between various cell-type pairs (e.g. myeloid-tumor and tumor-myeloid) based on the direction of signal between the two cell types.

### Batch correction and integration of Visium data using Reciprocal Principal Component analysis

Preliminary analyses indicated that “batch correction” or “data integration” across samples increased the biological signal within clustered spots (data not shown). Therefore, Visium data from all samples was combined and batch-corrected (treating each patient as a “batch”) using Seurat’s^99^ reciprocal principal component analysis (RPCA) method. Briefly, unique variable features were identified across samples using the ‘SCT’ assay, and the merged dataset was normalized with ‘SCTransform.’ For each patient, ‘SelectIntegrationFeatures’ was applied with ‘nfeatures = 2000’. RPCA-based batch correction was then performed using FindIntegrationAnchors, leveraging the previously selected variable features. The data was integrated using ‘IntegrateData’, with k-nearest neighbors (‘k.weight’) set to 50. Statistically significant Principal Components (nPC) were selected using the Jackstraw test. The “integrated” assay of the merged data was then scaled and ‘RunPCA’ was used with nPC=63. Clustering was then performed on the resulting object using multiple resolutions. Cluster resolution 0.9 was used for further downstream analysis as it was able to capture spatial diversity as indicated in pathologist annotations.

### Cluster Adjacency

We developed a metric, Cluster Adjacency (C_A_), that reflects the probability that spots from two clusters, *i* and *j,* were “adjacent” given the frequency of each cluster and the total number of adjacent spot pairs. Here, two spots were considered adjacent if the Euclidian distance between them was ∼100μm (which is the minimum distance between 2 spots on a Visium slide). Below, *P(i)P(j)* is the probability of drawing one spot from cluster *i* and one spot from cluster *j* given the number of spots in each cluster; *P(i,j)* is the observed frequency at which spots from *i* and *j* are adjacent. Then,

Cluster Adjacency 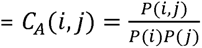

Where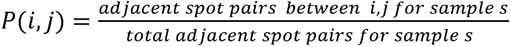

and 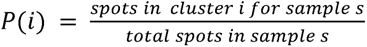 (similar for *P*(*j*))

To summarize the *CA* score across samples, we defined the following:

weighted mean adjacency 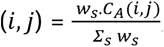, where

*w_s_*= adjacent spot pairs for *i,j* in sample s, and

*Σ_s_w_s_*= adjacent spot pairs for *i,j* across samples

The weighted mean adjacency score was calculated for each (*i,j*) combination across all RPCA cluster pairs. The cluster adjacency network was drawn using Cytoscape, where circos nodes represent the cell type distribution (from CellTrek) for each cluster. The edge thickness indicates the Weighted mean adjacency (*i,j*) between a cluster pair and the edge color/transparency indicate proportion of samples in which the cluster pair was present with minimum sample proportion set to 20%. Spatially adjacent spots for selected cluster pairs are shown in Fig. S15.

### Visium CNV analysis and spot malignancy classification

InferCNV was run for each patient using a normal brain sample from Ravi et al.^24^ as normal reference. A window length of 151 base pairs was used for smoothing; the simulation method was set to “meanvar,” with pnorm_bayes = 0.3 and analysis_mode set to “subclusters”. CNV partitioning was done using “leiden algorithm” at “clustering resolution=0.05”. The resulting tumor state output of HMM inferred CNV states for each CNV region was then used to assign a CNV score (CS*_i_*) using an approach similar to that described in Xun et al^107^: *CS*_*i*_ = Σ_*j*_ │*cs_ij_* – 3│ where gene *j* is in spot *i*, and CNV states are defined as in the InferCNV HMM wiki (https://github.com/broadinstitute/infercnv/wiki/inferCNV-HMM-based-CNV-Prediction-Methods), with state 3 as neutral, values >3 indicating amplification, and <3 indicating deletion for each gene. Spots with CNV scores (CSi) greater than +1 standard deviation (s.d.) above the mean were classified as having highly malignancy, those below −1 s.d. as low malignancy, and the remainder as intermediate malignancy. Malignant CNV subclusters were further consolidated to a maximum of 20 subclusters per patient based on the similarity of their CNV state assignment vectors.

### Ligand receptor interaction analysis in Visium data

To analyze the spatial variation in ligand-receptor (LR) interactions of Visium data at per-spot resolution, stLearn^108^ was utilized with ligand-receptor pairs obtained from the Cellchat V2^109^ database. For contact-dependent interactions, calculations were performed *within spot mode*, while for "Secreted Signaling" and "Extracellular Matrix Remodeling" interactions (diffusion-dependent interactions), statistically significant spot pairs were identified over a given distance. It has been shown that different LR pairs interact across varying distance ranges^110^. To identify the optimal set of significant spot pairs, multiple distance parameters were explored, and a distance of 2000 μm was selected. Statistically significant contact-dependent LR pairs for each spot were identified using stLearn, and their enrichment within each cluster was assessed using the hypergeometric distribution. For contact dependent interactions if they were significant for at least 10 spots in a cluster and 100 spots over all were used in further analysis.Hypergeometric test was done for calculating significance of enrichment for each LR pair in each cluster, where:

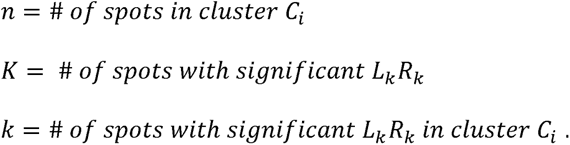

*p*-values were corrected using the Bonferroni correction. The top five contact-dependent LR pairs for each cluster are shown in Fig. 4B, while all significant LR pairs meeting the above threshold are provided in the Supplementary Data. For LR analysis over a distance, stLearn reports significant spots for each LR pair. To distinguish a sender spot with Ligand L_k_ from a receiver spot with Receptor R_k_ (where k indicates a known interaction based on Cellchat V2 database) for significant spots over a distance, spots with L_k_ expression ≥ 2 were classified as sender spot/s similarly, spot with expression R_k_ ≥ 2 were designated as receiver spots. For all directionally unique spot pairs within/between any pair of batch-corrected clusters, we calculated the LR connectivity score as follows:

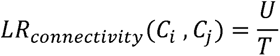

Where:

*U = Unique sender and receiver spot pairs significant for at least one LR pair between Ci and Cj*

*T = Total possible spot pairs involving Ci (sender) and Cj (receiver)*

LR connectivity is shown in Fig. 4C, with thickness indicating the connectivity score and color indicating the number of samples where the cluster pairs interact.

To further understand enrichment of certain LR pairs in inter-niche (inter-Cluster) and intra-niche (intra-Cluster) LR signaling between a cluster-pair or between two groups (with groups being edge/interface/non-specific/core) we conducted a hypergeometric test of enrichment for each LR pair (L_k_R_k_) such that

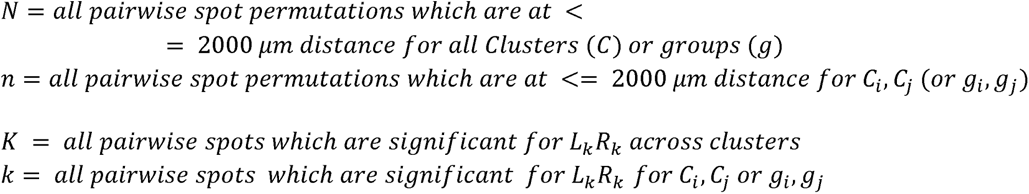

The resulting *p*-values for each analysis were corrected using the Bonferronni correction. Significant LRs and their enrichment scores meeting the threshold criteria and present in at least 3 samples and/or >2 patients are shown (Fig. S19).

### Xenium panel design

We designed a panel of oligonucleotides targeting transcripts from 362 genes, comprising 266 genes from the predesigned 10X Genomics human brain-and-glioblastoma panel and 96 custom genes indicated by our scRNA-seq analyses (Table S7). Together, this customized panel captures diverse neurons, glia, endothelial cells, microglia, immune cells, GBM cells (including both spatial GBM cell states derived from our Visium data and published scRNA-seq-derived GBM cell states), and ligand-receptor pairs that putatively mediate tumor/neuron, tumor/T cell, and tumor/macrophage interactions.

### Xenium In Situ Workflow

#### Cohorts

Xenium data was collected in three cohorts. Key analyses were performed using the primary and secondary cohorts, which did not overlap the Visium cohort. The tertiary Xenium cohort was comprised of six tissue sections, each taken from the same block as a Visium sample, and was primarily used for technical comparisons with Visium.

#### Sample Preparation - Primary Cohort and Tertiary cohort

Formalin-fixed, paraffin-embedded (FFPE) tumor samples were sectioned into 5-um sections using a microtome onto Xenium slides according to Protocol CG000578 published by 10X Genomics. The Xenium slides were then deparaffinized and de-crosslinked according to the Protocol CG000580. Both a pre-designed gene expression panel (Human Brain Probe set) and a custom probe set were used during probe hybridization, ligation, and amplification according to Protocol CG000749, which also contained cell segmentation staining. The Xenium slide was then loaded into the Xenium Analyzer using Protocol CG000584 and regions of interest (ROIs) were initially selected using the on-instrument software.

#### Sample Preparation - Secondary Cohort

Samples were processed similarly as previously published^21^. In brief, formalin-fixed, paraffin-embedded (FFPE) tumor samples were sectioned into 5-um sections using a microtome onto Xenium slides according to Protocol CG000578 published by 10X Genomics. The Xenium slides were then deparaffinized and decrosslinked according to the Protocol CG000580. Both a pre-designed gene expression panel (Human Brain Probe set) and a custom probe set were used during probe hybridization, ligation, and amplification according to Protocol CG000582. (NB: Cell segmentation staining had not yet been introduced by 10X Genomics at the time this data was generated.) The Xenium slide was then loaded into the Xenium Analyzer using Protocol CG000584 and regions of interest (ROIs) were initially selected using the on-instrument software.

#### Sample Filtering and Cell Type Identification - Primary Cohort

Raw FASTQ files were analyzed in Xenium Ranger (version 2.0.0.10) using the Xenium Multi-Tissue Stain cell segmentation method. Data were then processed using the Seurat R package (5.0.1). Initial filtering and subsequent analysis were performed on a per-sample basis. Counts were then log-normalized and PCA was performed with the optimal number of PCs determined based upon elbow plots, jackstraw resampling, and PC expression heatmaps. Dimensionality reduction and visualization were performed with the UMAP algorithm (Seurat implementation) followed by unsupervised graph-based clustering. Clusters with overall low nCount and nFeature were removed, and PCA and UMAP visualization was performed again on the filtered object. Cell types in each sample were individually characterized using marker genes and differentially expressed genes (Table S9-S10). Samples were then merged using *SCTransform* (Seurat implementation) and PCA and UMAP visualization was performed again on the merged object. Cell types were then standardized by performing hierarchical clustering on different groups of cell types, such as immune cells, tumor cells, etc. Cell types with consistent gene expression profiles were labeled similarly and differentially expressed genes of each cell type were analyzed to confirm correct identification.

Visium-based tumor signature scores were generated using *AddModuleScore* (Seurat implementation) and applied to patient-specific clusters within the primary cohort to identify tumor cell identities. All patient-specific clusters were subsequently hierarchically clustered according to Visium-based tumor signature scores and appropriately classified.

#### Sample Filtering and Cell Type Identification - Secondary Cohort

Raw FASTQ files were reprocessed in Xenium Ranger (version 1.7.0.2) with an expansion distance of 0um. Data were then processed using the Seurat R package (5.0.1). Initial filtering excluded cells that had any of the following: an nFeature count less than 5, an nFeature count greater than 125, or an nCount value greater than 400. Samples were then merged and down-sampled to 50,000 cells using SketchData (Seurat). Counts were then log-normalized and PCA was performed with the optimal number of PCs determined based upon elbow plots, jackstraw resampling, and PC expression heatmaps. Dimensionality and visualization were performed with the UMAP algorithm (Seurat implementation) followed by unsupervised graph-based clustering. Clusters with overall low nCount and nFeature were removed, and PCA and UMAP visualization was performed again on the filtered object. To label cells, the primary cohort Seurat object was first log-normalized and PCA was applied. Then, FindTransferAnchors and TransferData was performed using the primary cohort Seurat object as a reference and applied to the secondary cohort Seurat object. Similarity between gene expression profiles of primary and secondary cohort cell types was assessed by performing a correlation analysis.

#### Xenium Niche Analysis

Niche analysis was performed using the BANKSY R package^44^. On a per sample basis, *RunBanksy* was performed on log-normalized data with a lambda value of 1.0 for both primary and secondary cohort samples. Relative proportions of each cell type per sample-specific niche were then aggregated and log-normalized. Hierarchical clustering was then performed and niches were appropriately classified based upon cell type composition. Statistical significance for cell type enrichment per niche was calculated using the hypergeometric test.

#### Xenium Enrichment Score

Enrichment score was calculated by summing the relative frequency of a target cell state within a specific radius (or radial range) of a source cell for all source cells, scaled to the overall frequency of the particular cell state.

#### Xenium Niche Distance Analysis

Average minimum distances between niches were quantified by first calculating the distance between a cell of a given primary niche (Niche A) to its nearest neighbor in a different, secondary niche (Niche B) within a given sample. This distance was labeled as the “minimum distance” and calculated for every cell in a primary niche (Niche A), on a per-sample basis, and averaged to calculate the minimum distance from Niche A to Niche B for a given sample. This was then averaged across samples to give the final average minimum distance. The node-edge map was then generated using *layout_with_kk* (igraph package^111^), which generates a layout using the Kamada-Kawai algorithm.

#### Xenium Ligand Receptor Analysis

Ligand receptor analysis was performed using the CellChat v2 R package^109^ on a per-sample basis. Significant ligand-receptor (LR) pairs between every cell type across samples were aggregated. For cell types *A* and *B* and LR *i*, we considered *A* to interact with *B* via *i* if the interaction was observed in more than 3 samples. Then, for each cell type pair, we counted unique *i* and total *i*.The total number of ligand-receptor interactions were quantified by summing the number of statistically significant ligand-receptor interactions across samples between two given pairs of cell types. The number of unique ligand-receptor interactions was quantified by summing the number of unique interactions across samples between two given pairs of cell types.

### Integrated Cross-Platform Ligand-Receptor Interaction analysis

We used CellChat V2^109^ to analyze Xenium and single-cell reference (scRef) data to infer cell-cell communication. For both datasets, we collated ligand-receptor interactions at the level of broad cell types, which included Myeloid, Lymphoid, Vascular, Glia, and Neuron, but preserved distinct tumor states as specific sender and receiver identities. Communication probabilities were aggregated for each combination of broad cell type or tumor state and L-R pair across all Xenium samples, yielding a mean probability for each sender-receiver-ligand-receptor (L-R) combination hereon defined as cell–cell communication event (CCE). The same procedure was applied to the scRef dataset, resulting in a mean communication probability for each CCE for both datasets.

For the Visium data, contact-dependent ligand-receptor (L-R) analysis was conducted within individual spots, while diffusible L-R analysis was performed between spots, maintaining a maximum distance of 2000 μm. Complex ligand-receptor interactions from the CellChat V2 database were simplified into individual pairs for use with Stlearn^108^, which only supports non-complex interactions. (A complex L-R interaction was considered significant if all its components were significant for the respective spot or spot-pair. For example, the *ADM-CALCR-RAMP2* interaction was deemed significant if both *ADM-CALCR* and *ADM-RAMP2* were significant for that spot pair.) We further calculated the total number of spots or spot pairs for each domain-L-R combination. An L-R interaction was classified as "Core-enriched" if the Log_2_ fold change (FC) between core and edge spots was ≥2, and "Edge-enriched" if Log_2_FC ≤ -2. Interactions that didn’t meet these criteria were labeled as "Not-Domain specific." We also recorded the number of distinct patients and samples for each L-R interaction across Visium data.

We utilized CellChat v2 to analyze nearly half of all the GBMap data, validating our findings from other assays and our single-cell reference. We summarized the average communication probability between major cell types (e.g., Tumor, Myeloid, etc.) for each ligand-receptor pair across all combinations of broad sender and receiver cell types. Since GBMap did not provide tumor-state resolution, all tumor-state-specific interactions identified in the Xenium– scRef dataset were grouped into the general "Tumor" category for merging with GBMap. As a result, if a specific ligand-receptor interaction was found between Tumor and a particular broad cell class in GBMap, that interaction was combined with all corresponding tumor-state-specific interactions involving the same ligand-receptor pair and broad cell class in the Xenium–scRef communication table. Finally, the resulting file was merged with the Visium output described above.

The final set of cross-platform LR interactions were required to meet the following criteria: present in ≥2 independent evidence sources, ≥1 spatial platform (Visium or Xenium), ≥1 single-cell resolution platform (scRef, GBMap, or Xenium), with Xenium-only interactions excluded.

### NicheNet analysis

To identify the target genes of the inferred cell-cell interactions, ligand-based target prediction was performed using the NicheNet (v2.2.1.1). For each sender cell type, interactions showing above-average interaction probabilities across multiple platforms, including scRNA-seq and Xenium-based interaction analyses, were defined as major interactions. Ligands involved in these major interactions were selected for downstream NicheNet analysis. For each ligand, we first selected the top 10% of target genes with regulatory potential scores. After applying the ligand-target link extraction and visualization cutoff, we further retained the top 20% of targets based on the final ligand-target weight. To prioritize biologically relevant target genes within cancer cells, candidate targets were selected using two criteria: (1). the top 500 most highly expressed genes among genes expressed in at least 30% of cells across all five cancer cell subtypes (2). cancer subtype-specific marker genes, which were identified using the Seurat FindAllMarkers (|logFC | >= 1, min.pct = 0.3, min.diff.pct = 0.1, and p_val_adj < 0.01). For sender-specific NicheNet analyses, interactions that were primarily observed in a given sender cell types were first isolated. For each ligand, target genes with top 10% regulatory potential scores were selected. Candidate target genes were then defined using the same subtype-shared and subtype-specific criteria described above. The results were visualized using the pheatmap (v1.0.13), with target gene expression represented as the aggregated average expression levels across each tumor subtype.

### ISH-IF (RNAscope)

#### Tissue Staining

Six FFPE tissue sections from mice GBM samples were stained using a Leica Bond RX automated stainer. Three staining panels were developed using RNAscope ISH probes (Advanced Cell Diagnostics) with Opal fluorophores (Akoya Biosciences, a Quanterix Company). The first panel consisted of 5 RNA probes (Hs-CD44, Hs-CALCRL, Hs-ADM, Hs-PECAM1, and Hs-SOX2) and 1 antibody CD45 (Leica Bioscience LCA-L-U). The second panel consisted of 5 RNA probes (Hs-PDGFRB, Hs-HES1, Hs-PECAM1, and Hs-SOX2) and 2 antibodies JAG1(Jagged1 (D4Y1R) Rabbit Monoclonal Antibody #70109) and Notch-3 Antibody (Novus Biologicals Notch-3 Antibody (1G5) - Azide and BSA Free H00004854-M01. The third panel consisted of 5 RNA probes (Hs-CD44, Hs-PTN, Hs-NCL, Hs-PECAM1, and Hs-SOX2) and 1 antibody CD45 (Leica Bioscience LCA-L-U). Tyramide Signal Amplification (TSA) was used to boost fluorophore signal (Opal 480, Opal 520, Opal 570, Opal 620, Opal 690, and Opal 780). Opal fluorophore concentrations were optimized to balance signal intensity across all channels. DAPI was used as a nuclear counterstain.

#### Image Acquisition

Whole slide images were acquired on an Akoya PhenoImager HT multi-spectral slide scanner at 40x (0.26 um/pixel) resolution. Exposures were set for each image individually to avoid pixel saturation or underexposure. Akoya inForm software was used to spectrally separate signals from each fluorophore and to remove native tissue autofluorescence. Unmixed tiles produced by inForm were stitched to reproduce a whole-slide pyramidal TIF using Indica Labs HALO software. Matched hematoxylin and eosin stained slides were scanned at 40x (0.13um/pixel) resolution using a MoticEasyScan Infinity digital pathology scanner.

#### Image Analysis

Quantitative image analysis was performed using the HALO image analysis platform (Indica Labs). Each sample image was manually-annotated to exclude areas of debris, tissue folds. Cell segmentation and phenotyping was performed across the entire tissue regions using HALO FISH-IF and AI modules. Nuclear detection and cell segmentation were performed using the default AI nuclear segmentation algorithm based on the DAPI channel. Cells were phenotyped based on signal intensity of the antibody channel within the cell. RNA-ISH positivity was based on signal intensity and dot size within a cell, with brighter and larger dots having more weight. The positivity thresholds were manually validated for each sample for cell phenotyping quality. Summary tables containing cell phenotype information were exported for further analysis across all samples. HALO Nearest Neighbor Spatial Analysis was performed on the phenotyped cells to determine the average distance from each index cell to the nearest phenotype cell of interest.

### Multiplexed Immunofluorescence

#### Tissue preparation and antigen retrieval

Formalin-fixed and paraffin-embedded (FFPE) tissue samples were sectioned at 4μm thickness onto standard histopathology slides by the Brigham and Women’s Pathology core. Immediately prior to staining, slides were de-paraffinized in two consecutive xylene baths for 5 minutes each at room temperature with agitation. This was followed by two consecutive 100% ethanol baths for 2 minutes each with agitation. Slides were baked in a laboratory oven at 60C for 5 minutes. Antigen retrieval was performed on a BOND RX Automated Research Stainer using the ER2 solution at 98C for 1 hour (Leica Biosystems). Slides were briefly stored in Multistaining Buffer (Lunaphore Technologies SA, BU06) before being loaded onto the COMET machine (Lunaphore Technologies SA).

#### Immunofluorescent staining

Sequential immunofluorescent staining was performed on the COMET platform via 10 automated sequential cycles of staining, imaging, and elution of antibodies as previously described (1). In brief, slides were stained with primary antibodies and subsequently incubated with a mixture of secondary antibodies and DAPI. After antibody incubation, fluorescent images were captured in the DAPI, TRITC, and Cy5 channels at standard exposure times set by the manufacturer. Following the imaging step, antibodies were eluted via incubation with Elution Buffer (Lunaphore Technologies SA, BU07). All reagent dilutions and incubation times were optimized prior to staining. A complete list of antibodies and staining conditions can be found in Table S17.

#### Immunofluorescent image analysis

During the sequential immunofluorescent staining protocol, images were automatically registered, stitched together, and compiled into OME.TIFFs by the COMET acquisition software. Image analysis was carried out using the HORIZON software (Lunaphore Technologies SA). An autofluorescence image was subtracted from each channel to reduce background. 7-9 rectangular regions of interest (ROIs) were selected per sample to represent the whole tissue. Each ROI is approximately approximately 430μm x 360μm. Nuclear segmentation was performed using DAPI signal with the StarDIST algorithm. Positivity for an individual marker was determined by a threshold on the mean fluorescent intensity of the nucleus or whole cell for nuclear and cytoplasmic/membranous markers respectively. Thresholds were set independently for each sample to account for variations in tissue and staining quality.

#### Immunofluorescence (non-multiplexed)

Formalin-fixed, paraffin-embedded (FFPE) tumor samples were sectioned into 5-um sections and baked at 55°C for 2 hours. Slides were then deparaffinized in: xylene for 10 minutes (twice), 100% ethanol for 5 minutes (twice), 90% ethanol for 3 minutes (once), 70% ethanol for 3 minutes (once), and PBS for 5 minutes (twice). Slides were then incubated in 1X Tris-EDTA buffer, pH 9 (Abcam) at 95C for 30 minutes and then cooled in antigen retrieval buffer for 25 minutes. Slides were then washed in PBS for 5 minutes and subsequently incubated in blocking buffer (1X PBS with 0.1% Triton X-100, 5% BSA, and 5% donkey serum). Primary antibodies (diluted in blocking buffer) were used as follows: goat anti-LYVE1 antibody 1:30 (AF2089, RND Systems), rabbit anti-CD163 1:500 (93498, Cell Signaling Technology). Tissues were stained with primary antibody solution overnight at 4°C and then washed in PBS for 10 minutes thrice. Secondary antibodies (diluted in blocking buffer) were used as follows: AlexaFluor 488 donkey anti-goat 1:500 (Jackson ImmunoResearch), AlexaFluor 647 donkey anti-rabbit 1:500 (Jackson ImmunoResearch). Slides were incubated in secondary antibody solution at room temperature for 1 hour and then washed in PBS for 10 minutes thrice. Slides were then stained with DAPI at a final concentration of 5 μg/mL for 30 minutes and then mounted with VectaShield Vibrance mounting medium (VectorLabs). Images were acquired by fluorescence microscopy with a Nikon AXR microscope.

## RESOURCE AVAILABILITY

### Lead contact

Allegra A. Petti

### Materials availability

Standardized reagents are available from the indicated suppliers.

## Data and code availability

Software used for analysis is public and described in the Methods section. Additional code used in the analysis is available at https://github.com/Petti-Lab/GBM_Spatial_paper_figures_code/ or https://github.com/smk5g5/GBM_Spatial_paper_figures_code

## ACKNOWLEDGEMENTS.

This work was supported by a Cancer Research Foundation Young Investigator’s Award (to A.A.P.), National Institutes of Health grant R21CA286215 (to A.A.P.), an American Brain Tumor Association Discovery Grant (to A.A.P.), National Institutes of Health grants R01 NS094670, R01 NS128470 (to A.H.K.), the Christopher Davidson and Knight Family Fund (to A.H.K.); and the Duesenberg Research Fund (to A.H.K.). We thank GTAC-MGI at Washington University School of Medicine in St. Louis for generating Xenium data, and Jonathan L. Weinstein for assistance with tissue blocks for mIF experiments. During the preparation of this work, the authors used Claude in two ways: to improve the brevity and clarity of the writing, and to write Python code used to create plots (Fig. 4A, Fig. S16) of the cross-platform ligand-receptor interactions we identified as described above. After using this tool/service, the author(s) reviewed and edited the content as needed and take(s) full responsibility for the content of the published article.

## AUTHOR CONTRIBUTIONS

Conceptualization, A.A.P., G.P.D., A.H.K.; Data curation A.A.P., R.R.D., S.M.K., A.Z.W., C.R.M.; Formal analysis, S.M.K., A.Z.W., A.A.P., G.D.K., C.R.M., J.F-S., A.F.W., Y.S., E.S., H.O.; Funding acquisition, A.A.P., G.P.D., A.H.K.; Investigation, R.R.D., A.Z.W., G.P.D., A.H.K., R.S., S.M.D., M.A., J.M., N.D.S., L.L., T.W., A.K., B.D.C., E.C.L., B.P., K.O., H.O., Y.S.; Methodology, A.A.P., G.P.D., A.H.K., R.R.D., S.M.K., A.Z.W., G.D.K., R.S., N.P.G., K.P., L.N.; Resources, S.M.K., A.Z.W., R.R.D.; Software, S.M.K., A.Z.W.; Supervision, A.A.P., G.P.D.; Writing-original draft, A.A.P., A.Z.W.; Writing-review & editing, S.M.K., G.D.K., A.C., A.H.K., G.P.D., R.R.D.

## REFERENCES

1. Weller, M., Butowski, N., Tran, D.D., Recht, L.D., Lim, M., Hirte, H., Ashby, L., Mechtler, L., Goldlust, S.A., Iwamoto, F., et al. (2017). Rindopepimut with temozolomide for patients with newly diagnosed, EGFRvIII-expressing glioblastoma (ACT IV): a randomised, double-blind, international phase 3 trial. Lancet Oncol. 18, 1373–1385.

2. Liau, L.M., Ashkan, K., Brem, S., Campian, J.L., Trusheim, J.E., Iwamoto, F.M., Tran, D.D., Ansstas, G., Cobbs, C.S., Heth, J.A., et al. (2023). Association of autologous tumor lysate-loaded dendritic cell vaccination with extension of survival among patients with newly diagnosed and recurrent glioblastoma: A phase 3 prospective externally controlled cohort trial: A phase 3 prospective externally controlled cohort trial. JAMA Oncol. 9, 112–121.

3. Omuro, A., Brandes, A.A., Carpentier, A.F., Idbaih, A., Reardon, D.A., Cloughesy, T., Sumrall, A., Baehring, J., van den Bent, M., Bähr, O., et al. (2023). Radiotherapy combined with nivolumab or temozolomide for newly diagnosed glioblastoma with unmethylated MGMT promoter: An international randomized phase III trial. Neuro. Oncol. 25, 123–134.

4. Cancer Genome Atlas Research Network (2008). Comprehensive genomic characterization defines human glioblastoma genes and core pathways. Nature 455, 1061–1068.

5. Brennan, C.W., Verhaak, R.G.W., McKenna, A., Campos, B., Noushmehr, H., Salama, S.R., Zheng, S., Chakravarty, D., Sanborn, J.Z., Berman, S.H., et al. (2013). The somatic genomic landscape of glioblastoma. Cell 155, 462–477.

6. Schaettler, M.O., Richters, M.M., Wang, A.Z., Skidmore, Z.L., Fisk, B., Miller, K.E., Vickery, T.L., Kim, A.H., Chicoine, M.R., Osbun, J.W., et al. (2022). Characterization of the Genomic and Immunologic Diversity of Malignant Brain Tumors through Multisector Analysis. Cancer Discov. 12, 154–171.

7. Verhaak, R.G.W., Hoadley, K.A., Purdom, E., Wang, V., Qi, Y., Wilkerson, M.D., Miller, C.R., Ding, L., Golub, T., Mesirov, J.P., et al. (2010). Integrated genomic analysis identifies clinically relevant subtypes of glioblastoma characterized by abnormalities in PDGFRA, IDH1, EGFR, and NF1. Cancer Cell 17, 98–110.

8. Wang, Q., Hu, B., Hu, X., Kim, H., Squatrito, M., Scarpace, L., Decarvalho, A.C., Lyu, S., Li, P., Li, Y., et al. (2017). Tumor Evolution of Glioma-Intrinsic Gene Expression Subtypes Associates with Immunological Changes in the Microenvironment. Cancer Cell 32, 42–56.e6.

9. Neftel, C., Laffy, J., Filbin, M.G., Hara, T., Shore, M.E., Rahme, G.J., Richman, A.R., Silverbush, D., Shaw, M.L., Hebert, C.M., et al. (2019). An Integrative Model of Cellular States, Plasticity, and Genetics for Glioblastoma. Cell 178, 835–849.e21.

10. Patel, A.P., Tirosh, I., Trombetta, J.J., Shalek, A.K., Gillespie, S.M., Wakimoto, H., Cahill, D.P., Nahed, B.V., Curry, W.T., Martuza, R.L., et al. (2014). Single-cell RNA-seq highlights intratumoral heterogeneity in primary glioblastoma. Science 344, 1396–1401.

11. Bhaduri, A., Di Lullo, E., Jung, D., Müller, S., Crouch, E.E., Espinosa, C.S., Ozawa, T., Alvarado, B., Spatazza, J., Cadwell, C.R., et al. (2020). Outer Radial Glia-like Cancer Stem Cells Contribute to Heterogeneity of Glioblastoma. Cell Stem Cell 26, 48–63 e6.

12. Couturier, C.P., Ayyadhury, S., Le, P.U., Nadaf, J., Monlong, J., Riva, G., Allache, R., Baig, S., Yan, X., Bourgey, M., et al. (2020). Single-cell RNA-seq reveals that glioblastoma recapitulates a normal neurodevelopmental hierarchy. Nat. Commun. 11, 3406.

13. Wang, L., Babikir, H., Müller, S., Yagnik, G., Shamardani, K., Catalan, F., Kohanbash, G., Alvarado, B., Di Lullo, E., Kriegstein, A., et al. (2019). The phenotypes of proliferating glioblastoma cells reside on a single axis of variation. Cancer Discov. 9, 1708–1719.

14. Garofano, L., Migliozzi, S., Oh, Y.T., D’Angelo, F., Najac, R.D., Ko, A., Frangaj, B., Caruso, F.P., Yu, K., Yuan, J., et al. (2021). Pathway-based classification of glioblastoma uncovers a mitochondrial subtype with therapeutic vulnerabilities. Nat. Cancer 2, 141–156.

15. Darmanis, S., Sloan, S.A., Croote, D., Mignardi, M., Chernikova, S., Samghababi, P., Zhang, Y., Neff, N., Kowarsky, M., Caneda, C., et al. (2017). Single-Cell RNA-Seq Analysis of Infiltrating Neoplastic Cells at the Migrating Front of Human Glioblastoma. Cell Rep. 21, 1399–1410.

16. Nomura, M., Spitzer, A., Johnson, K.C., Garofano, L., Nehar-Belaid, D., Galili Darnell, N., Greenwald, A.C., Bussema, L., Oh, Y.T., Varn, F.S., et al. (2025). The multilayered transcriptional architecture of glioblastoma ecosystems. Nat. Genet. 57, 1155–1167.

17. Brooks, W.H., Netsky, M.G., Normansell, D.E., and Horwitz, D.A. (1972). Depressed cell-mediated immunity in patients with primary intracranial tumors. Characterization of a humoral immunosuppressive factor. J. Exp. Med. 136, 1631–1647.

18. Chongsathidkiet, P., Jackson, C., Koyama, S., Loebel, F., Cui, X., Farber, S.H., Woroniecka, K., Elsamadicy, A.A., Dechant, C.A., Kemeny, H.R., et al. (2018). Sequestration of T cells in bone marrow in the setting of glioblastoma and other intracranial tumors. Nat. Med. 24, 1459–1468.

19. Mathewson, N.D., Ashenberg, O., Tirosh, I., Gritsch, S., Perez, E.M., Marx, S., Jerby-Arnon, L., Chanoch-Myers, R., Hara, T., Richman, A.R., et al. (2021). Inhibitory CD161 receptor identified in glioma-infiltrating T cells by single-cell analysis. Cell 184, 1281–1298.e26.

20. Naulaerts, S., Datsi, A., Borras, D.M., Antoranz Martinez, A., Messiaen, J., Vanmeerbeek, I., Sprooten, J., Laureano, R.S., Govaerts, J., Panovska, D., et al. (2023). Multiomics and spatial mapping characterizes human CD8^+^ T cell states in cancer. Sci. Transl. Med. 15, eadd1016.

21. Wang, A.Z., Mashimo, B.L., Schaettler, M.O., Sherpa, N.D., Leavitt, L.A., Livingstone, A.J., Khan, S.M., Li, M., Anzaldua-Campos, M.I., Bradley, J.D., et al. (2024). Glioblastoma-Infiltrating CD8+ T Cells Are Predominantly a Clonally Expanded GZMK+ Effector Population. Cancer Discov. 14, 1106–1131.

22. Xuan, W., Lesniak, M.S., James, C.D., Heimberger, A.B., and Chen, P. (2021). Context-Dependent Glioblastoma-Macrophage/Microglia Symbiosis and Associated Mechanisms. Trends Immunol. 42, 280–292.

23. Ravi, V.M., Neidert, N., Will, P., Joseph, K., Maier, J.P., Kückelhaus, J., Vollmer, L., Goeldner, J.M., Behringer, S.P., Scherer, F., et al. (2022). T-cell dysfunction in the glioblastoma microenvironment is mediated by myeloid cells releasing interleukin-10. Nat. Commun. 13, 925.

24. Ravi, V.M., Will, P., Kueckelhaus, J., Sun, N., Joseph, K., Salié, H., Vollmer, L., Kuliesiute, U., von Ehr, J., Benotmane, J.K., et al. (2022). Spatially resolved multi-omics deciphers bidirectional tumor-host interdependence in glioblastoma. Cancer Cell 40, 639–655.e13.

25. Greenwald, A.C., Darnell, N.G., Hoefflin, R., Simkin, D., Mount, C.W., Gonzalez Castro, L.N., Harnik, Y., Dumont, S., Hirsch, D., Nomura, M., et al. (2024). Integrative spatial analysis reveals a multi-layered organization of glioblastoma. Cell 187, 2485–2501.e26.

26. Ren, Y., Huang, Z., Zhou, L., Xiao, P., Song, J., He, P., Xie, C., Zhou, R., Li, M., Dong, X., et al. (2023). Spatial transcriptomics reveals niche-specific enrichment and vulnerabilities of radial glial stem-like cells in malignant gliomas. Nat. Commun. 14, 1028.

27. Hara, T., Chanoch-Myers, R., Mathewson, N.D., Myskiw, C., Atta, L., Bussema, L., Eichhorn, S.W., Greenwald, A.C., Kinker, G.S., Rodman, C., et al. (2021). Interactions between cancer cells and immune cells drive transitions to mesenchymal-like states in glioblastoma. Cancer Cell 39, 779–792.e11.

28. Moffet, J.J.D., Fatunla, O.E., Freytag, L., Kriel, J., Jones, J.J., Roberts-Thomson, S.J., Pavenko, A., Scoville, D.K., Zhang, L., Liang, Y., et al. (2023). Spatial architecture of high-grade glioma reveals tumor heterogeneity within distinct domains. Neurooncol. Adv. 5, vdad142.

29. Lin, J., Chen, C., Li, S., Chen, L., Fang, M., Fu, S., Chen, K., Xu, H., Zhang, F., Wang, R., et al. (2026). Spatial and single-cell characterization of human glioblastoma tumor microenvironment reveals malignant cellular communities. Nat. Neurosci., 1–13.

30. Harwood, D.S.L., Pedersen, V., Bager, N.S., Schmidt, A.Y., Stannius, T.O., Areškevičiūtė, A., Josefsen, K., Nørøxe, D.S., Scheie, D., Rostalski, H., et al. (2024). Glioblastoma cells increase expression of notch signaling and synaptic genes within infiltrated brain tissue. Nat. Commun. 15, 7857.

31. Zheng, Y., Carrillo-Perez, F., Pizurica, M., Heiland, D.H., and Gevaert, O. (2023). Spatial cellular architecture predicts prognosis in glioblastoma. Nat. Commun. 14, 4122.

32. Migliozzi, S., Adabbo, B., Garofano, L., Wu, F., Davila, P., Komotar, R.J., Ivan, M.E., Shah, A.H., Currall, B.B., Williams, S., et al. (2025). Restraint of cancer cell plasticity by spatial homotypic clustering. Cancer Cell 43, 2206–2223.e10.

33. Mathur, R., Wang, Q., Schupp, P.G., Nikolic, A., Hilz, S., Hong, C., Grishanina, N.R., Kwok, D., Stevers, N.O., Jin, Q., et al. (2024). Glioblastoma evolution and heterogeneity from a 3D whole-tumor perspective. Cell 187, 446–463.e16.

34. De Bonis, P., Anile, C., Pompucci, A., Fiorentino, A., Balducci, M., Chiesa, S., Lauriola, L., Maira, G., and Mangiola, A. (2013). The influence of surgery on recurrence pattern of glioblastoma. Clin. Neurol. Neurosurg. 115, 37–43.

35. Siletti, K., Hodge, R., Mossi Albiach, A., Lee, K.W., Ding, S.-L., Hu, L., Lönnerberg, P., Bakken, T., Casper, T., Clark, M., et al. (2023). Transcriptomic diversity of cell types across the adult human brain. Science 382, eadd7046.

36. Ruiz-Moreno, C., Salas, S.M., Samuelsson, E., Minaeva, M., Ibarra, I., Grillo, M., Brandner, S., Roy, A., Forsberg-Nilsson, K., Kranendonk, M.E.G., et al. (2025). Charting the single-cell and spatial landscape of IDH-wild-type glioblastoma with GBmap. Neuro. Oncol. 27, 2281–2295.

37. Megill, C., Martin, B., Weaver, C., Bell, S., Prins, L., Badajoz, S., McCandless, B., Pisco, A.O., Kinsella, M., Griffin, F., et al. (2021). Cellxgene: A performant, scalable exploration platform for high dimensional sparse matrices. bioRxiv, 2021.04.05.438318. 10.1101/2021.04.05.438318.

38. Hadjipanayis, C.G., Widhalm, G., and Stummer, W. (2015). What is the surgical benefit of utilizing 5-aminolevulinic acid for fluorescence-guided surgery of malignant gliomas? Neurosurgery 77, 663–673.

39. Wei, R., He, S., Bai, S., Sei, E., Hu, M., Thompson, A., Chen, K., Krishnamurthy, S., and Navin, N.E. (2022). Spatial charting of single-cell transcriptomes in tissues. Nat. Biotechnol. 40, 1190–1199.

40. Mahlokozera, T., Taiwo, R., Patel, B., Hafez, D., Paturu, M., Salehi, A., Mao, D.D., Gujar, A., Dunn, G.P., Mosammaparast, N., et al. (2021). Competitive binding of E3 ligases TRIM26 and WWP2 controls SOX2 proteostasis in glioblastoma. Under Revision.

41. Kleshchevnikov, V., Shmatko, A., Dann, E., Aivazidis, A., King, H.W., Li, T., Elmentaite, R., Lomakin, A., Kedlian, V., Gayoso, A., et al. (2022). Cell2location maps fine-grained cell types in spatial transcriptomics. Nat. Biotechnol. 40, 661–671.

42. Wang, L., Jung, J., Babikir, H., Shamardani, K., Jain, S., Feng, X., Gupta, N., Rosi, S., Chang, S., Raleigh, D., et al. (2022). A single-cell atlas of glioblastoma evolution under therapy reveals cell-intrinsic and cell-extrinsic therapeutic targets. Nat. Cancer 3, 1534– 1552.

43. Jackson, C., Cherry, C., Bom, S., Dykema, A.G., Wang, R., Thompson, E., Zhang, M., Li, R., Ji, Z., Hou, W., et al. (2025). Distinct myeloid-derived suppressor cell populations in human glioblastoma. Science 387, eabm5214.

44. Singhal, V., Chou, N., Lee, J., Yue, Y., Liu, J., Chock, W.K., Lin, L., Chang, Y.-C., Teo, E.M.L., Aow, J., et al. (2024). BANKSY unifies cell typing and tissue domain segmentation for scalable spatial omics data analysis. Nat Genet 56, 431–441.

45. Tomaszewski, W., Sanchez-Perez, L., Gajewski, T.F., and Sampson, J.H. (2019). Brain tumor microenvironment and host state: Implications for immunotherapy. Clin. Cancer Res. 25, 4202–4210.

46. Collot, R., Ruiz-Moreno, C., Honhoff, C., van den Broek, T.J.M., Wezenaar, A.K.L., Kloosterman, D.J., Ariese, H.C.R., Johnson, H., Vervoort, B.M.T., Jeiroshi, A., et al. (2026). IGSF11-VISTA is a critical and targetable immune checkpoint axis in diffuse midline glioma. Cancer Cell 44, 641–657.e9.

47. Liu, X., Kwon, H., Li, Z., and Fu, Y.-X. (2017). Is CD47 an innate immune checkpoint for tumor evasion? J. Hematol. Oncol. 10, 12.

48. He, C., Sheng, L., Pan, D., Jiang, S., Ding, L., Ma, X., Liu, Y., and Jia, D. (2021). Single-cell transcriptomic analysis revealed a critical role of SPP1/CD44-mediated crosstalk between macrophages and cancer cells in glioma. Front. Cell Dev. Biol. 9, 779319.

49. Plate, K.H., Breier, G., Weich, H.A., and Risau, W. (1992). Vascular endothelial growth factor is a potential tumour angiogenesis factor in human gliomas in vivo. Nature 359, 845– 848.

50. Reardon, D.A., Turner, S., Peters, K.B., Desjardins, A., Gururangan, S., Sampson, J.H., McLendon, R.E., Herndon, J.E., 2nd, Jones, L.W., Kirkpatrick, J.P., et al. (2011). A review of VEGF/VEGFR-targeted therapeutics for recurrent glioblastoma. J. Natl. Compr. Canc. Netw. 9, 414–427.

51. Ribatti, D. (2022). Immunosuppressive effects of vascular endothelial growth factor. Oncol. Lett. 24, 369.

52. Cerezo-Wallis, D., Contreras-Alcalde, M., Troulé, K., Catena, X., Mucientes, C., Calvo, T.G., Cañón, E., Tejedo, C., Pennacchi, P.C., Hogan, S., et al. (2020). Midkine rewires the melanoma microenvironment toward a tolerogenic and immune-resistant state. Nat. Med. 26, 1865–1877.

53. Saikia, M., Prem, S., and Kapoor, V. (2026). Rewiring immunity: Midkine’s emerging role in cancer immune escape and drug resistance. iScience 29, 115497.

54. Zheng, Y., Jiang, H., Yang, N., Shen, S., Huang, D., Jia, L., Ling, J., Xu, L., Li, M., Yu, K., et al. (2024). Glioma-derived ANXA1 suppresses the immune response to TLR3 ligands by promoting an anti-inflammatory tumor microenvironment. Cell. Mol. Immunol. 21, 47–59.

55. Sun, Y., Senger, K., Baginski, T.K., Mazloom, A., Chinn, Y., Pantua, H., Hamidzadeh, K., Ramani, S.R., Luis, E., Tom, I., et al. (2012). Evolutionarily conserved paired immunoglobulin-like receptor α (PILRα) domain mediates its interaction with diverse sialylated ligands. J. Biol. Chem. 287, 15837–15850.

56. Ma, C., Chen, J., Ji, J., Zheng, Y., Liu, Y., Wang, J., Chen, T., Chen, H., Chen, Z., Zhou, Q., et al. (2024). Therapeutic modulation of APP-CD74 axis can activate phagocytosis of TAMs in GBM. Biochim. Biophys. Acta Mol. Basis Dis. 1870, 167449.

57. Taylor, K.R., Barron, T., Hui, A., Spitzer, A., Yalçin, B., Ivec, A.E., Geraghty, A.C., Hartmann, G.G., Arzt, M., Gillespie, S.M., et al. (2023). Glioma synapses recruit mechanisms of adaptive plasticity. Nature 623, 366–374.

58. Venkatesh, H.S., Johung, T.B., Caretti, V., Noll, A., Tang, Y., Nagaraja, S., Gibson, E.M., Mount, C.W., Polepalli, J., Mitra, S.S., et al. (2015). Neuronal activity promotes glioma growth through neuroligin-3 secretion. Cell 161, 803–816.

59. Hai, L., Zhang, C., Li, T., Zhou, X., Liu, B., Li, S., Zhu, M., Lin, Y., Yu, S., Zhang, K., et al. (2018). Notch1 is a prognostic factor that is distinctly activated in the classical and proneural subtype of glioblastoma and that promotes glioma cell survival via the NF-κB(p65) pathway. Cell Death Dis. 9, 158.

60. Bazzoni, R., and Bentivegna, A. (2019). Role of Notch signaling pathway in glioblastoma pathogenesis. Cancers (Basel) 11, E292.

61. Lachota, M., Zielniok, K., and Zagożdżon, R. (2026). Midkine and pleiotrophin in glioma: From mechanistic insights to therapeutic potential. Neoplasia 76, 101304.

62. Venkatesh, H.S., Morishita, W., Geraghty, A.C., Silverbush, D., Gillespie, S.M., Arzt, M., Tam, L.T., Espenel, C., Ponnuswami, A., Ni, L., et al. (2019). Electrical and synaptic integration of glioma into neural circuits. Nature 573, 539–545.

63. Wang, W., Li, T., Cheng, Y., Li, F., Qi, S., Mao, M., Wu, J., Liu, Q., Zhang, X., Li, X., et al. (2024). Identification of hypoxic macrophages in glioblastoma with therapeutic potential for vasculature normalization. Cancer Cell 42, 815–832.e12.

64. Puchalski, R.B., Shah, N., Miller, J., Dalley, R., Nomura, S.R., Yoon, J.-G., Smith, K.A., Lankerovich, M., Bertagnolli, D., Bickley, K., et al. (2018). An anatomic transcriptional atlas of human glioblastoma. Science 360, 660–663.

65. Chen, P., Huang, Y., Bong, R., Ding, Y., Song, N., Wang, X., Song, X., and Luo, Y. (2011). Tumor-associated macrophages promote angiogenesis and melanoma growth via adrenomedullin in a paracrine and autocrine manner. Clin. Cancer Res. 17, 7230–7239.

66. Zeng, Q., Li, S., Chepeha, D.B., Giordano, T.J., Li, J., Zhang, H., Polverini, P.J., Nor, J., Kitajewski, J., and Wang, C.-Y. (2005). Crosstalk between tumor and endothelial cells promotes tumor angiogenesis by MAPK activation of Notch signaling. Cancer Cell 8, 13– 23.

67. Domenga, V., Fardoux, P., Lacombe, P., Monet, M., Maciazek, J., Krebs, L.T., Klonjkowski, B., Berrou, E., Mericskay, M., Li, Z., et al. (2004). Notch3 is required for arterial identity and maturation of vascular smooth muscle cells. Genes Dev. 18, 2730–2735.

68. Henshall, T.L., Keller, A., He, L., Johansson, B.R., Wallgard, E., Raschperger, E., Mäe, M.A., Jin, S., Betsholtz, C., and Lendahl, U. (2015). Notch3 is necessary for blood vessel integrity in the central nervous system. Arterioscler. Thromb. Vasc. Biol. 35, 409–420.

69. Tefft, J.B., Bays, J.L., Lammers, A., Kim, S., Eyckmans, J., and Chen, C.S. (2022). Notch1 and Notch3 coordinate for pericyte-induced stabilization of vasculature. Am. J. Physiol. Cell Physiol. 322, C185–C196.

70. Browaeys, R., Saelens, W., and Saeys, Y. (2020). NicheNet: modeling intercellular communication by linking ligands to target genes. Nat. Methods 17, 159–162.

71. Schalck, A., Sakellariou-Thompson, D., Forget, M.-A., Sei, E., Hughes, T.G., Reuben, A., Bai, S., Hu, M., Kumar, T., Hurd, M.W., et al. (2022). Single-Cell Sequencing Reveals Trajectory of Tumor-Infiltrating Lymphocyte States in Pancreatic Cancer. Cancer Discov. 12, 2330–2349.

72. Anstee, J.E., Feehan, K.T., Opzoomer, J.W., Dean, I., Muller, H.P., Bahri, M., Cheung, T.S., Liakath-Ali, K., Liu, Z., Choy, D., et al. (2023). LYVE-1+ macrophages form a collaborative CCR5-dependent perivascular niche that influences chemotherapy responses in murine breast cancer. Dev. Cell 58, 1548–1561.e10.

73. Nalio Ramos, R., Missolo-Koussou, Y., Gerber-Ferder, Y., Bromley, C.P., Bugatti, M., Núñez, N.G., Tosello Boari, J., Richer, W., Menger, L., Denizeau, J., et al. (2022). Tissue-resident FOLR2+ macrophages associate with CD8+ T cell infiltration in human breast cancer. Cell 185, 1189–1207.e25.

74. Elfstrum, A.K., Rumahorbo, A.H., Reese, L.E., Nelson, E.V., McCluskey, B.M., and Schwertfeger, K.L. (2024). LYVE-1-expressing macrophages modulate the hyaluronan-containing extracellular matrix in the mammary stroma and contribute to mammary tumor growth. Cancer Res. Commun. 4, 1380–1397.

75. Siret, C., van Lessen, M., Bavais, J., Jeong, H.W., Reddy Samawar, S.K., Kapupara, K., Wang, S., Simic, M., de Fabritus, L., Tchoghandjian, A., et al. (2022). Deciphering the heterogeneity of the Lyve1+ perivascular macrophages in the mouse brain. Nat. Commun. 13, 7366.

76. Faraco, G., Park, L., Anrather, J., and Iadecola, C. (2017). Brain perivascular macrophages: characterization and functional roles in health and disease. J. Mol. Med. 95, 1143–1152.

77. Wen, W., Cheng, J., and Tang, Y. (2024). Brain perivascular macrophages: current understanding and future prospects. Brain 147, 39–55.

78. Alanio, C., Binder, Z.A., Chang, R.B., Nasrallah, M.P., Delman, D., Li, J.H., Tang, O.Y., Zhang, L.Y., Zhang, J.V., Wherry, E.J., et al. (2022). Immunologic features in DE Novo and recurrent Glioblastoma are associated with survival outcomes. Cancer Immunol. Res. 10, 800–810.

79. Mu, L., Yang, C., Gao, Q., Long, Y., Ge, H., DeLeon, G., Jin, L., Chang, Y.E., Sayour, E.J., Ji, J., et al. (2017). CD4+ and perivascular Foxp3+ T cells in glioma correlate with angiogenesis and tumor progression. Front. Immunol. 8, 1451.

80. Miller, T.E., El Farran, C.A., Couturier, C.P., Chen, Z., D’Antonio, J.P., Verga, J., Villanueva, M.A., Gonzalez Castro, L.N., Tong, Y.E., Saadi, T.A., et al. (2025). Programs, origins and immunomodulatory functions of myeloid cells in glioma. Nature, 1–11.

81. Sonpatki, P., Park, H.J., Xing, Y.L., Han, K.Y., Schroeder, B.A., Yu, H.J., Kim, H.J., Chowdhury, T., Hwang, J.H., Nam, S.M., et al. (2026). A spatially resolved human glioblastoma atlas reveals distinct cellular and molecular patterns of anatomical niches. Nat. Commun. 17, 2951.

82. Park, J.H., and Lee, H.K. (2022). Current understanding of hypoxia in glioblastoma multiforme and its response to immunotherapy. Cancers (Basel) 14, 1176.

83. Scharping, N.E., Menk, A.V., Whetstone, R.D., Zeng, X., and Delgoffe, G.M. (2017). Efficacy of PD-1 blockade is potentiated by metformin-induced reduction of tumor hypoxia. Cancer Immunol. Res. 5, 9–16.

84. Kim, A.-R., Choi, S.J., Park, J., Kwon, M., Chowdhury, T., Yu, H.J., Kim, S., Kang, H., Kim, K.-M., Park, S.-H., et al. (2022). Spatial immune heterogeneity of hypoxia-induced exhausted features in high-grade glioma. Oncoimmunology 11, 2026019.

85. Haley, M.J., Bere, L., Minshull, J., Georgaka, S., Garcia-Martin, N., Howell, G., Coope, D.J., Roncaroli, F., King, A., Wedge, D.C., et al. (2024). Hypoxia coordinates the spatial landscape of myeloid cells within glioblastoma to affect survival. Sci Adv 10, eadj3301.

86. Sattiraju, A., Kang, S., Giotti, B., Chen, Z., Marallano, V.J., Brusco, C., Ramakrishnan, A., Shen, L., Tsankov, A.M., Hambardzumyan, D., et al. (2023). Hypoxic niches attract and sequester tumor-associated macrophages and cytotoxic T cells and reprogram them for immunosuppression. Immunity 56, 1825–1843.e6.

87. Cannon, M., Stevenson, J., Stahl, K., Basu, R., Coffman, A., Kiwala, S., McMichael, J.F., Kuzma, K., Morrissey, D., Cotto, K., et al. (2024). DGIdb 5.0: rebuilding the drug-gene interaction database for precision medicine and drug discovery platforms. Nucleic Acids Res. 52, D1227–D1235.

88. Gilbert, M.R., Dignam, J.J., Armstrong, T.S., Wefel, J.S., Blumenthal, D.T., Vogelbaum, M.A., Colman, H., Chakravarti, A., Pugh, S., Won, M., et al. (2014). A randomized trial of bevacizumab for newly diagnosed glioblastoma. N. Engl. J. Med. 370, 699–708.

89. Fu, M., Zhou, Z., Huang, X., Chen, Z., Zhang, L., Zhang, J., Hua, W., and Mao, Y. (2023). Use of Bevacizumab in recurrent glioblastoma: a scoping review and evidence map. BMC Cancer 23, 544.

90. Butler, A., Hoffman, P., Smibert, P., Papalexi, E., and Satija, R. (2018). Integrating single-cell transcriptomic data across different conditions, technologies, and species. Nat. Biotechnol. 36, 411–420.

91. Tirosh, I., Izar, B., Prakadan, S.M., Wadsworth, M.H., 2nd, Treacy, D., Trombetta, J.J., Rotem, A., Rodman, C., Lian, C., Murphy, G., et al. (2016). Dissecting the multicellular ecosystem of metastatic melanoma by single-cell RNA-seq. Science 352, 189–196.

92. Yu, G., Wang, L.-G., Han, Y., and He, Q.-Y. (2012). clusterProfiler: an R package for comparing biological themes among gene clusters. OMICS 16, 284–287.

93. Xu, S., Hu, E., Cai, Y., Xie, Z., Luo, X., Zhan, L., Tang, W., Wang, Q., Liu, B., Wang, R., et al. (2024). Using clusterProfiler to characterize multiomics data. Nat. Protoc. 19, 3292– 3320.

94. Zheng, G.X.Y., Terry, J.M., Belgrader, P., Ryvkin, P., Bent, Z.W., Wilson, R., Ziraldo, S.B., Wheeler, T.D., McDermott, G.P., Zhu, J., et al. (2017). Massively parallel digital transcriptional profiling of single cells. Nat Commun 8, 14049.

95. Lun, A.T.L., McCarthy, D.J., and Marioni, J.C. (2016). A step-by-step workflow for low-level analysis of single-cell RNA-seq data with Bioconductor. F1000Res. 5, 2122.

96. Bais, A.S., and Kostka, D. (2019). scds: Computational Annotation of Doublets in Single Cell RNA Sequencing Data. bioRxiv, 564021. 10.1101/564021.

97. Germain, P.-L., Lun, A., Garcia Meixide, C., Macnair, W., and Robinson, M.D. (2021). Doublet identification in single-cell sequencing data using *scDblFinder*. F1000Res. 10, 979.

98. McGinnis, C.S., Murrow, L.M., and Gartner, Z.J. (2019). DoubletFinder: Doublet Detection in Single-Cell RNA Sequencing Data Using Artificial Nearest Neighbors. Cell Syst 8, 329–337.e4.

99. Hao, Y., Hao, S., Andersen-Nissen, E., Mauck, W.M., 3rd, Zheng, S., Butler, A., Lee, M.J., Wilk, A.J., Darby, C., Zager, M., et al. (2021). Integrated analysis of multimodal single-cell data. Cell 184, 3573–3587.e29.

100. Aran, D., Looney, A.P., Liu, L., Wu, E., Fong, V., Hsu, A., Chak, S., Naikawadi, R.P., Wolters, P.J., Abate, A.R., et al. (2019). Reference-based analysis of lung single-cell sequencing reveals a transitional profibrotic macrophage. Nat. Immunol. 20, 163–172.

101. Guo, H., and Li, J. (2021). scSorter: assigning cells to known cell types according to marker genes. Genome Biol. 22, 69.

102. Pombo Antunes, A.R., Scheyltjens, I., Lodi, F., Messiaen, J., Antoranz, A., Duerinck, J., Kancheva, D., Martens, L., De Vlaminck, K., Van Hove, H., et al. (2021). Single-cell profiling of myeloid cells in glioblastoma across species and disease stage reveals macrophage competition and specialization. Nat. Neurosci. 24, 595–610.

103. Song, L., Pan, S., Zhang, Z., Jia, L., Chen, W.-H., and Zhao, X.-M. (2021). STAB: a spatio-temporal cell atlas of the human brain. Nucleic Acids Res. 49, D1029–D1037.

104. Müller, S., Cho, A., Liu, S.J., Lim, D.A., and Diaz, A. (2018). CONICS integrates scRNA-seq with DNA sequencing to map gene expression to tumor sub-clones. Bioinformatics 34, 3217–3219.

105. Gu, Z., Schlesner, M., and Hübschmann, D. (2021). cola: an R/Bioconductor package for consensus partitioning through a general framework. Nucleic Acids Res. 49, e15.

106. Jin, S., Guerrero-Juarez, C.F., Zhang, L., Chang, I., Ramos, R., Kuan, C.-H., Myung, P., Plikus, M.V., and Nie, Q. (2021). Inference and analysis of cell-cell communication using CellChat. Nat. Commun. 12, 1088.

107. Xun, Z., Ding, X., Zhang, Y., Zhang, B., Lai, S., Zou, D., Zheng, J., Chen, G., Su, B., Han, L., et al. (2023). Reconstruction of the tumor spatial microenvironment along the malignant-boundary-nonmalignant axis. Nat. Commun. 14, 933.

108. Pham, D., Tan, X., Balderson, B., Xu, J., Grice, L.F., Yoon, S., Willis, E.F., Tran, M., Lam, P.Y., Raghubar, A., et al. (2023). Robust mapping of spatiotemporal trajectories and cell-cell interactions in healthy and diseased tissues. Nat. Commun. 14, 7739.

109. Jin, S., Plikus, M.V., and Nie, Q. (2025). CellChat for systematic analysis of cell-cell communication from single-cell transcriptomics. Nat. Protoc. 20, 180–219.

110. Cang, Z., Zhao, Y., Almet, A.A., Stabell, A., Ramos, R., Plikus, M.V., Atwood, S.X., and Nie, Q. (2023). Screening cell-cell communication in spatial transcriptomics via collective optimal transport. Nat. Methods 20, 218–228.

111. Csárdi, G., and Nepusz, T. (2006). The igraph software package for complex network research. 1695.

