## Supplementary figures and images for "Mapping the spatial architecture of glioblastoma from core to edge delineates niche-specific tumor cell states and intercellular interactions"

### Fig. S1

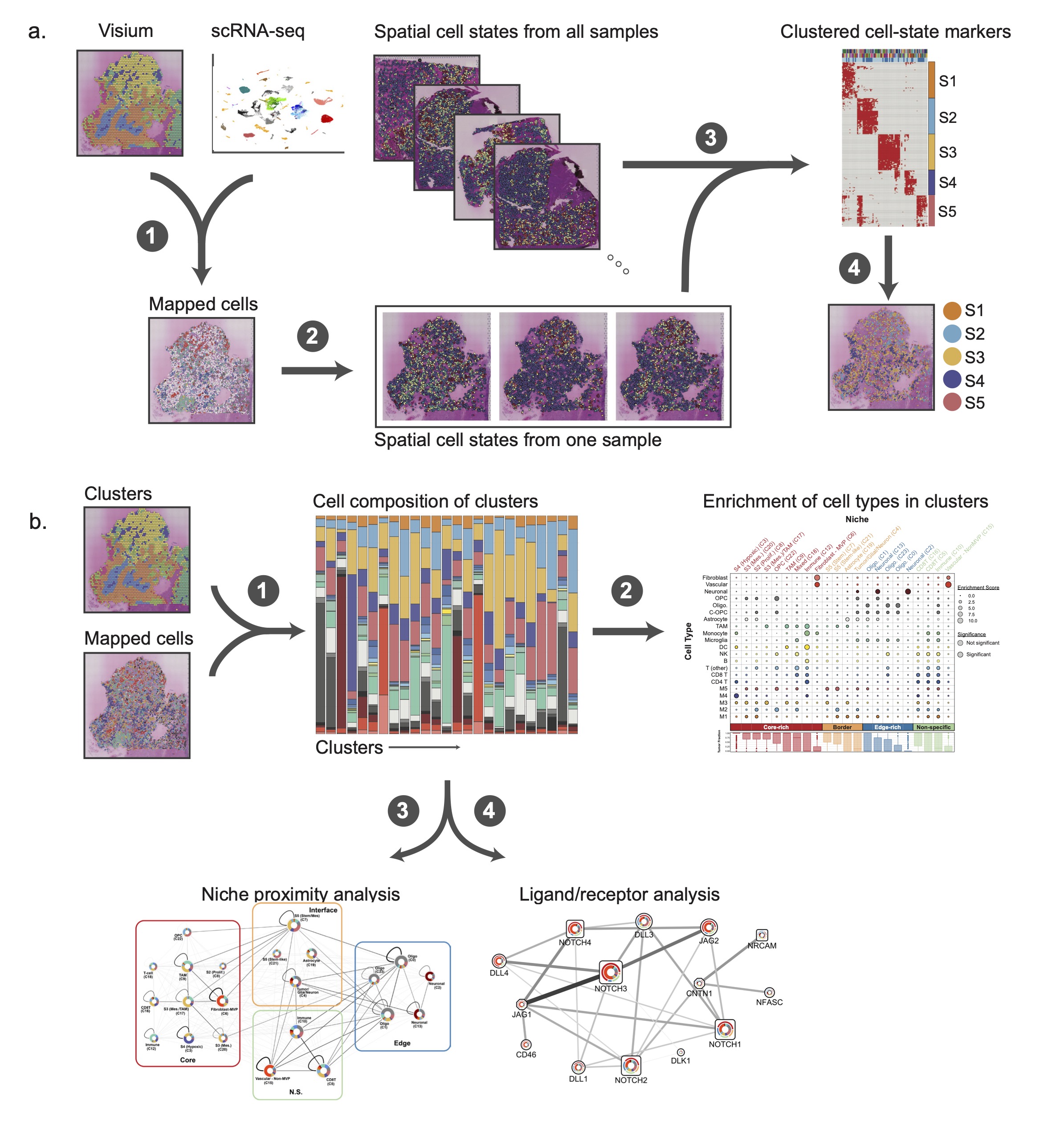

### Fig. S2

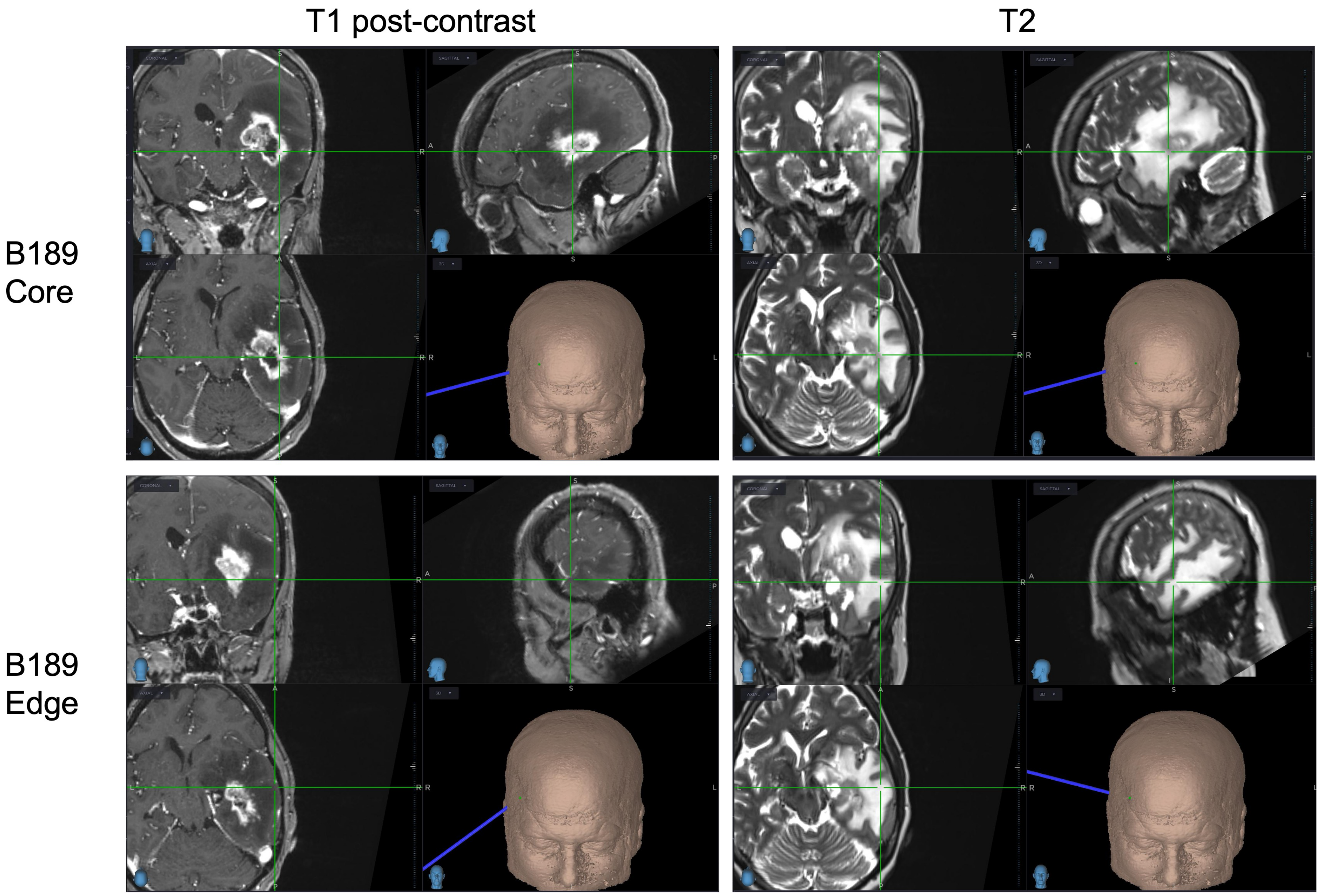

### Fig. S3

#### Visium OCT samples

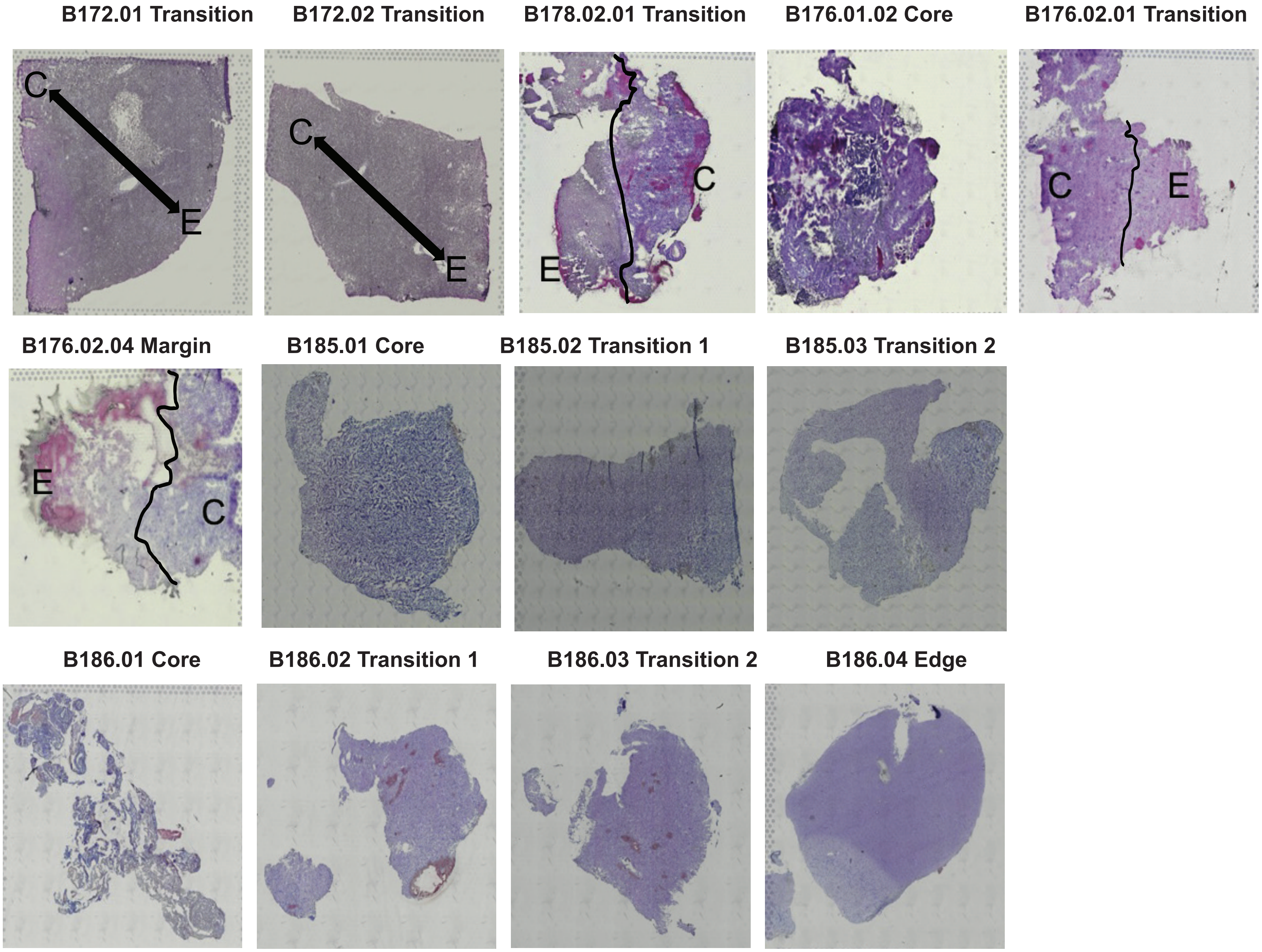

### Fig. S5

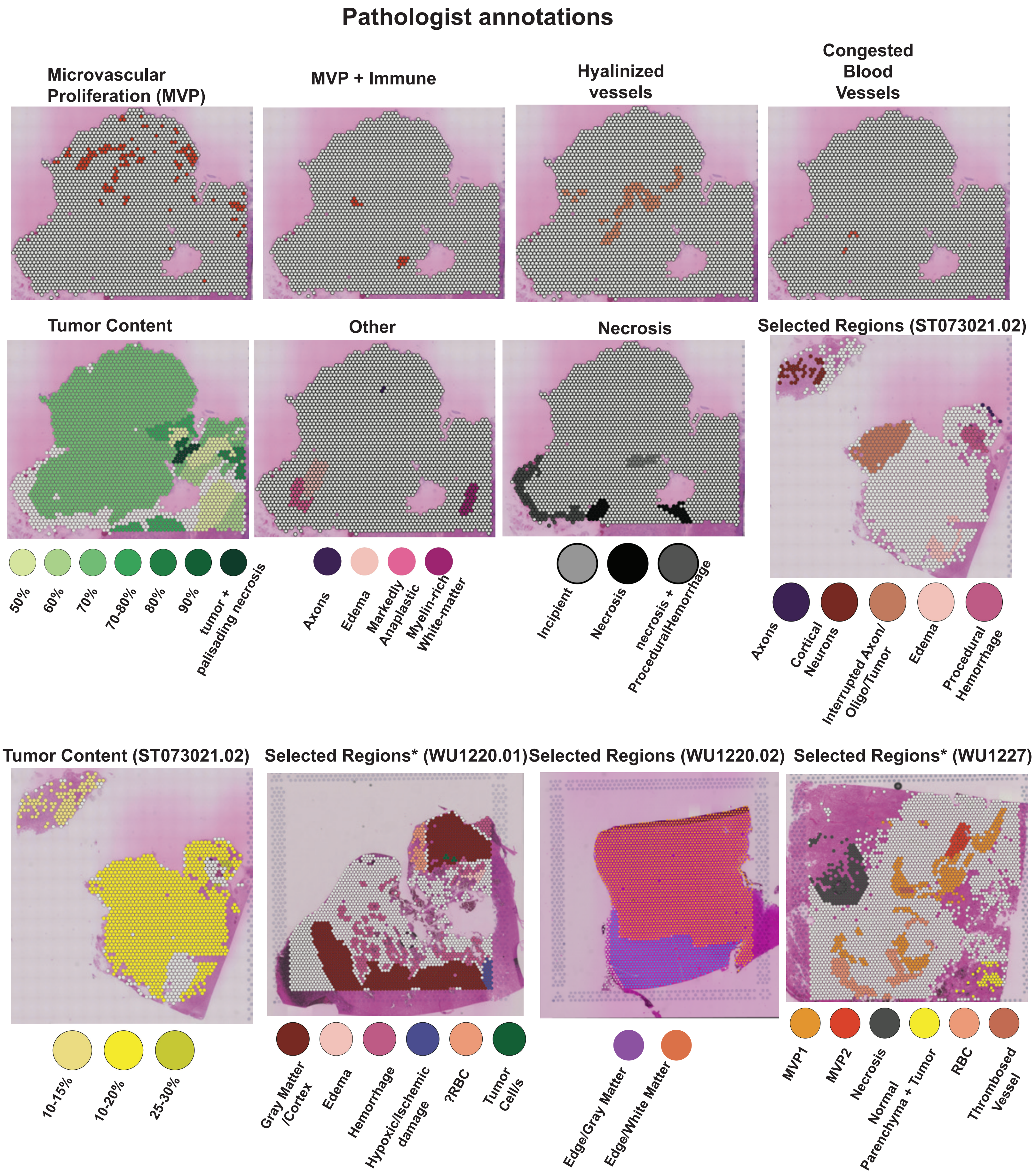


*Spots not otherwise annotated are tumor

### Fig. S6

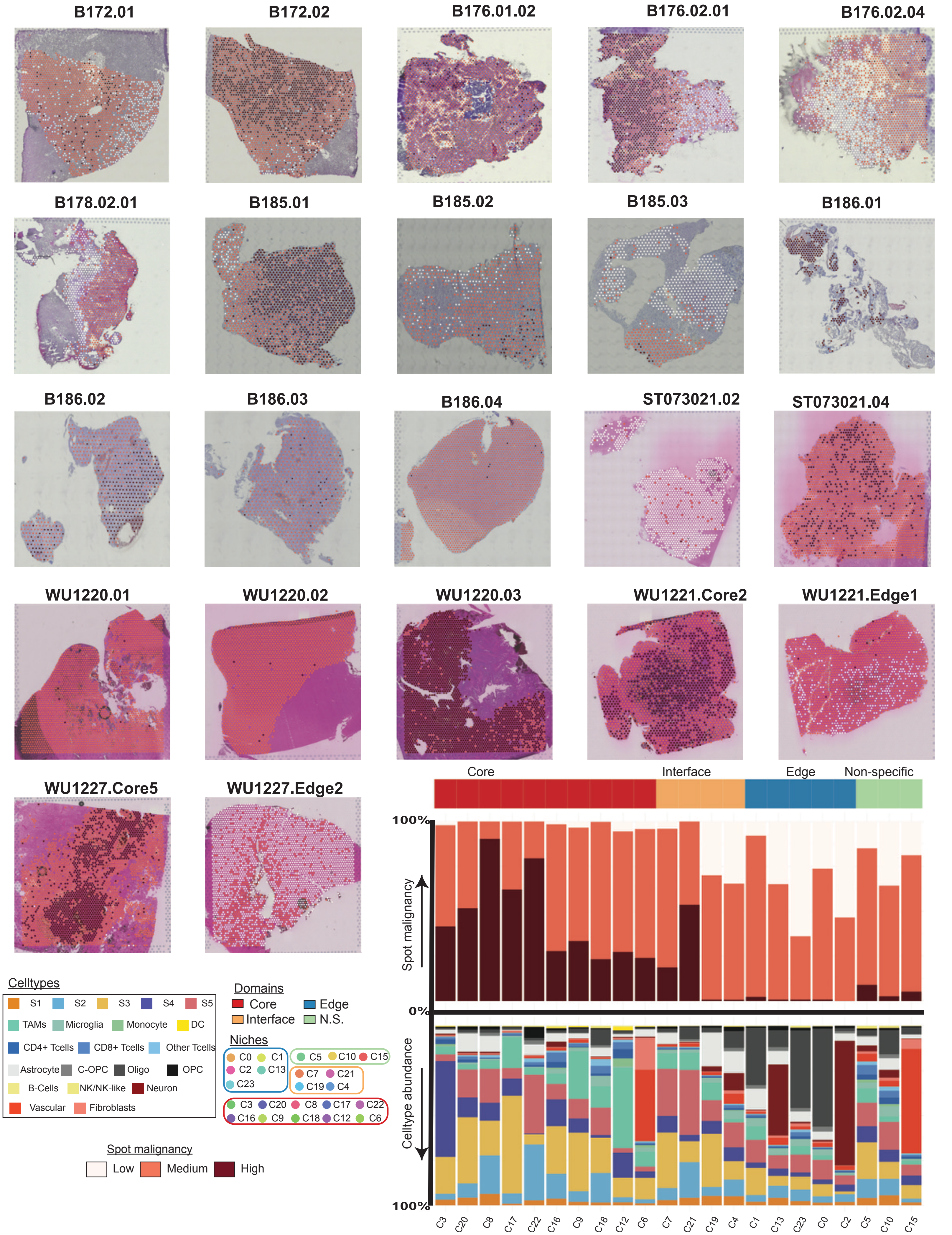

### Fig. S7

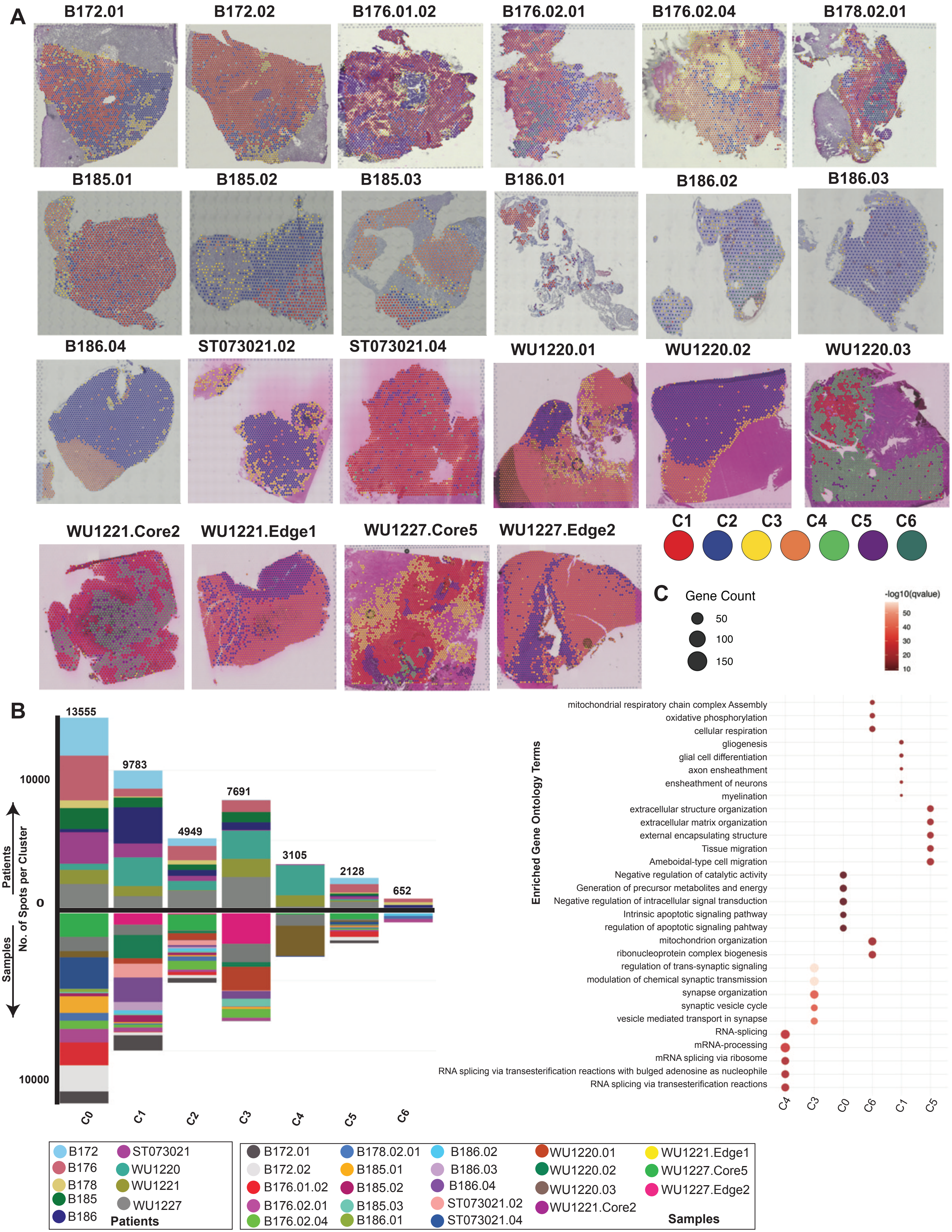

### Fig. S8

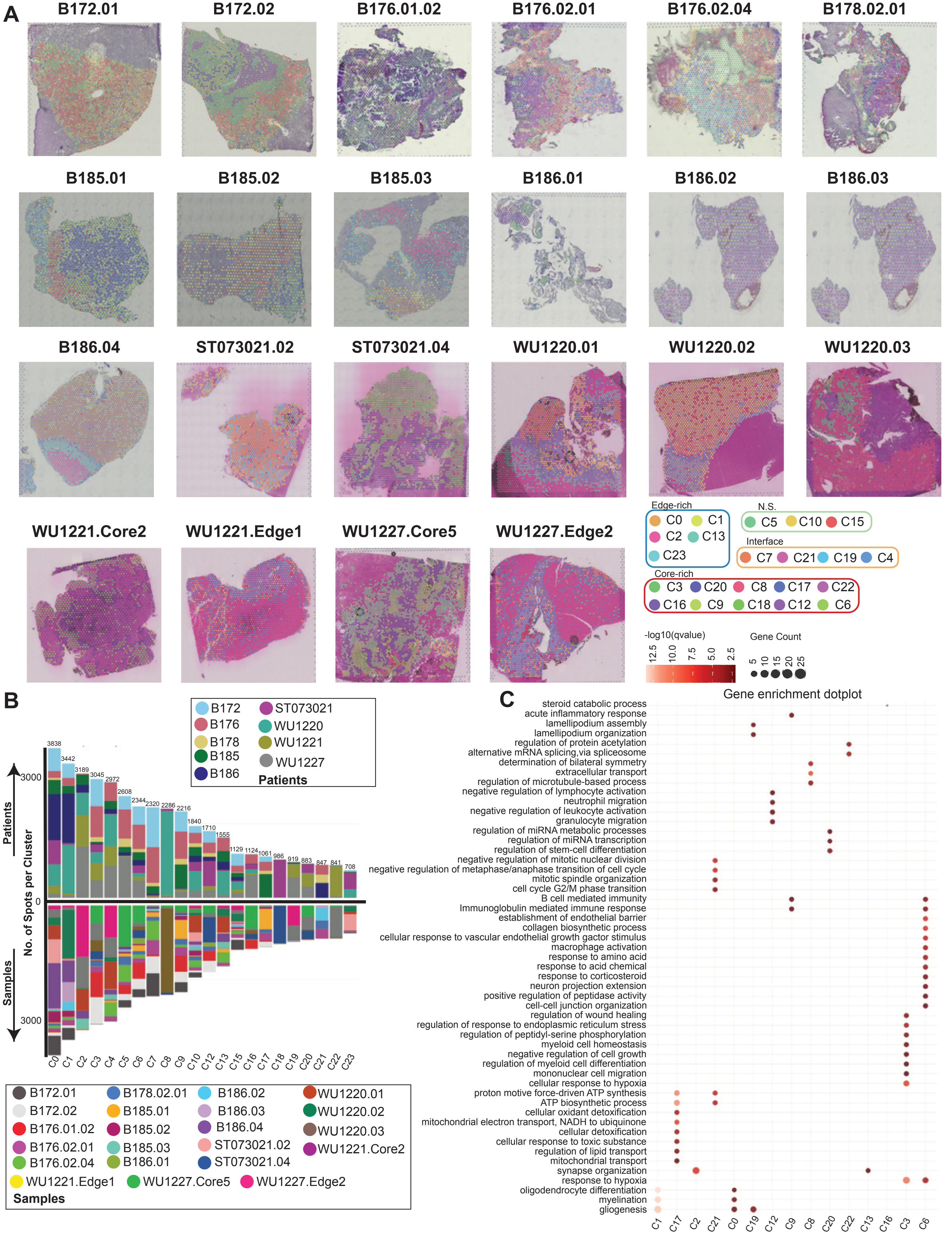

### Fig. S9

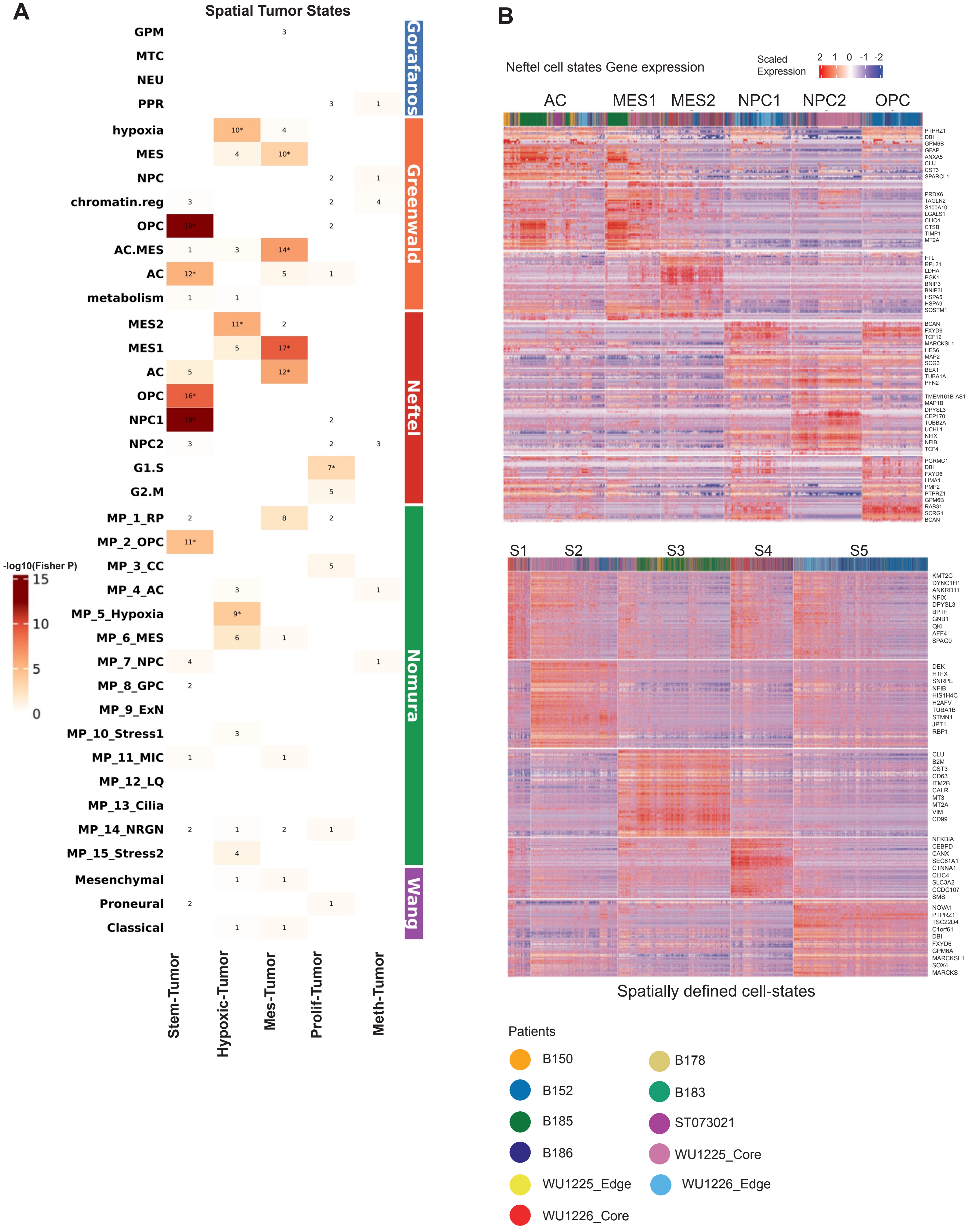

### Fig. S11

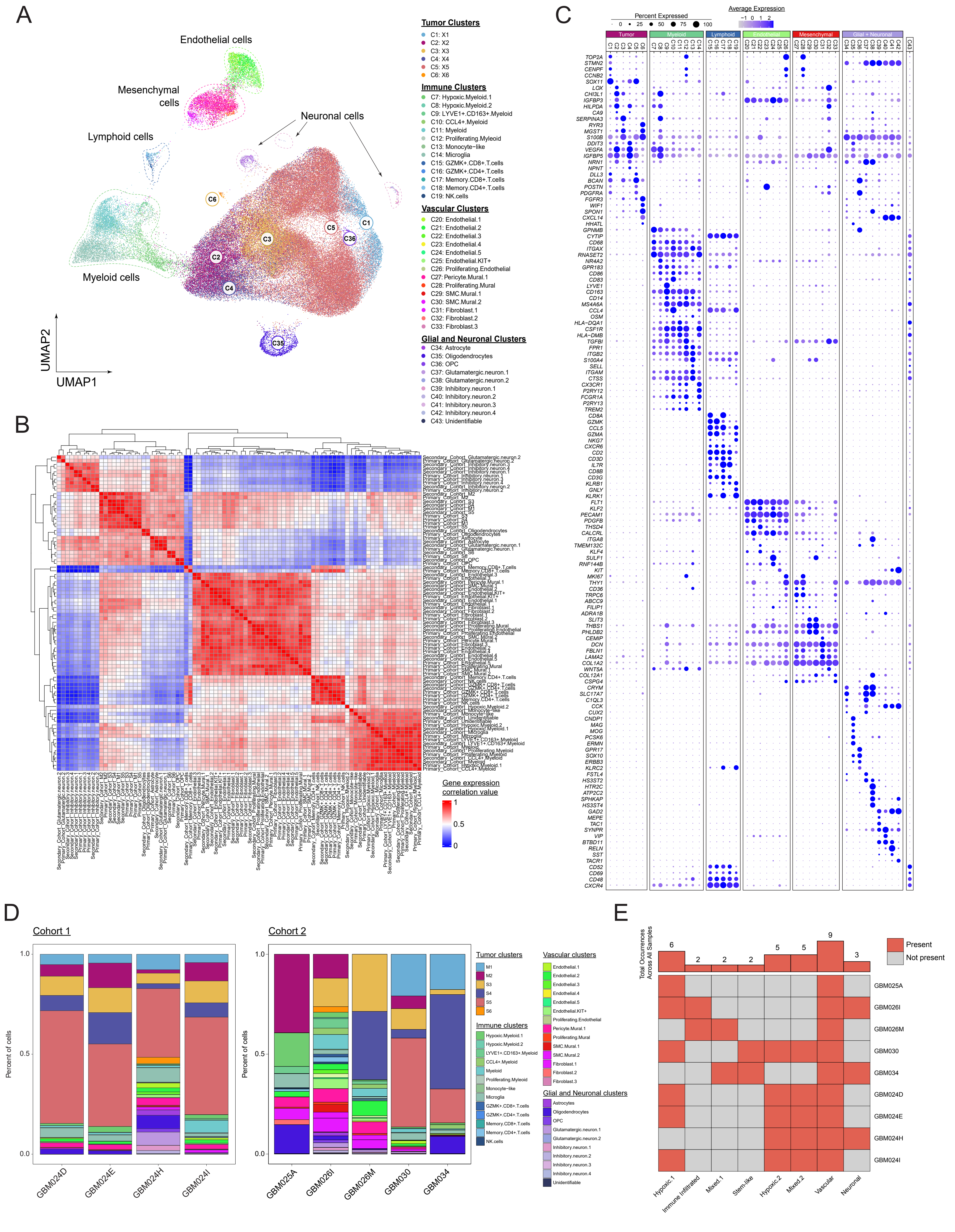

### Fig. S12

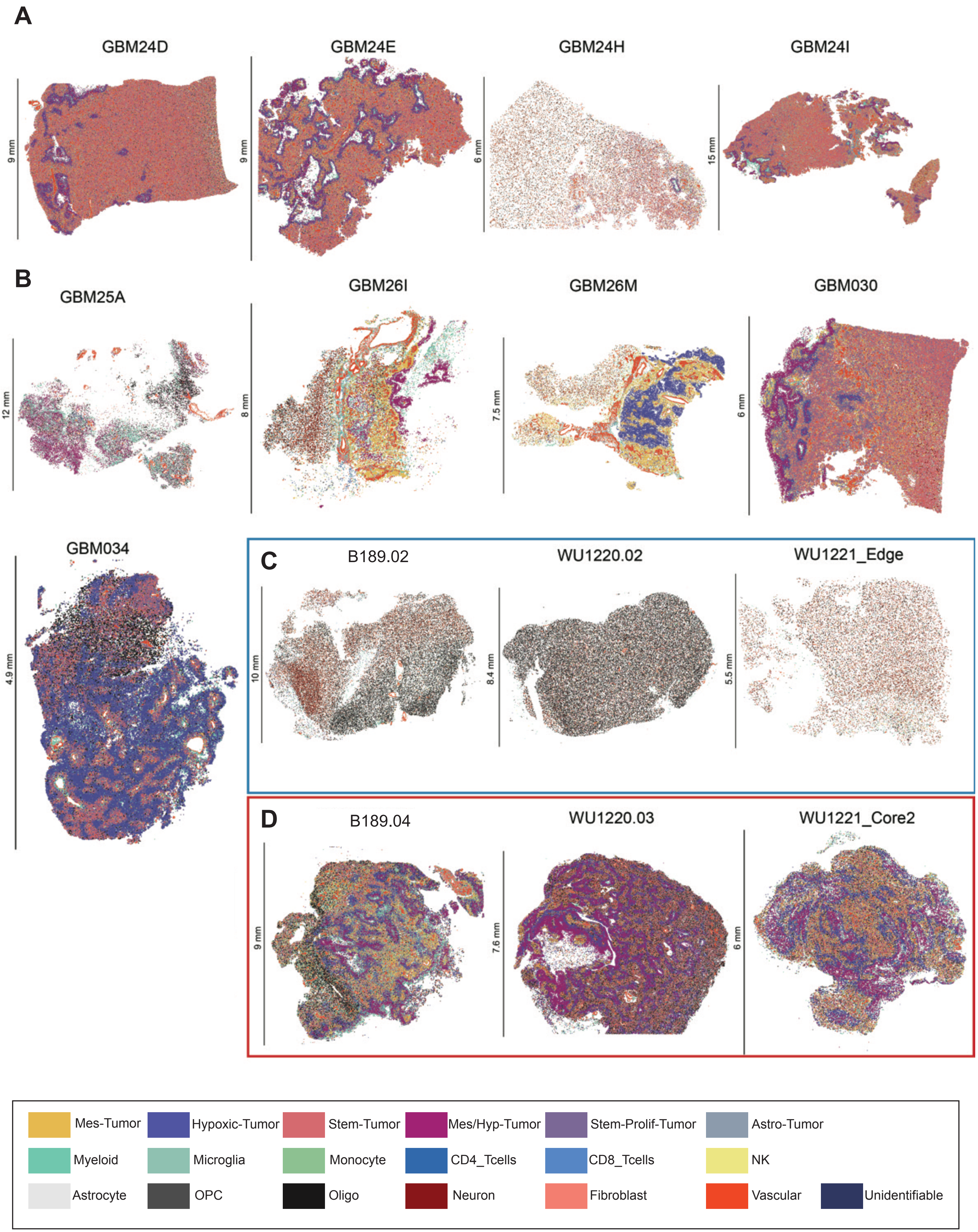

### Fig. S13

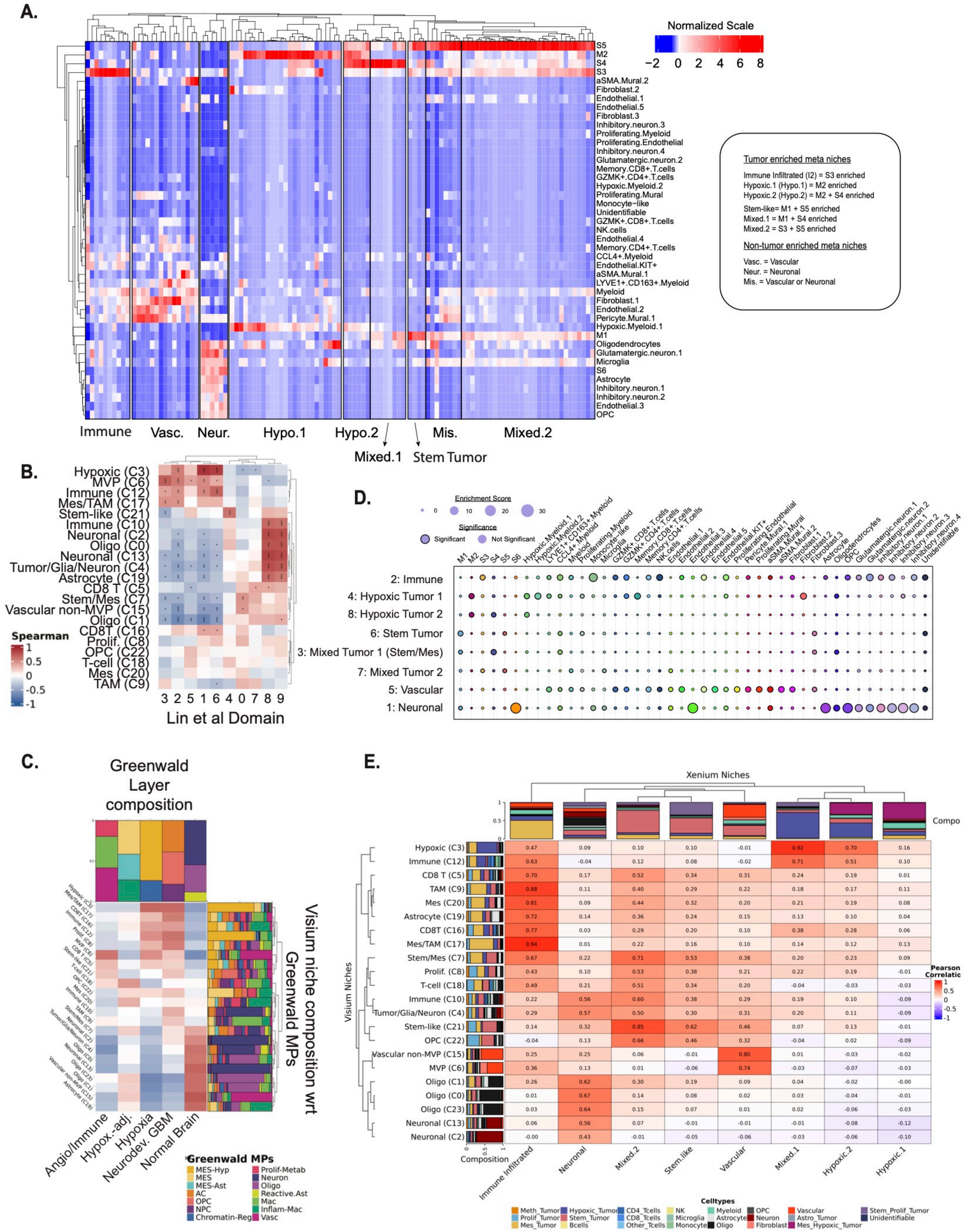

### Fig. S14

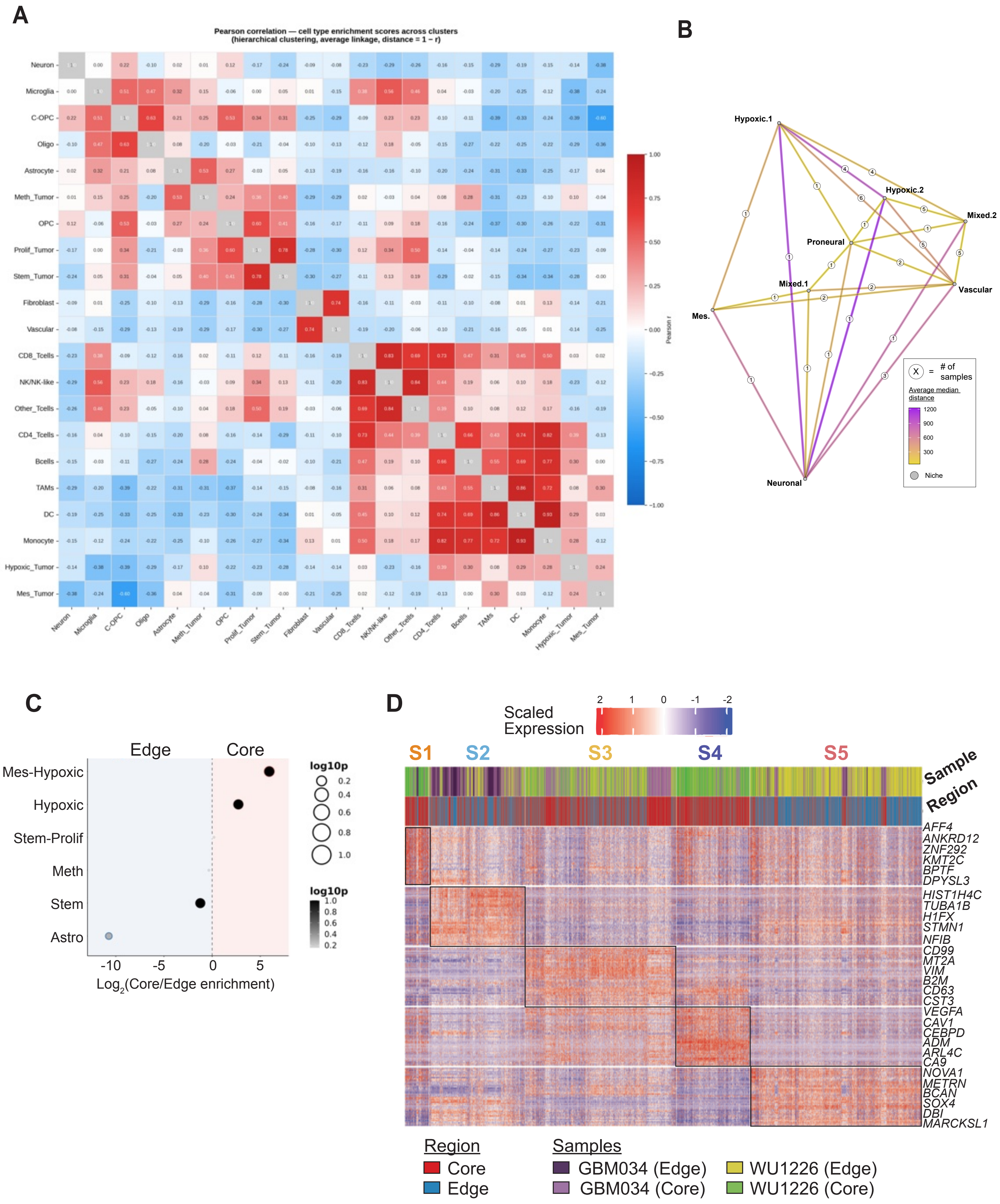

### Fig. S15

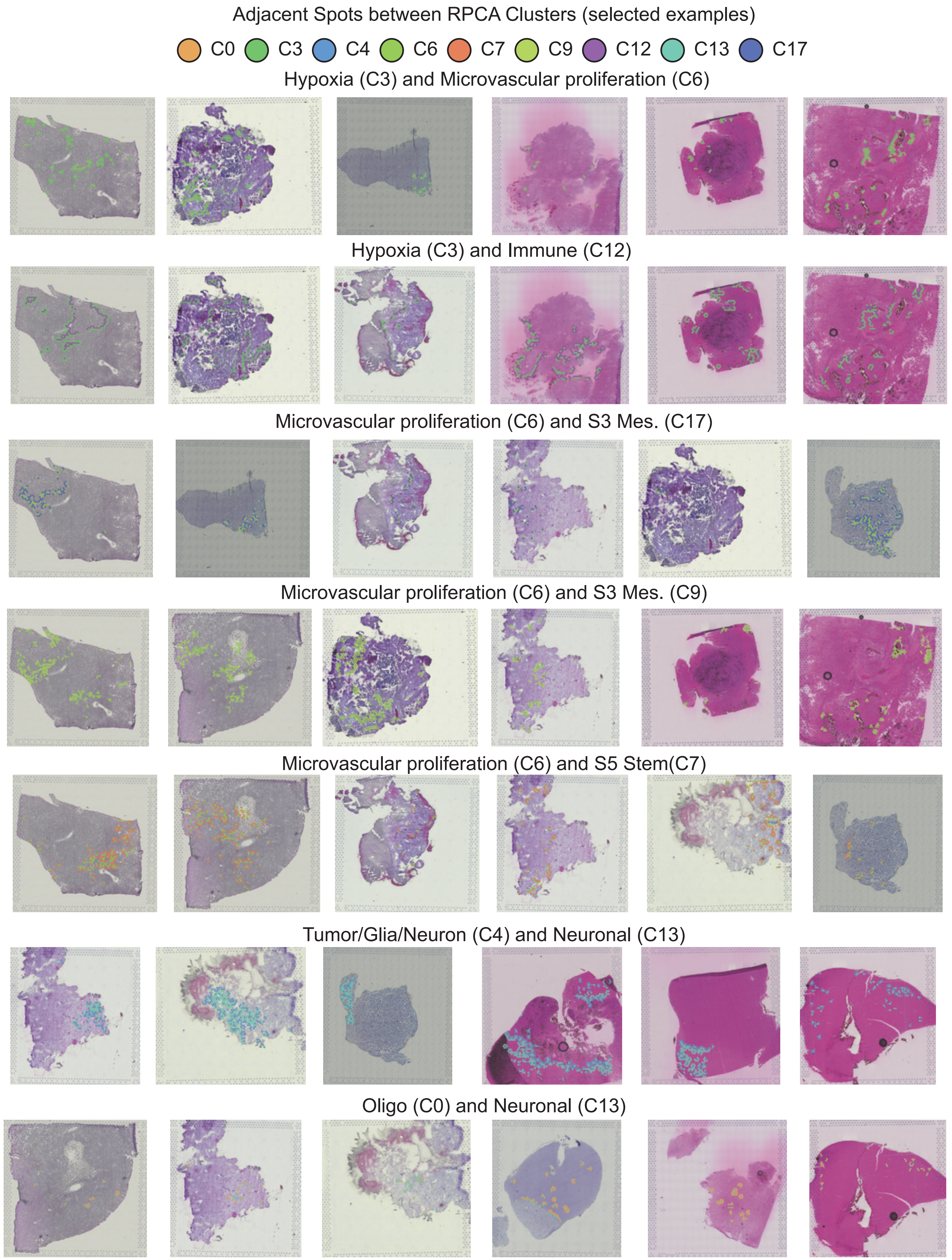

### Fig. S16

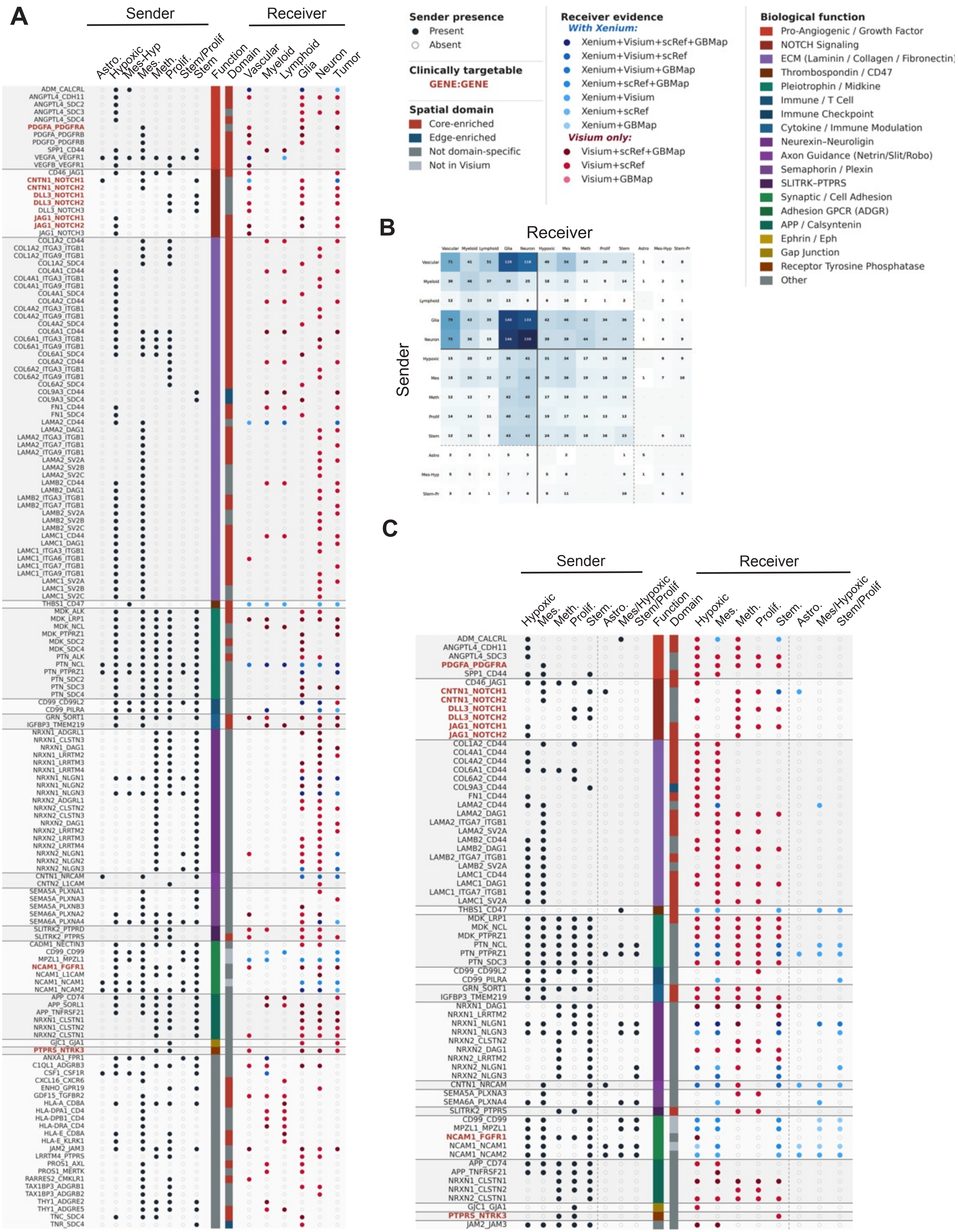

### Fig. S17

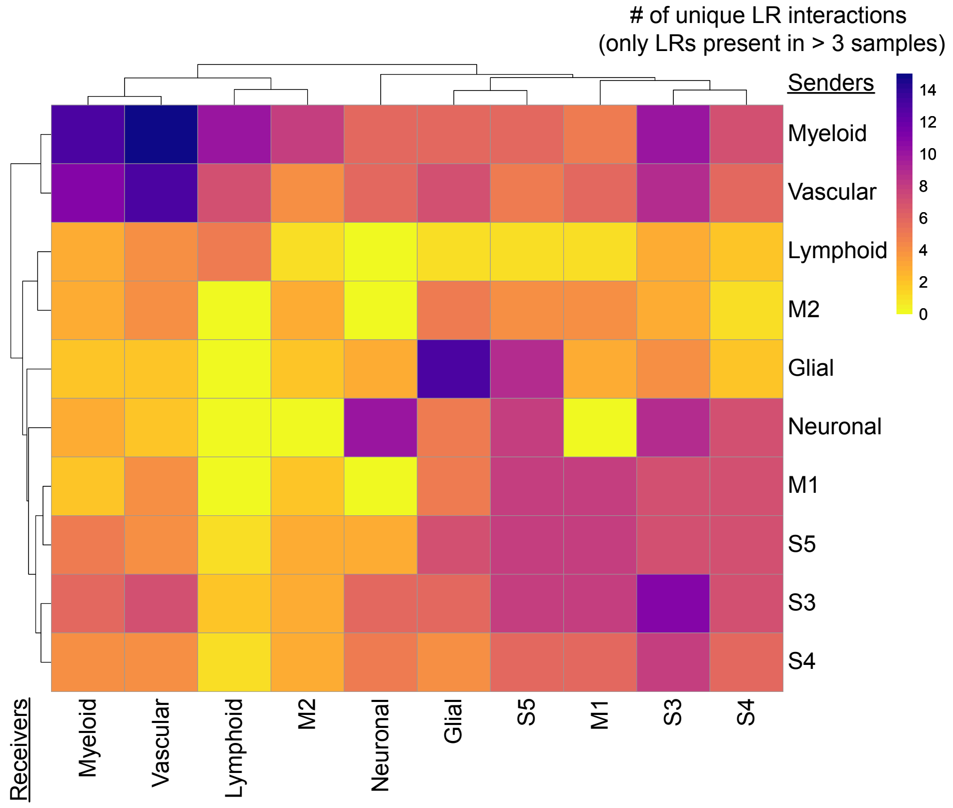

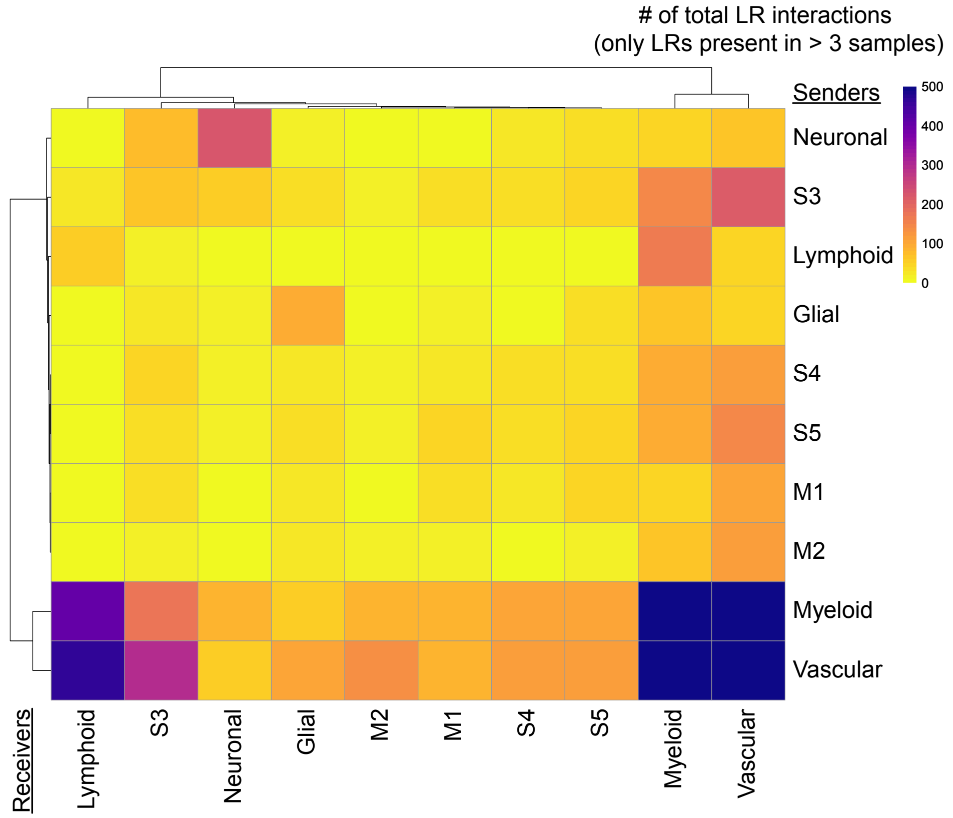

### Fig. S18

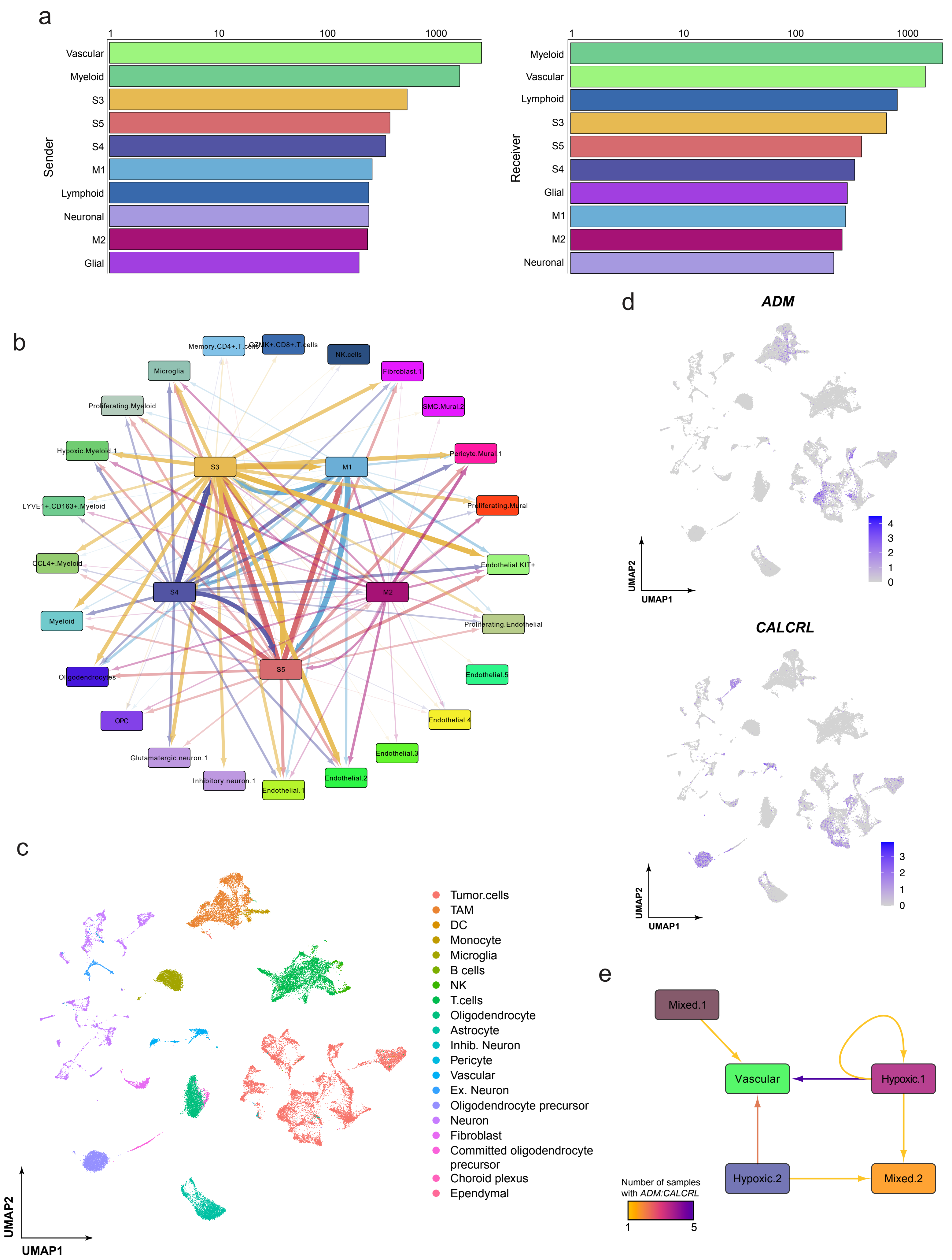

### Fig. S19

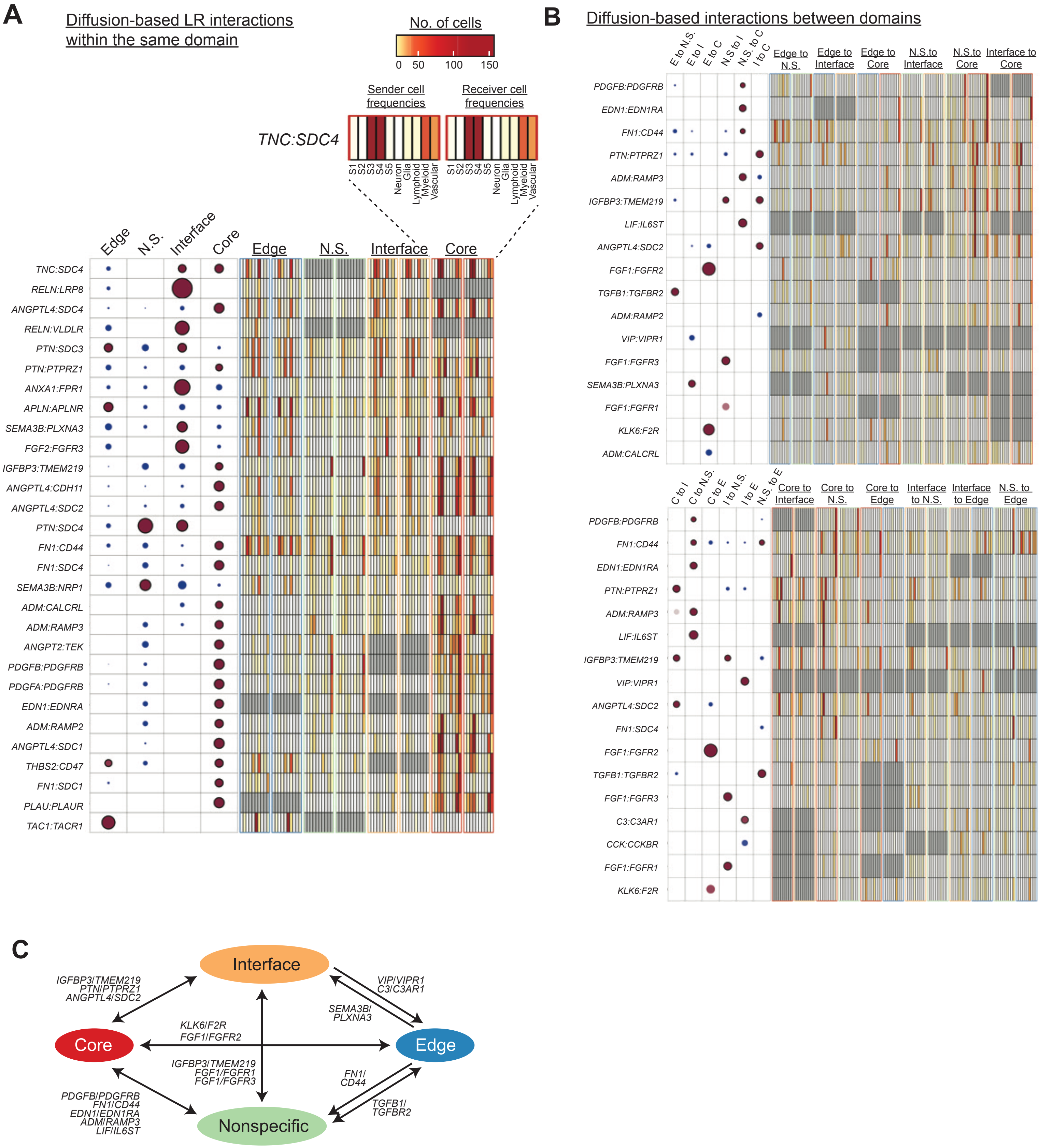

### Fig. S20

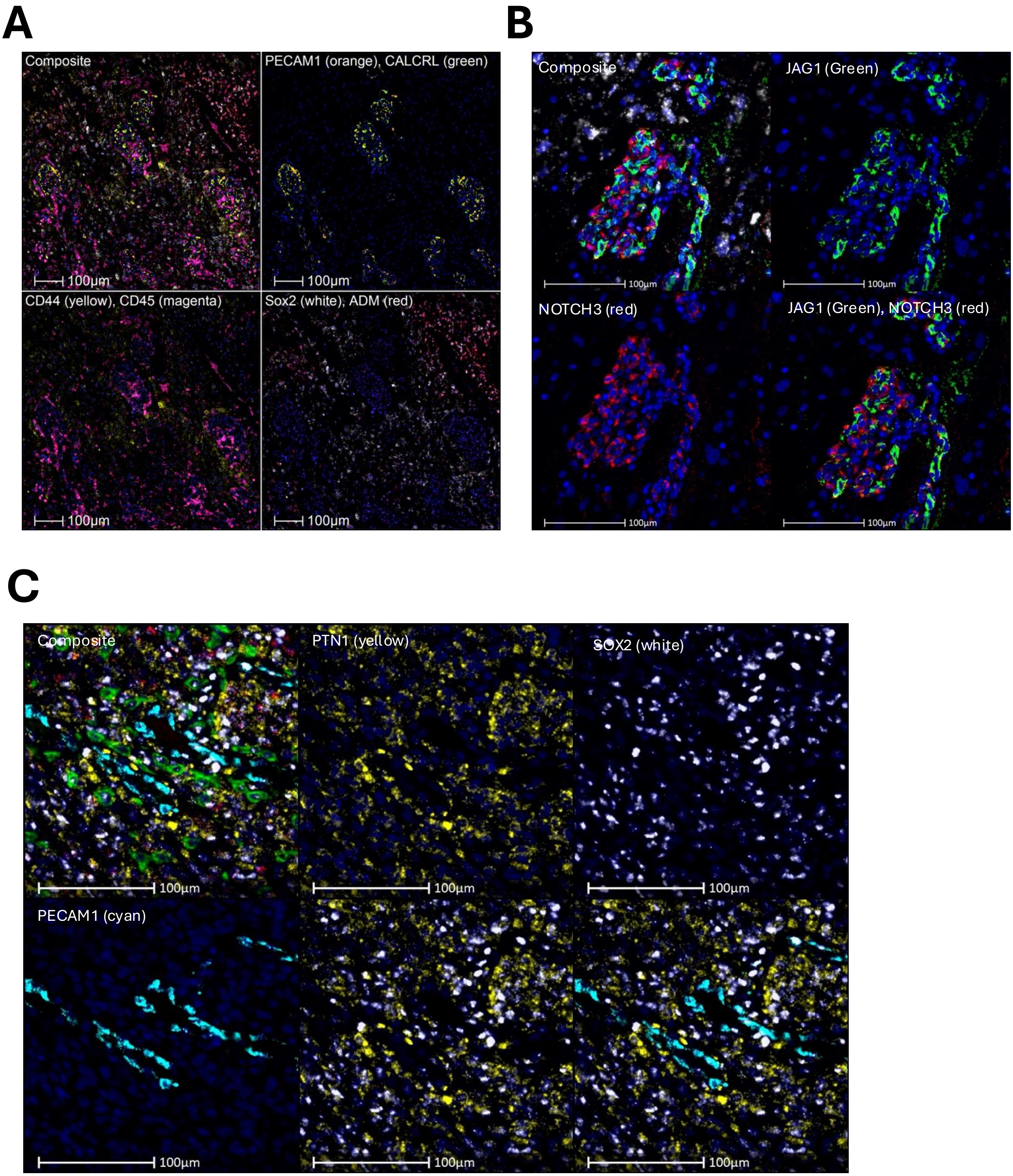

### Fig. S21

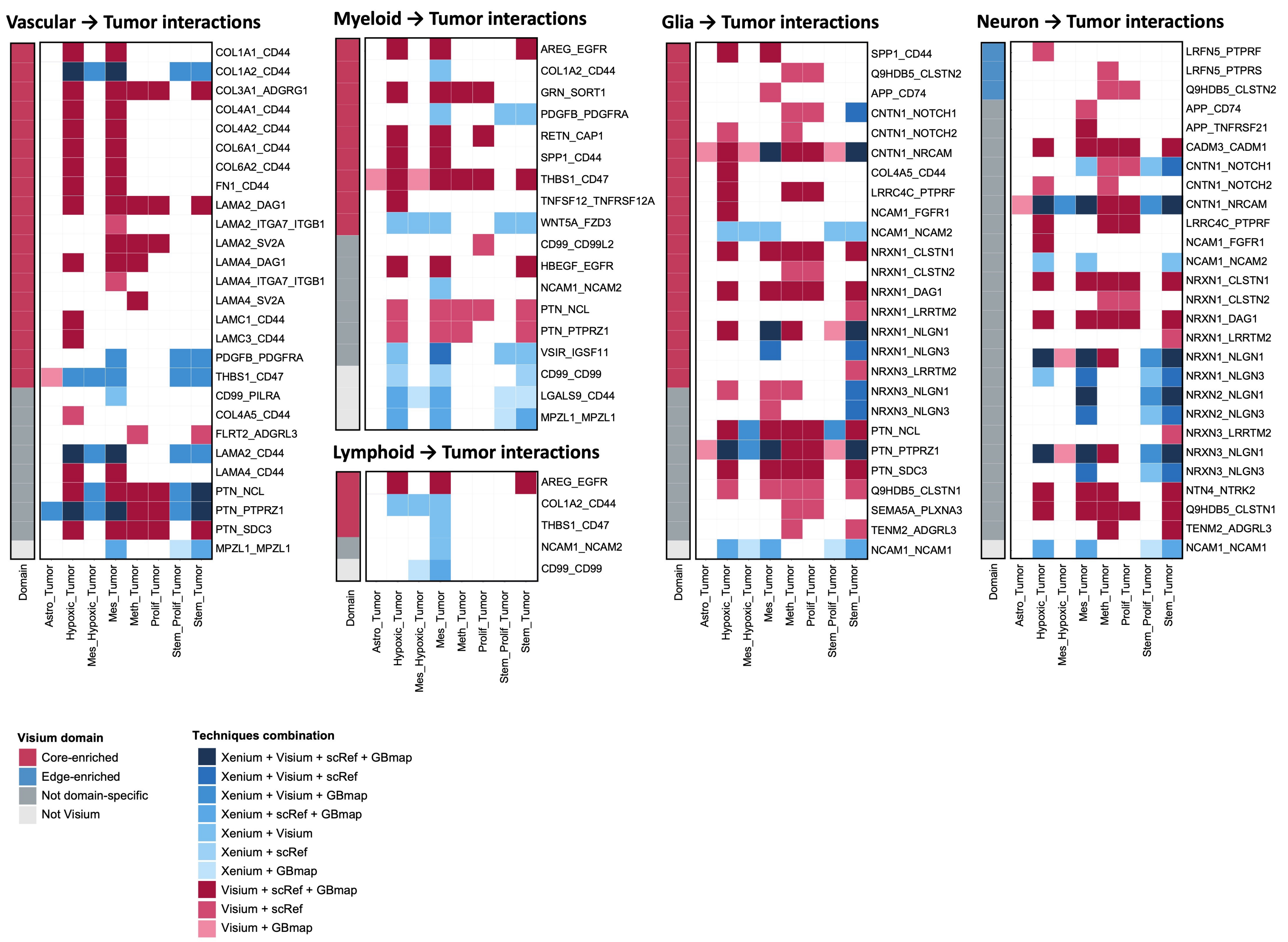

### Fig. S22

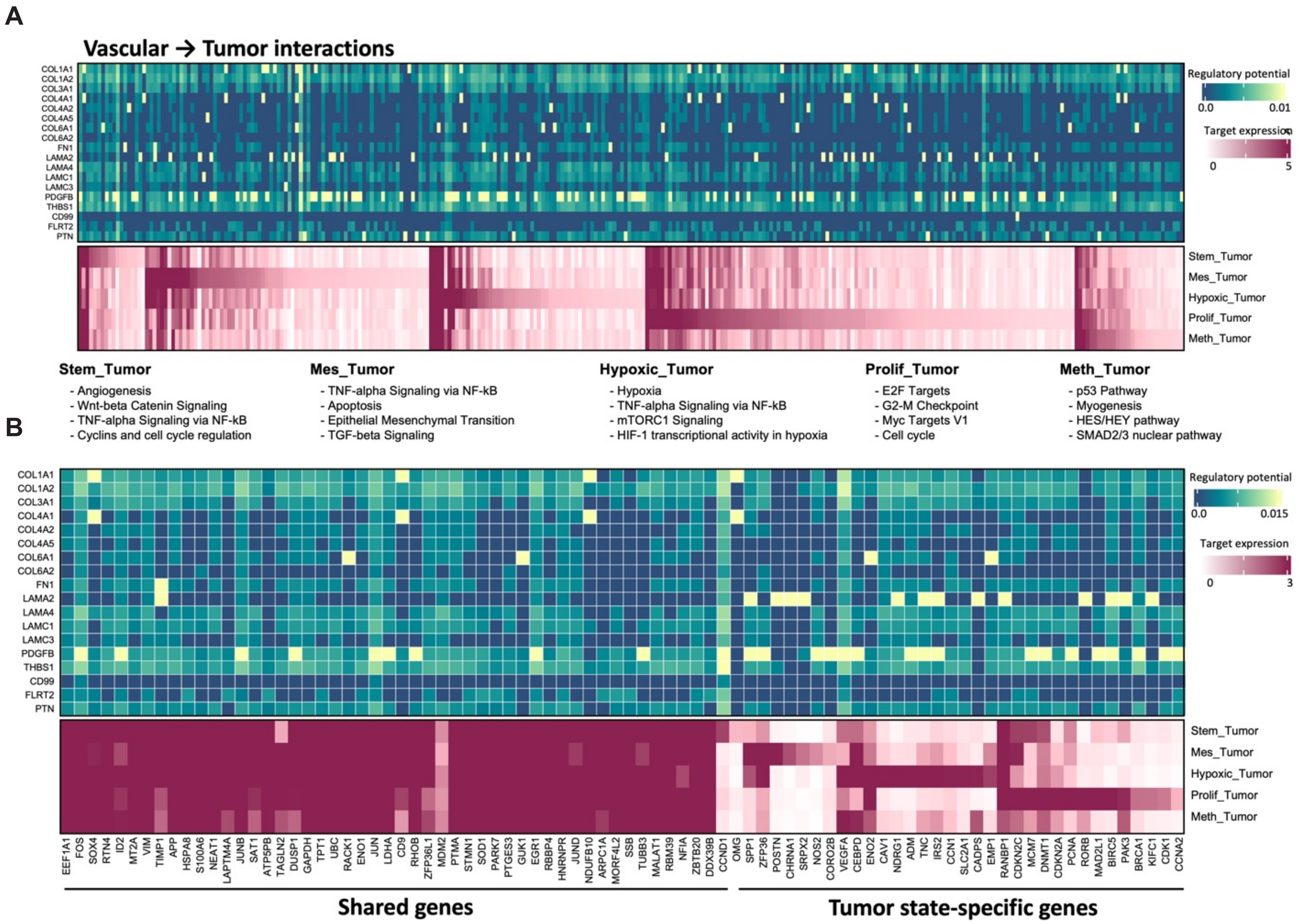

### Fig. S23

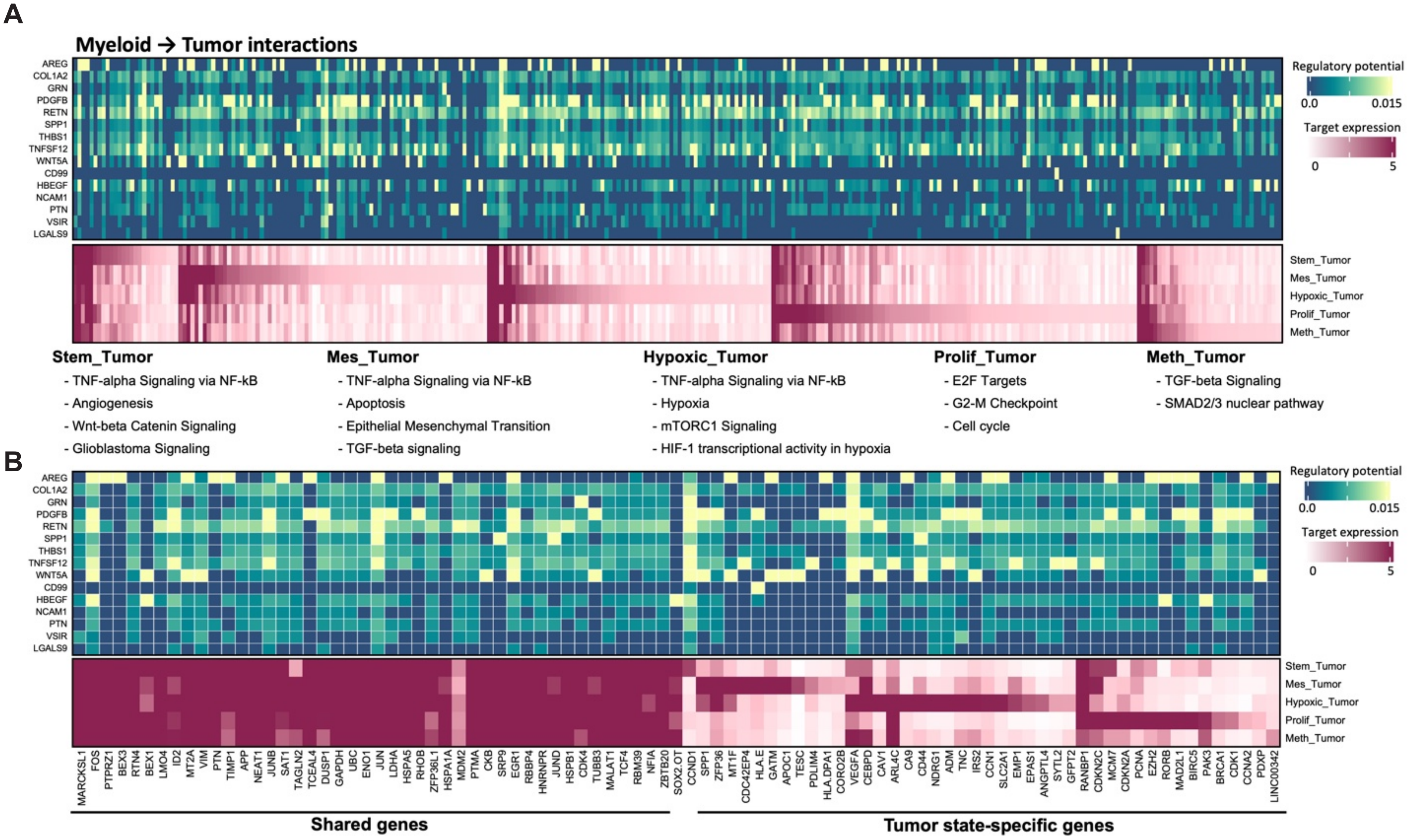

### Fig. S24

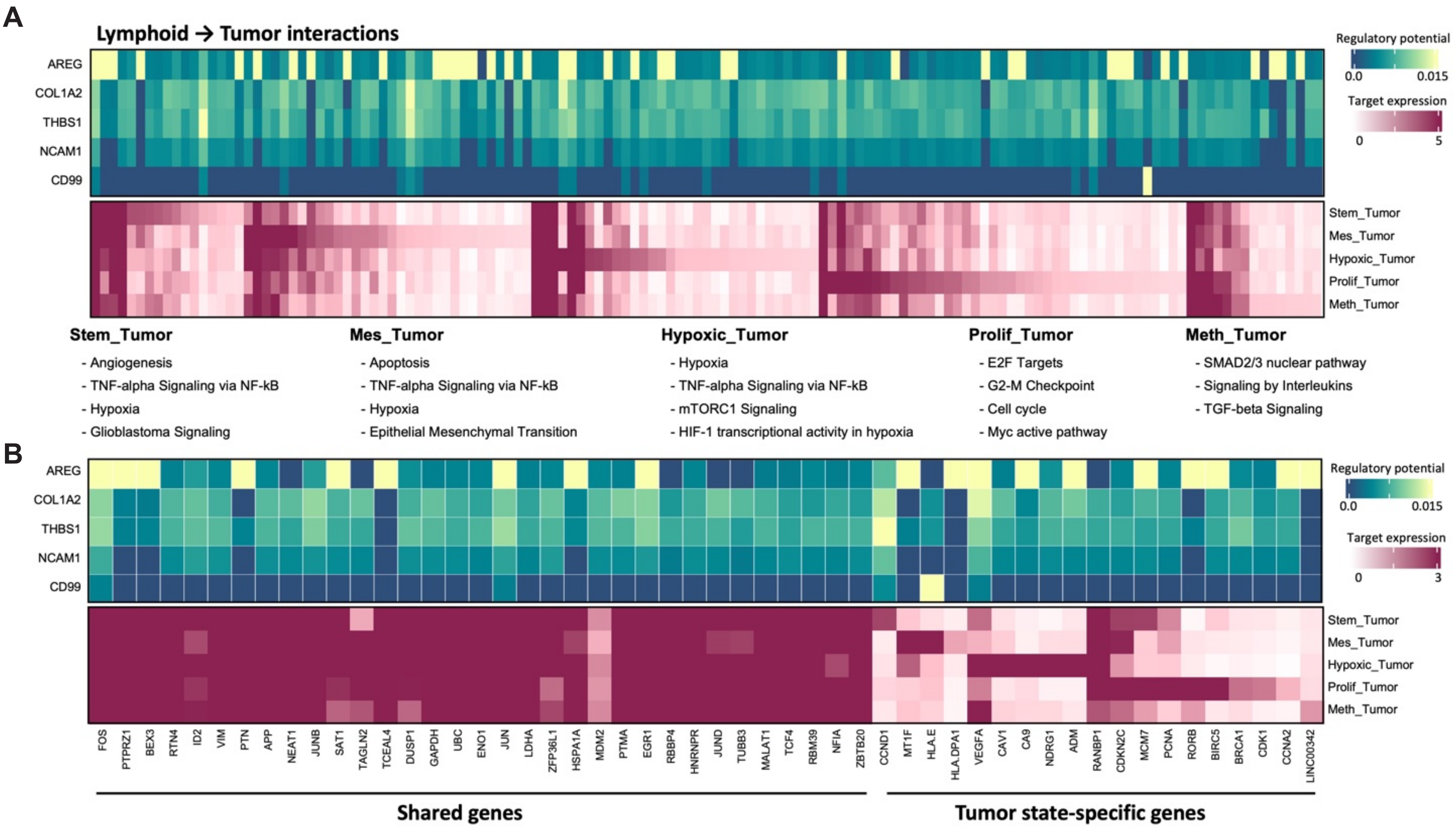

### Fig. S25

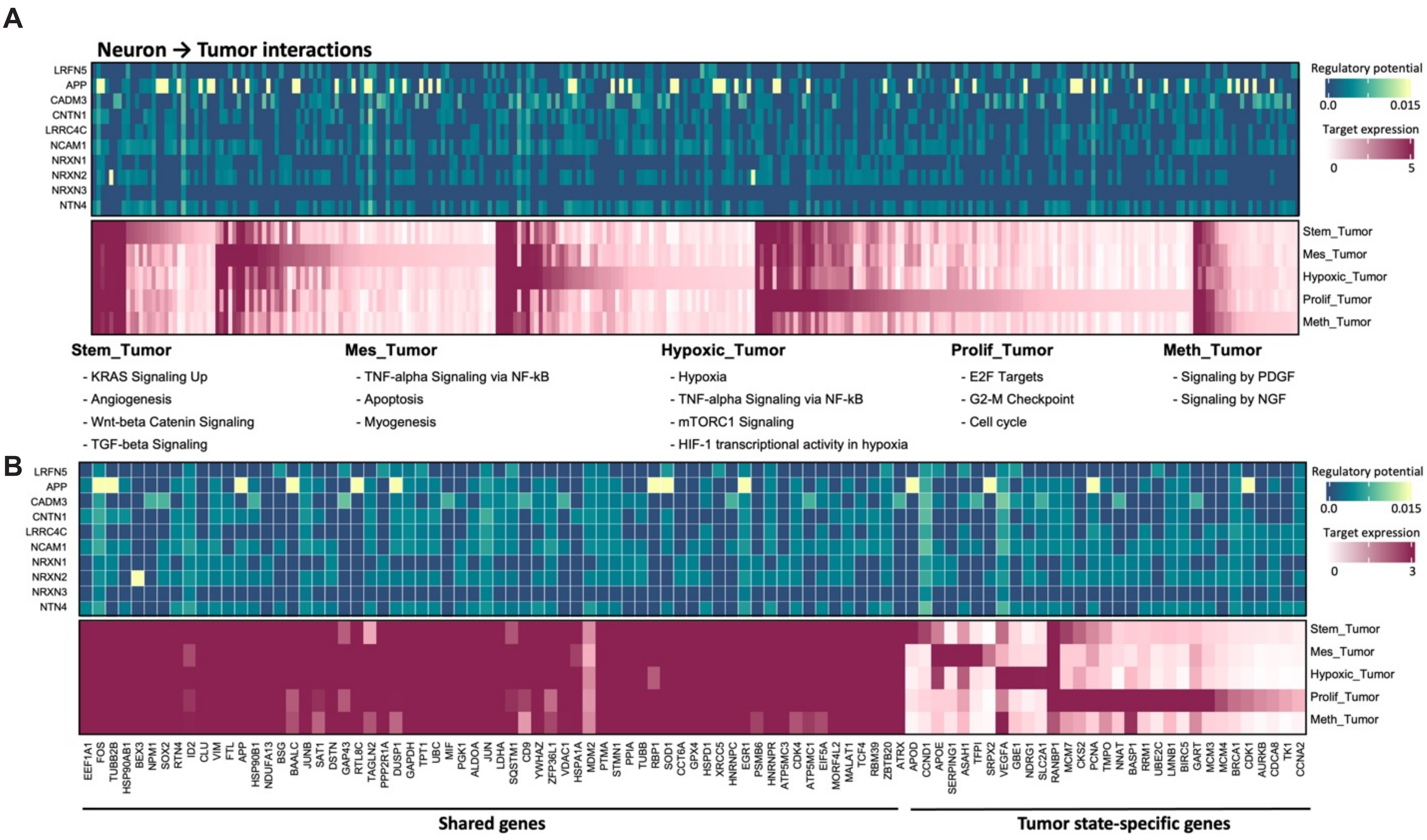

### Fig. S26

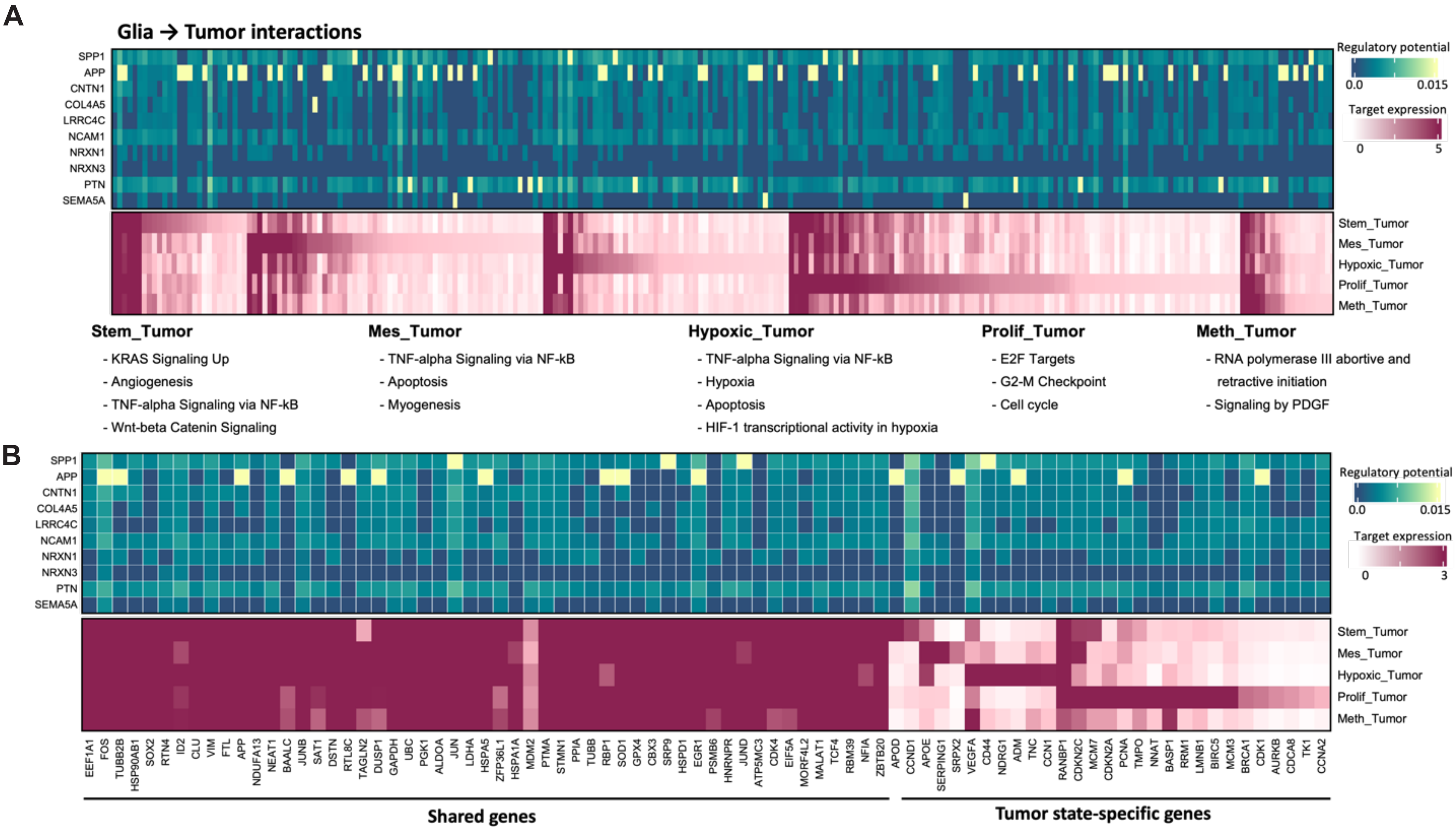

### Fig. S28

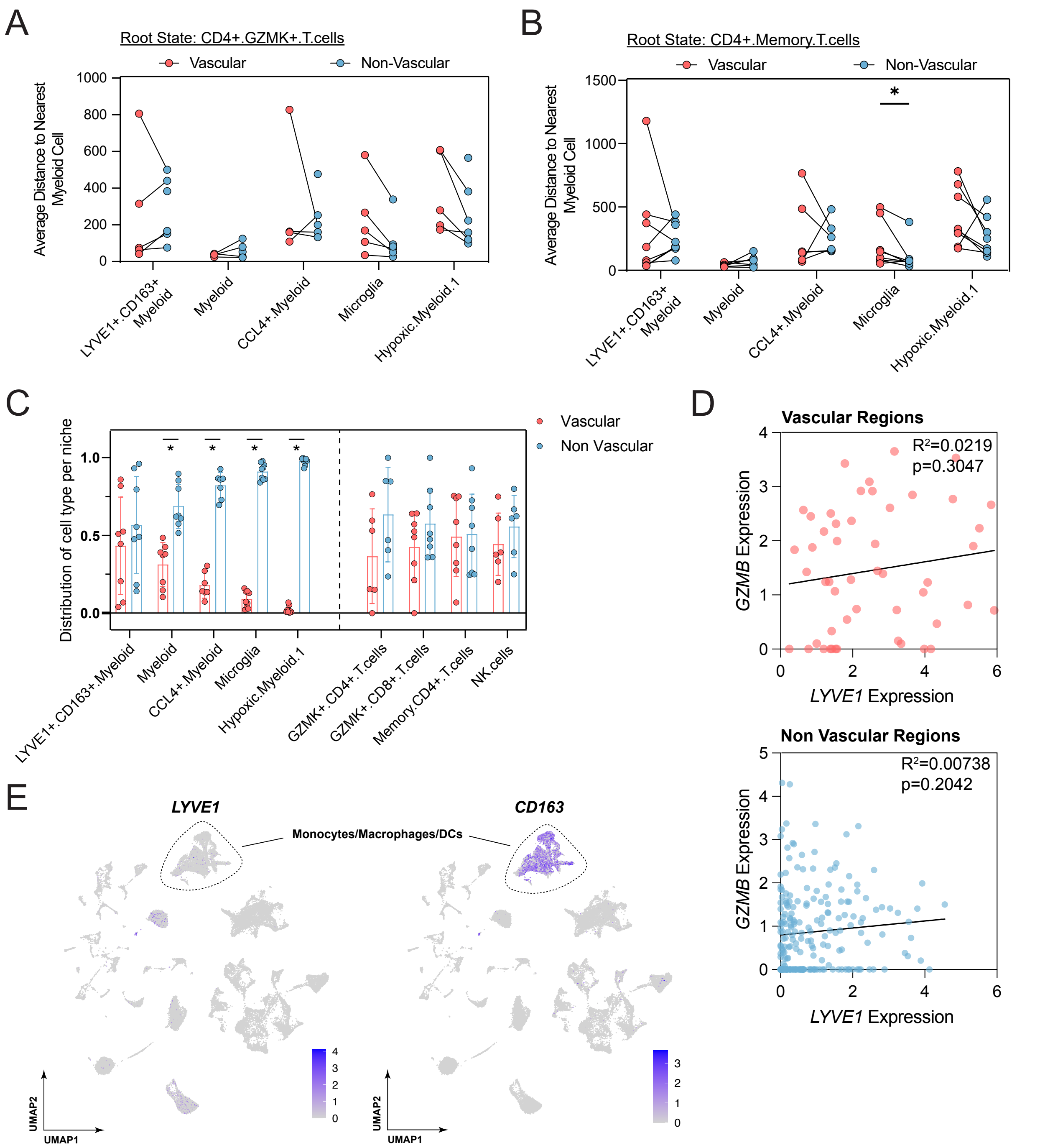
