## Supplementary material for "Mapping the spatial architecture of glioblastoma from core to edge delineates niche-specific tumor cell states and intercellular interactions": Fig. S4

### Visium FFPE samples (and adjacent slices analyzed using Xenium, where applicable)

**ST073021.02 Edge**
**Visium**

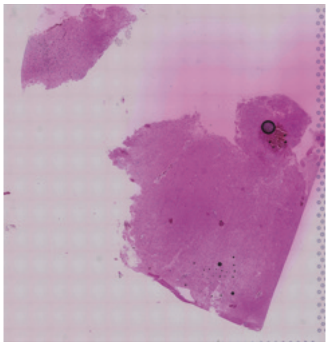

**ST073021.02 Edge**
**Xenium**

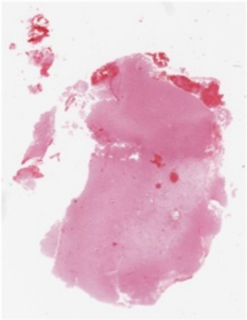

**ST073021.04 Core**
**Visium**

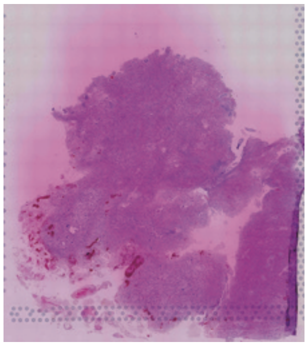

**ST073021.04 Core**
**Xenium**

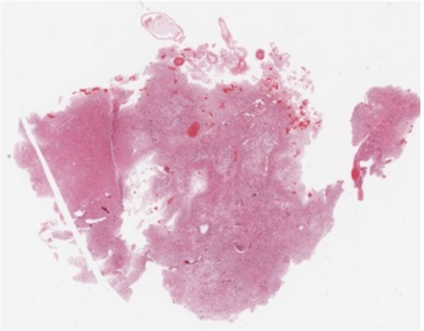

**WU1220.01 Transition**
**Visium**

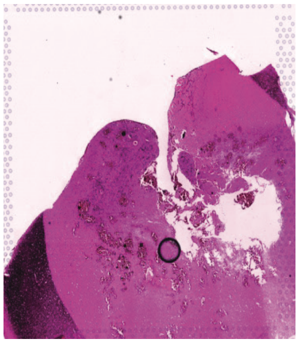

**WU1220.02 Edge**
**Visium**

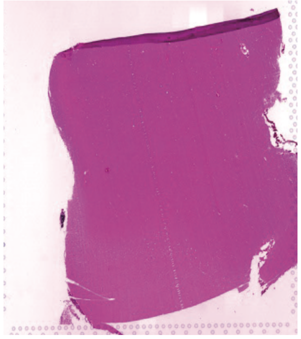

**WU1220.02 Edge**
**Xenium**

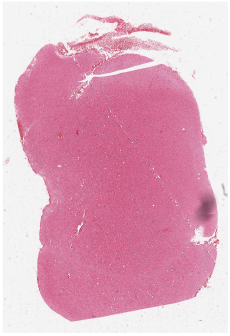

**WU1220.03 Core**
**Visium**

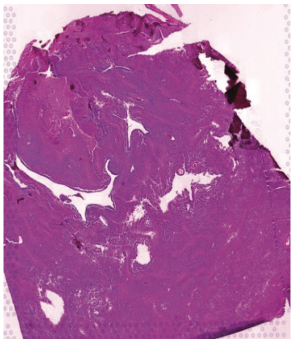

**WU1220.03 Core**
**Xenium**

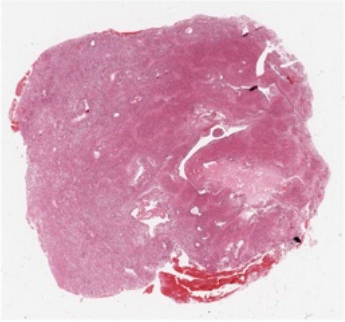

**WU1221 Edge**
**Visium**

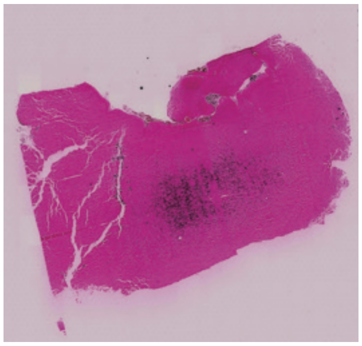

**WU1221 Edge**
**Xenium**

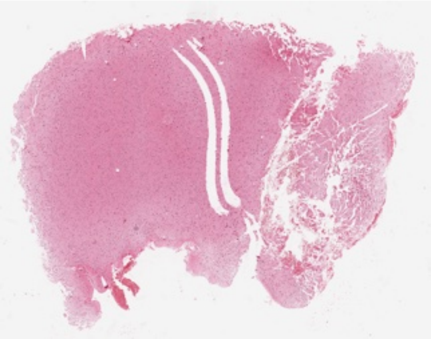

**WU1221 Core**
**Visium**

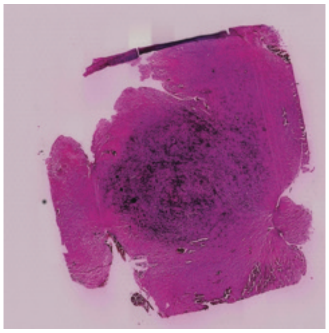

**WU1221 Core**
**Xenium**

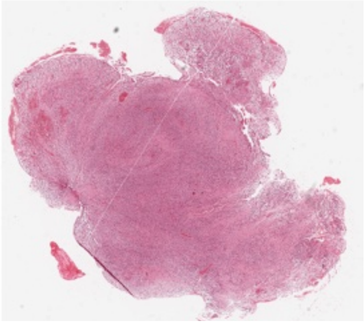

**WU1227 Edge**
**Visium**

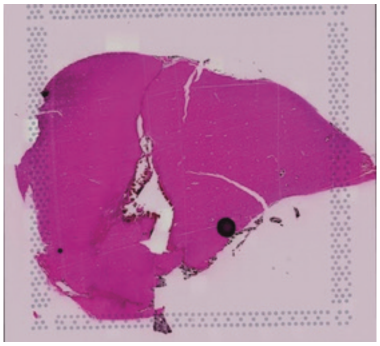

**WU1227 Core**
**Visium**

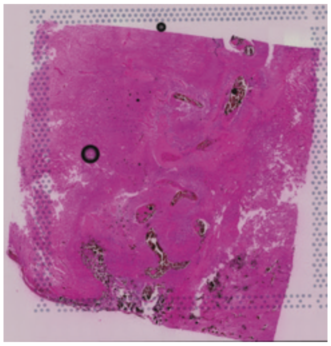
