## Supplementary material for "Mapping the spatial architecture of glioblastoma from core to edge delineates niche-specific tumor cell states and intercellular interactions": Fig. S10

| Spatial Tumor State | Processes | Overlapping Genes | Neftel overlap | Nomura overlap | Greenwald overlap |
|---|---|---|---|---|---|
| **Meth_Tumor** | Methylation, neuronal projections, acetylcholine response | DPYSL3, NFIX, ANKRD12, APC, BPTF, CEP170, PITPNC1, PLCB1, | NPC2 (3) | MP_4_AC (1), MP_7_NPC (1) | NPC (1), chromatin (4) |
| **Prolif_Tumor** | Cell cycle | CENPF, HES6, HIST1H4C, NFIB, SMC4, STMN1, TMPO, ANP32B, CKAP2, DHFR, DUT, EZH2, GMNN, JPT1, KPNA2, LMNB1, MCM3, MCM7, PCNA, RRM1, TUBA1B, | G1 (7), G2 (5), NPC1 (2), NPC2 (2) | MP_14_NRGN (1), MP_1_RP (2), MP_3_CC (5) | AC (1), NPC (2), OPC (2), chromatin (2) |
| **Mes_Tumor** | Mesenchymal programs, immune response | APOE, CHI3L1, CLU, CST3, MT2A, MT3, SPP1, VIM, AGT, ATP1B2, EFEMP1, FABP7, HOPX, IGFBP7, LGALS3, RAMP1, RCAN1, S100A16, SERPING1, APLP2, B2M, CHPF, CTSB, GPC1, IFITM3, ITM2C, MT1E, PDPN, PLTP, PON2, PSAP, SAT1, SERPINA3, SPARC, TIMP1, TNFRSF1A, ZYX, | AC (12), MES1 (17), MES2 (2) | MP_11_MIC (1), MP_14_NRGN (2), MP_1_RP (8), MP_6_MES (1) | AC (19), MES (10), hypoxia (4) |
| **Hypoxic_Tumor** | Hypoxia response | ACTN1, ADM, AKAP12, CD44, PLOD2, SLC2A3, CA9, CLIC4, EGLN3, ERO1A, GBE1, P4HA1, SLC2A1, SOD2, TNC, UFM1, VEGFA, ARL4C, CADPS, CAMK2N1, CAV1, FAM20C, GADD45A, PLP2, PPP1R15A, PPP2CB, SEC61A1, SLC16A1, TMEM158, TNFRSF12A, | MES1 (5), MES2 (11) | MP_10_Stress1 (3), MP_14_NRGN (1), MP_15_Stress2 (4), MP_4_AC (3), MP_5_Hypoxia (9), MP_6_MES (6) | AC (3), MES (4), hypoxia (10), metabolism (1) |
| **Stem_Tumor** | Neural development | CADM2, DBI, ETV1, SOX4, BCAN, CHD7, DLL3, FXYD6, GLCCI1, HIP1, LRRN1, MARCKS, MARCKSL1, NDRG2, NKAIN4, PGRMC1, PTPRZ1, SCD5, SCRG1, SIRT2, SOX11, TCF12, THY1, TTYH1, VCAN, ABAT, ASCL1, BEX1, C1orf61, COL9A3, CSPG5, DBN1, ELAVL3, FAM181B, GRIA2, LIMA1, MAP2, MMP16, MTSS1, NRXN1, OLIG2, PFN2, PHYHIPL, RTKN, SEMA5A, SHD, SLC44A1, SOX8, TSC22D4, | AC (5), NPC1 (19), NPC2 (3), OPC (16) | MP_11_MIC (1), MP_14_NRGN (2), MP_1_RP (2), MP_2_OPC (11), MP_7_NPC (4), MP_8_GPC (2) | AC (13), OPC (19), chromatin (3), metabolism (1) |

Neftel
Neftel+Greenwald
Nomura
Nomura+Greenwald
Greenwald
Neftel+Nomura
Neftel+Nomura+Greenwald
