## Supplementary material for "Mapping the spatial architecture of glioblastoma from core to edge delineates niche-specific tumor cell states and intercellular interactions": Fig. S27

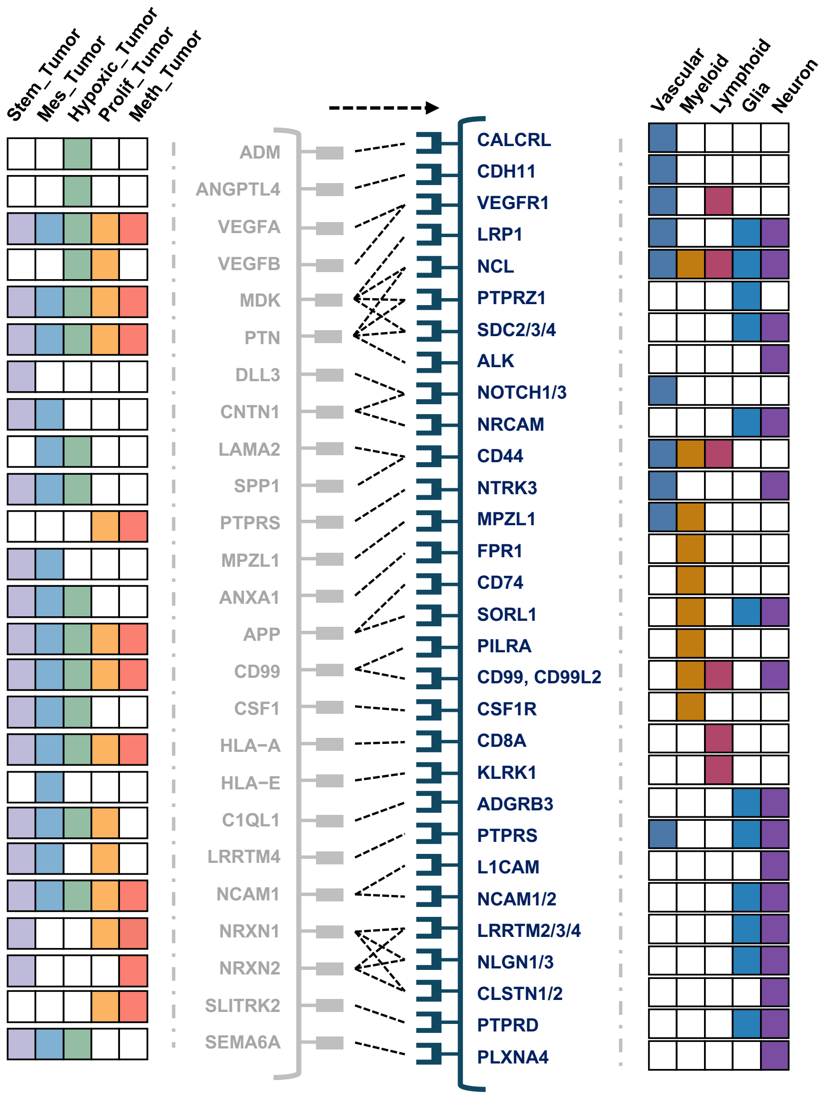


**Vascular**

- HIF-2-alpha transcription factor network
- KRAS Signaling Up
- Pathways in cancer

**Myeloid**

- TNF-alpha Signaling via NF-kB
- Immune System
- Innate Immune System

**Lymphoid**

- TNF-alpha Signaling via NF-kB
- Interferon Response
- Cytokine Signaling in Immune System
- Immune System

**Glia**

- KRAS Signaling Up
- Notch Signaling

**Neuron**

- MAPK signaling path
- Axon guidance
- Acetylcholine Neurotransmitter Release Cycle
