## Supplementary Figure and Table Legends for "Mapping the spatial architecture of glioblastoma from core to edge delineates niche-specific tumor cell states and intercellular interactions"

**Fig. S1. Visium analysis workflow.**

**(A)** Single-cell RNA-sequencing data was used as a reference, and cells were mapped onto the corresponding Visium slides using CellTrek (with interpolation) (1). Spatially variable tumor cell clusters were identified for each patient using CellTrek. A binary matrix representing these tumor sub-clusters across patients was constructed (2,3). Consensus partitioning and ranking methods were then applied to this matrix to define spatially variable tumor cell states (4) (see Methods). **(B)** Visium data was integrated across patients using RPCA-based batch correction and graph-based clustering at a resolution of 0.9. CellTrek (without interpolation) was then used to assign cell types to each Visium spot (1). Cell type frequencies and cell type enrichments were calculated for each cluster, as shown in Fig. 3A (2). Subsequently, spatial adjacency analysis of cellular niches (3) and ligand–receptor interaction analysis (4) were performed on the Visium spots, and cell type frequencies were visualized.

**Fig. S2: Representative pre-operative MRIs of edge and core (for patient B189)**

**Fig. S3: H&E images for OCT samples analyzed using Visium.** H&E Visium slides from OCT-preserved specimens. In some cases, distinct core and edge regions were available, as illustrated; in others, transition regions—annotated by the surgeons or pathologists—were obtained, as shown.

**Fig. S4: H&E images for FFPE samples analyzed using Visium and same-block samples analyzed using Xenium.**

**Fig. S5: Pathologist annotations of the Visium slides.** Annotations by a neuropathologist (S.D.) for the H&E slides of ST073021.04 (Core), ST073021.02 (Edge), WU1220.01 (Edge), WU1220.02 (Edge), and WU1227 (Core), with Visium spots superimposed, are presented. Key histopathological features characteristic of GBM such as microvascular proliferation (MVP), palisading necrosis, myelin-rich regions, and tumor-rich regions are indicated.

**Fig. S6. CNV Spot classification.**
Spots were classified into Low/Medium/High malignancy based on the frequency and magnitude of copy number alterations. High malignancy spots (i.e. Spots with high CNV scores) usually reside in the Core-rich domains/Niches whereas Low malignancy spots (i.e. Spots with low CNV scores) are in Edge-rich domains as shown in the spatial plots. Spot malignancy proportions align well with cell type proportions across clusters, with regions enriched in tumor cells exhibiting correspondingly higher CNV scores as shown in the bar plots.

**Fig. S7. Batch-corrected RPCA clustering across patients at resolution 0.1.**
**(A)** Visium spots colored by cluster assignment across patients and samples. **(B)** Bar plots depicting the distribution of patients and samples within each cluster. **(C)** Gene Ontology (GO) biological processes enriched among the top 200 differentially expressed genes for each cluster (adjusted $p \leq 0.05$, ranked by average $\log_2$ fold change). The analysis indicates that low-resolution batch integration effectively identifies biologically distinct clusters, including regions associated with microvascular proliferation (MVP), as well as glia-rich and neuron-rich areas.

**Fig. S8. Batch-corrected RPCA clustering across patients at resolution 0.9.**
**(A)** Visium spots colored by cluster assignment at resolution 0.9 across patients. **(B)** Bar plots showing the distribution of patients and samples within each cluster. **(C)** Gene Ontology (GO) (biological processes) enriched among the top 200 differentially expressed genes for each cluster (adjusted $p \leq 0.05$, ranked by average log fold change). Higher-resolution (0.9) batch-integration

clustering captured more extensive transcriptional variation, revealing both shared and sample-specific clusters, as well as features specifically enriched in either the tumor core or edge.

**Fig. S9. Spatial tumor states compared with published single-cell-derived tumor cell states. (A)** The heatmap displays adjusted *p*-values reflecting the significance of overlap between tumor states from five publications and our spatially defined tumor states, highlighting both shared and distinct features of these definitions. **(B)** Heatmaps indicate tumor state classifications of tumor cells from scRNA-seq data using Neftel (top) and spatially-derived (bottom) definitions. The spatial classifications reveal more distinct transcriptional differences, suggesting improved stratification of tumor cells using spatially-informed cell state definitions.

**Fig. S10. Spatial tumor states compared with published single-cell-derived tumor cell states, part 2: marker gene comparison.**

**Fig. S11: Analysis of Cohort 1 and Cohort 2 Xenium samples. (A)** UMAP visualization of Cohort 2 samples alone. **(B)** Correlation analysis of gene expression of respective cell type annotations in Cohort 1 and Cohort 2. **(C)** Dot plot of expression of select genes used for cell type annotation. **(D)** Bar plot of frequency of each cell type per sample in both Cohorts 1 and 2. **(E)** Heatmap of niche occurrence within every sample with the total occurrence of each niche across all samples summarized at the top.

**Fig. S12: Xenium images colored by cell type. (A)** Primary cohort. **(B)** Secondary cohort. **(C)** Tertiary cohort (edge samples). **(D)** Tertiary cohort (Core samples)

**Fig. S13. Niche definitions and comparisons. (A)** Hierarchical clustering of BANKSY clusters from both cohorts (primary and secondary) based upon normalized values of cell type frequency. **(B)** Heatmap of Spearman correlation coefficients between Visium niches and Lin, et al, domains. **(C)** Heatmap of Spearman correlation coefficients between Visium niches and Greenwald layers. **(D)** Xenium niches using Xenium-derived cell types instead of harmonized cell types. **(E)** Heatmap of Pearson correlation coefficients between Visium and Xenium niches.

**Fig. S14: Cell type and spatial enrichment of niches. (A)** Pearson correlation of cell type pairs across Visium niches. **(B)** Spatial proximity of Xenium niches. **(C)** Edge/Core enrichment of tumor cell states in Xenium data (Cohort 3: tissue blocks matched to Visium tissue blocks). **(D)** Edge/Core enrichment of tumor cell states in paired edge/core scRNA-seq samples.

**Fig. S15: Spatial adjacency between transcriptional clusters.** As shown in Fig. 3G, multiple clusters display spatial co-localization. Spatial adjacency between the Hypoxia cluster (C3) and the microvascular proliferation (MVP) cluster (C6) is demonstrated on the Visium spots across multiple samples, consistent with the adjacency map in Fig. 3G. Additional examples of spatially adjacent clusters include MVP (C6) with S3 Mesenchymal (C17), MVP (C6) with S3 Mesenchymal (C9), and MVP (C6) with S5 Stem-like (C7), emphasizing the central role of microvascular proliferation in GBM tumor architecture. In edge-enriched regions, spatial adjacency between Oligo-rich (C0) and Neuron-rich (C13) clusters is also observed.

**Fig. S16: Integrated Ligand-Receptor Interaction Analysis.** (A) Dot plot of high-confidence ligand-receptor (LR) interactions from individual GBM tumor cell states (senders, columns) to all cell types in the tumor microenvironment (receivers, rows). LR pairs (n=151) are grouped by biological function (“Function” color bar) and spatial domain enrichment (Domain). Sender columns represent eight tumor cell states. Receiver dot color reflects the platform evidence supporting the interaction: blue tones indicate Xenium-confirmed interactions; red/pink tones indicate Visium-only or Visium+GBMap confirmation. Receiver columns include non-tumor microenvironmental cell types (Vascular, Myeloid, Lymphoid, Glia, Neuron) and a grouped Tumor receiver category. Bold red LR pair labels denote clinically targetable interactions based on DGIdb (approved antineoplastic agents). Interactions meet the following criteria: ≥2 independent evidence sources, ≥1 spatial platform (Visium or Xenium), and ≥1 single-cell resolution platform (scRef, GBMap, or Xenium). (B) Number of unique LR interaction between indicated cell types. (C) Analogous to panel A, but for intratumoral interactions.

**Fig. S17: Ligand-Receptor Analysis of Xenium Samples.** Heatmaps of the (A) unique and (B) total number of ligand receptor interactions between general cell types.

**Fig. S18: Cell chat analysis of Xenium samples. (A)** Bar plot showing the number of ligand-receptor interactions with the specified cell type either as the sender or receiver. **(B)** Interaction network showing ligand receptor interactions with tumor cells as senders and all other cell types as receivers. The density and size of arrows were proportional to the number of interactions detected as significant for a given sender-receiver pair. **(C)** UMAP visualization of scRNA-seq data including both GBM samples and normal brain cell types. **(D)** UMAP visualization of both *ADM* and *CALCRL* gene expression within scRNA-seq data set. **(E)** Interaction network of niche sender-receiver pairs of the *ADM:CALCRL* ligand-receptor pair. The color and density of arrows were proportional to the number of samples the *ADM:CALCRL* interaction was detected as significant for a given sender-receiver pair.

**Fig. S19: Ligand-receptor interactions enriched in spatial domains in Visium data.** Diffusion-based LR interactions **(A)** within and **(B)** between larger spatial domains (Edge, Core, Interface, and Non-Specific). Dot color indicates the adjusted p-value (i.e. -log10(p.adj+1), hypergeometric test) for LR's that are significant in at least one domain. Dot size indicates enrichment score, and the black dot border is a binary indicator of statistical significance. The heatmap (right) indicates cell type composition for the spots in each domain that exhibit these LRs. **(C)** Summary of key interactions.

**Fig. S20: Representative RNAscope Images. (A)** Representative image of panel 1: CD45 (green), CD44 (red), CALCRL (magenta), ADM (yellow), PECAM1 (cyan), SOX2 (white), and DAPI (blue). **(B)** Representative image of panel 2: PDGFRB (magenta), HES1 (yellow), JAG1 antibody (green), NOTCH3 antibody (red), PECAM1 (cyan), SOX2 (white), DAPI (blue). **(C)** Representative image of panel 3: CD45 (green), CD44 (red), NCL (magenta), PTN (yellow), PECAM1 (cyan), SOX2 (white), DAPI (blue).

**Fig. S21. Interaction heatmap between non-tumor and tumor cells.**
Heatmap showing interactions between non-tumor cells, including vascular, myeloid, lymphoid, glial, and neuronal cells, and tumor cells. Colors indicate the techniques combination supporting each interaction. Blue indicates interactions detected between Xenium and single-cell transcriptomic techniques (scRef and GBmap), or detected by both spatial platforms (Xenium and Visium). Red indicates interactions detected between Visium and single-cell transcriptomic (scRef and GBmap). Because of the lower resolution of Visium, Visium-based interactions are represented at the domain level.

**Fig. S22. Vascular-tumor interaction and potential targets in tumor cells.**
Heatmaps illustrating the predicted regulatory potential of vascular-derived ligands and the expression of their predicted target genes in tumor cells. **(A)** Predicted target genes expressed in more than 30% of cells within each tumor subtype, together with the biological pathways enriched among ligand-regulated targets in each subtype. **(B)** Ligand-regulated target genes showing the

highest shared expression across tumor subtypes, together with subtype-specific target genes uniquely enriched in individual tumor subtypes.

**Fig. S23. Myeloid-tumor interaction and potential targets in tumor cells.**
Heatmaps illustrating the predicted regulatory potential of myeloid-derived ligands and the expression of their predicted target genes in tumor cells. **(A)** Predicted target genes expressed in more than 30% of cells within each tumor subtype, together with the biological pathways enriched among ligand-regulated targets in each subtype. **(B)** Ligand-regulated target genes showing the highest shared expression across tumor subtypes, together with subtype-specific target genes uniquely enriched in individual tumor subtypes.

**Fig. S24. Lymphocyte-tumor interaction and potential targets in tumor cells.**
Heatmaps illustrating the predicted regulatory potential of lymphocyte-derived ligands and the expression of their predicted target genes in tumor cells. **(A)** Predicted target genes expressed in more than 30% of cells within each tumor subtype, together with the biological pathways enriched among ligand-regulated targets in each subtype. **(B)** Ligand-regulated target genes showing the highest shared expression across tumor subtypes, together with subtype-specific target genes uniquely enriched in individual tumor subtypes.

**Fig. S25. Neuron-tumor interaction and potential targets in tumor cells.**
Heatmaps illustrating the predicted regulatory potential of neuron-derived ligands and the expression of their predicted target genes in tumor cells. **(A)** Predicted target genes expressed in more than 30% of cells within each tumor subtype, together with the biological pathways enriched among ligand-regulated targets in each subtype. **(B)** Ligand-regulated target genes showing the highest shared expression across tumor subtypes, together with subtype-specific target genes uniquely enriched in individual tumor subtypes.

**Fig. S26. Glia-tumor interaction and potential targets in tumor cells.**
Heatmaps illustrating the predicted regulatory potential of glia-derived ligands and the expression of their predicted target genes in tumor cells. **(A)** Predicted target genes expressed in more than 30% of cells within each tumor subtype, together with the biological pathways enriched among ligand-regulated targets in each subtype. **(B)** Ligand-regulated target genes showing the highest shared expression across tumor subtypes, together with subtype-specific target genes uniquely enriched in individual tumor subtypes.

**Fig. S27. Overall interaction map from tumor cells to non-tumor cells.**
Heatmaps illustrating interactions from multiple tumor subtype to non-tumor cells including vascular, myeloid, lymphoid, and neuron (top), and biological pathways enriched among the predicted target genes associated with the corresponding interactions (bottom).

**Fig. S28: Analysis of T cell and myeloid cell co-localization within Xenium samples. (A)** Distance to the nearest target myeloid cell of each *CD4+.GZMK+* T cell separated by localization within a vascular or non-vascular niche averaged on a per sample basis. Each dot represents an individual sample. **(B)** Distance to the nearest target myeloid cell of each *CD4+.Memory* T cell separated by localization within a vascular or non-vascular niche averaged on a per sample basis. Each dot represents an individual sample. **(C)** Frequency of lymphoid and myeloid cell types based upon vascular or non-vascular localization calculated on a per sample basis. **(D)** Correlation of *GZMB* and *LYVE1* expression based upon localization in vascular or non-vascular niches. Data derived from IVY-Gap. **(E)** UMAP visualization of *LYVE1* and *CD163* expression within the scRNA-seq data set. Significance in **(A-C)** was calculated using the Wilcoxon matched-pairs signed rank test.

**Table S1:** Clinical details of samples analyzed using Visium and Xenium

**Table S2:** QC metrics for Visium data

**Table S3:** Differentially expressed genes in each spot cluster, at cluster resolution 0.1 (Visium)

**Table S4:** Differentially expressed genes in each spot cluster, at cluster resolution 0.9 (Visium)

**Table S5:** Functional enrichment (with respect to Gene Ontology Biological Processes) of DEGs in each spot cluster, at cluster resolution 0.9

**Table S6:** Gene sets of tumor states (S1-S5) identified after Consensus Clustering

**Table S7:** Xenium Probe sets

**Table S8:** Xenium QC metrics

**Table S9:** Gene markers for cell type identification in Xenium data

**Table S10:** DEGs for each cell type in Xenium data

**Table S11:** Sample-specific Xenium niches obtained using BANKSY

**Table S12:** Enrichment of each cell type in each Visium niche and each Xenium niche

**Table S13:** Xenium inter-niche distances, used to calculate Xenium niche distance graph

**Table S14:** TME-to-Tumor LR interactions with platform-specific evidence and Visium spatial domain enrichment

**Table S15:** TME-to-Tumor LR interactions with platform-specific evidence and Visium spatial domain enrichment

**Table S16:** Predicted target genes downstream of LR interactions (organized by cell type pair)

**Table S17:** Antibodies and methodological details for multiplexed immunofluorescence

**Supplementary Data (Contact_dependent_LR)**: Within-spot Hypergeometric test enrichment results of LR (per cluster) for Contact-dependent ligand-receptor pairs only.

**Supplementary Data (diffusion_LR_enrichment_byDomain)**: Hypergeometric test results of LR enrichment for diffusion-based ligand-receptor pairs.

**Supplementary Data (diffusion_LR_enrichment_byCluster)**: Hypergeometric test results of LR enrichment for diffusion-based ligand-receptor pairs.

**Supplementary Data (scRNA-seq reference LR predictions CellChat V2)**: Ligand receptor interactions that are significant and have a score greater than overall mean communication probability are shown for each celltype.

**Supplementary Data (DGIDB_visium)**: Ligand receptor candidates from Visium and their associated drug interactions from DGIDB database.

**Supplementary Data (DGIDB_xenium)**: Ligand receptor candidates from Xenium and their associated drug interactions from DGIDB database.
